# The DNA legacy of Thomas Jefferson

**DOI:** 10.64898/2026.09.18.752809

**Authors:** Richard E. Green, Samuel H. Vohr, Joshua D. Kapp, Jane E. Ailes, Samuel Sacco, Remy Nguyen, James Cahill, Peter D. Heintzman, Joanne Flores, Logan Kistler, Courtney A. Hofman, Robert C. Fleischer, Beth Shapiro, Richard Kurin

## Abstract

Allegations that Thomas Jefferson fathered the children of the enslaved Sally Hemings emerged in the context of U.S. presidential politics in the late 1700s and early 1800s and entailed strongly contested denials and affirmations by family members, biographers, and historians ever since. Previous analysis of Y chromosome markers in contemporary members of the Jefferson and Hemings patrilines was consistent with the paternity hypothesis but rejected by those who pointed out that, absent Jefferson’s DNA and its direct comparison to the DNA of possible descendants, other Jefferson males could have fathered Sally Hemings’ children. In this study, we developed an innovative technique to recover DNA from multiple rootless hair samples in a museum collection whose provenance, mitochondrial genomes, Y chromosome markers, and identity to each other indicate that these hairs and the DNA from them are from Thomas Jefferson. Using these DNA data, we find clear evidence of Thomas Jefferson ancestry in descendants of both his wife Martha Wayles Jefferson and Sally Hemings. We measured the amount of identical-by-descent DNA shared between Thomas Jefferson and his known and putative descendants. We find that the amount of shared Jefferson DNA is more likely under the model of direct Thomas Jefferson paternity than other proposed scenarios.

## Main Text

During his lifetime and since, the question of Thomas Jefferson’s paternity of Sally Hemings’ children has been a controversial and contentious issue, bearing on the paradoxes of American liberty and equality. Jefferson was the principal author of the Declaration of Independence in 1776 with its famous clarion statement that “all men are created equal, that they are endowed by their Creator with certain unalienable Rights, that among these are Life, Liberty and the pursuit of Happiness.” Yet over his lifetime Jefferson kept more than 600 enslaved people at his Monticello estate and other properties. Allegations in the run-up to his election as U.S. president and through his first term suggested he fathered several children with the enslaved Sally Hemings, some thirty years his junior, magnifying what some observers noted as a basic hypocrisy. How could Jefferson articulate for people of a new nation the right to equality and liberty, yet consign his own children to slavery?

While Jefferson himself never directly addressed the allegation, his family steadfastly denied it, pointing over the decades to other family members like his nephews Peter and Samuel Carr as possible fathers. For well more than a century, historians and biographers also discounted the possibility of Jefferson’s paternity based on his character and lack of any eyewitness accounts or other convincing evidence. But several researchers and scholars, like Pearl Graham in the 1960s (1), Fawn Brodie in the 1970s (2), and most thoroughly Annette Gordon-Reed in the 1990s (3), pointed to oral historical accounts by Hemings family members and associates, the concurrence of Jefferson’s residencies at Monticello with Sally Hemings’ pregnancies, and several other factors to argue there was adequate evidence to accept his paternity.

The tide turned among historians and in popular opinion in 1998 with DNA evidence. Eugene Foster and colleagues analyzed Y chromosome markers of patriline relatives of Thomas Jefferson (although not Thomas Jefferson himself), a patriline descendant of Eston Hemings (a potential son of Sally Hemings and Thomas Jefferson), patriline descendants of the Carr family (proposed alternative fathers of Sally Hemings’ offspring), and patriline descendants of Thomas Woodson (the first purported son of Sally Hemings and Thomas Jefferson) (4). Foster could not find patriline descendants of Sally Hemings’s two other sons, Madison and Beverly. The analysis found that the Jefferson and Eston Hemings lines matched at seven bi-allelic SNP markers, eleven STR markers, and a minisatellite marker. The Carr and Woodson patriline genotypes did not match the Jefferson patriline genotypes. This observation was offered in support of the conclusion that a Jefferson and not a Carr (Peter or Samuel) fathered Eston Hemings with Sally Hemings. Further, a Jefferson was not the father of Thomas Woodson—a finding questioned by his descendants but confirmed through subsequent sampling and analysis (5).

A subsequent genetic analysis of the Foster-study samples further delineated the Jefferson Y chromosome haplotype, identifying a rare A to C derived allele at the M70 bi-allelic SNP locus (6). This derived allele was also found to be shared with the Eston Hemings patriline descendant.

The strength of this conclusion has been hotly debated. The genetic evidence is an analysis of a single, non-recombining marker: the Y chromosome for one of Sally Hemings’s sons. This finding combined with historical evidence led many to believe that Thomas Jefferson was the father of at least one and likely all of Sally Hemings’ children. Others disputed that conclusion. They argued that the evidence did not rule out paternity of Eston Hemings by some other Jefferson patriline relative like Thomas Jefferson’s brother Randolph (7) or even an enslaved descendant of such a kinsman (8). And Foster’s study did not address the paternity of Sally Hemings’s other sons or her daughter Harriet.

After reviewing Foster’s DNA study and the historical evidence, the Thomas Jefferson Foundation, which owns and operates Monticello, concluded in 2000 that “although paternity cannot be established with absolute certainty, our evaluation of the best evidence available suggests the strong likelihood that Thomas Jefferson and Sally Hemings had a relationship over time that led to the birth of one, and perhaps all, of the known children of Sally Hemings (9).” Another organization, the Thomas Jefferson Heritage Society, formed a scholars commission that strongly challenged that conclusion, finding that Foster’s study did not prove Thomas Jefferson’s paternity, but rather placed “him in a group of approximately twenty-five known Virginia men believed to carry the Jefferson family Y chromosome (10).” In the aftermath, even as Jefferson and Hemings descendants gathered annually at Monticello to discuss the issue, oppositional positions affirming and denying Jefferson’s paternity hardened. In 2002, the Monticello Association, which is distinct from the Thomas Jefferson Foundation and comprised of known Jefferson descendants who own and operate the family graveyard on the grounds, voted overwhelmingly not to admit the Hemings descendants as members, thus denying them the right to be buried there (11). The lead protagonist for the Monticello Association at the time noted that the standard for proving paternity in the Commonwealth of Virginia was “clear and convincing evidence,” and the only way of meeting that standard was to obtain Thomas Jefferson’s DNA and compare it directly to Hemings descendants to see if there was an appropriate and definitive match (12).

We collected DNA from Thomas Jefferson by extracting it from fragments of rootless hair of known provenance in museum collections. We also collected DNA samples from an array of living descendants of the Jefferson and Hemings lines for comparison. We identify regions that are identical by descent (IBD) from Thomas Jefferson and living descendants of both the Jefferson and the Hemings line. Statistical analyses of the amount of DNA shared between Thomas Jefferson and the Hemings’ descendants provide strong evidence that Thomas Jefferson himself, not a close relative, fathered multiple children with Sally Hemings.

### Identification of *bona fide* Thomas Jefferson hair

During this project, we extracted, sequenced and analyzed DNA from 12 different samples of historical hair all of which lacked follicles. These hairs included those recovered from the famed Jefferson Bible at the Smithsonian’s National Museum of American History, a supposed presidential reliquary, also at the Smithsonian, and several mementos and packets purportedly containing locks of Jefferson’s hair in the collections of the Thomas Jefferson Foundation at Monticello (Table S1) (13). A number of these were determined not to be Thomas Jefferson’s hair; others were indeterminable. We focus here on the hair samples that we conclude are from Thomas Jefferson.

We used several criteria for determining if a hair derives from Thomas Jefferson. First, hair should derive from a genetic male. To determine this, we used the ratio of reads that map to the X chromosome versus those that map to chromosome 10, a similarly sized autosome (Fig 1A). This strategy was previously shown to be reliable for determining genetic sex using DNA from hair (14). While some hairs had too little DNA to accurately determine this ratio and others clearly indicated that the individual was a genetic female, several samples indicated the hair was from a male individual.

**Figure 1.**
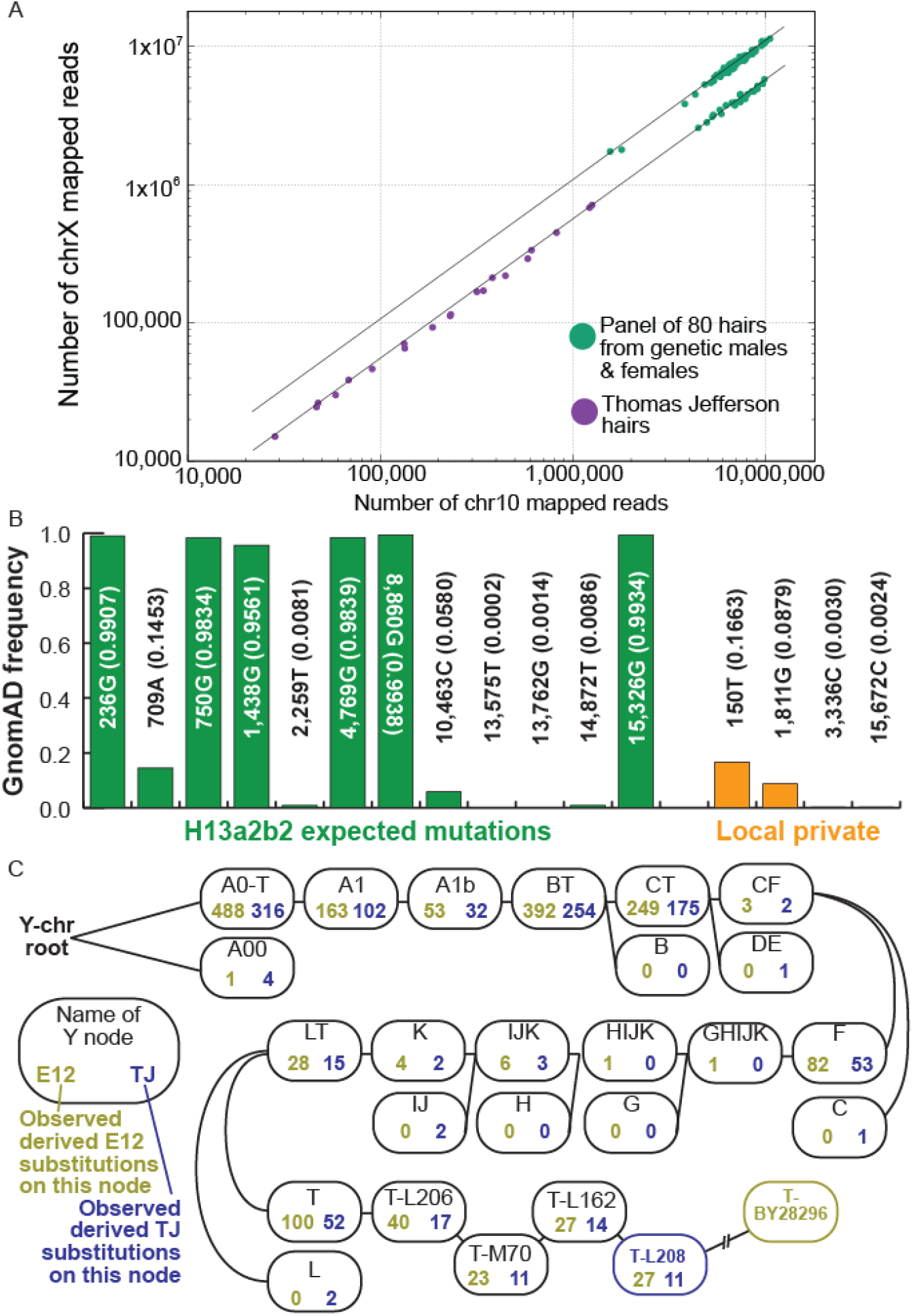
Authenticity criteria for Thomas Jefferson DNA. (A) Rate of fragments that map to chrX and chr10 from Thomas Jefferson hair libraries (purple) and a panel of 80 individual hairs, both male and female. (B) Frequencies of non-reference alleles present in mitochondrial genomes of both Thomas Jefferson and L7. (C) Y-chromosome tree of E12 and Thomas Jefferson. Note the higher coverage of E12 allows a more precise haplotype call.

The second authenticity criterion was the mitochondrial genome. Thomas Jefferson’s mitochondrial genome was not known beforehand. Therefore, we included a living direct matriline descendant of Thomas Jefferson’s sister, Lucy Jefferson, in the sampling strategy (L7). If the genealogy is correct – and because false maternity is extremely rare – *bona fide* Thomas Jefferson DNA should be a mitochondrial match of this individual. From saliva DNA, we assembled a complete mitochondrial genome from individual L7 from two independent libraries. Haplogrep v3.2.1 (15) analysis indicated it is haplogroup H13a2b2, identifying 12 haplogroup-informative mutations, 3 private hot-spot mutations, and 4 local private mutations. DNA from Thomas Jefferson should share these (Fig. 1B, SOI).

Third, DNA covering Y chromosome markers should be consistent with the Jefferson patriline markers described in the Foster (4) and King (6) studies. Further, *bona fide* Thomas Jefferson DNA may be expected to share any further substitutions seen in analysis of DNA from the patriline descendant of Eston Hemings, E12 (Fig. 1C).

Finally, DNA from any hair from Thomas Jefferson should be consistent with deriving from the same person in any pairwise comparison to another hair deriving from Thomas Jefferson. For pairwise comparisons of hair, we used the tilde program (16) which compares two DNA datasets and generates a likelihood ratio of the data deriving from the same or different individuals.

Evaluation of these four criteria across sequencing libraries revealed that hair samples from Monticello collection packets catalog numbers 1956-31, 1968-67-7, and 1968-67-8 are from Thomas Jefferson. DNA from libraries of these samples is highly fragmented (Fig. 2A) and has a high level of cytosine deamination (Fig. 2B) – consistent with hair DNA sampled in the past. Assembled mitochondrial genomes of these libraries match the Thomas Jefferson H13a2b2 matriline. A consensus mitochondrial genome sequence from these libraries is identical to the L7 consensus mitochondrial genome including the private hot-spot and local private mutations (Fig. 1B). Using yhaplo (17), we find that the DNA from these libraries that maps to the Y chromosome is consistent with the Jefferson/Eston Hemings patriline at hundreds of haplotype-defining loci (Fig. 1C). Finally, pairwise analysis of the libraries from these hairs indicates that the DNA in each derives from the same individual (Fig. S1).

**Figure 2.**
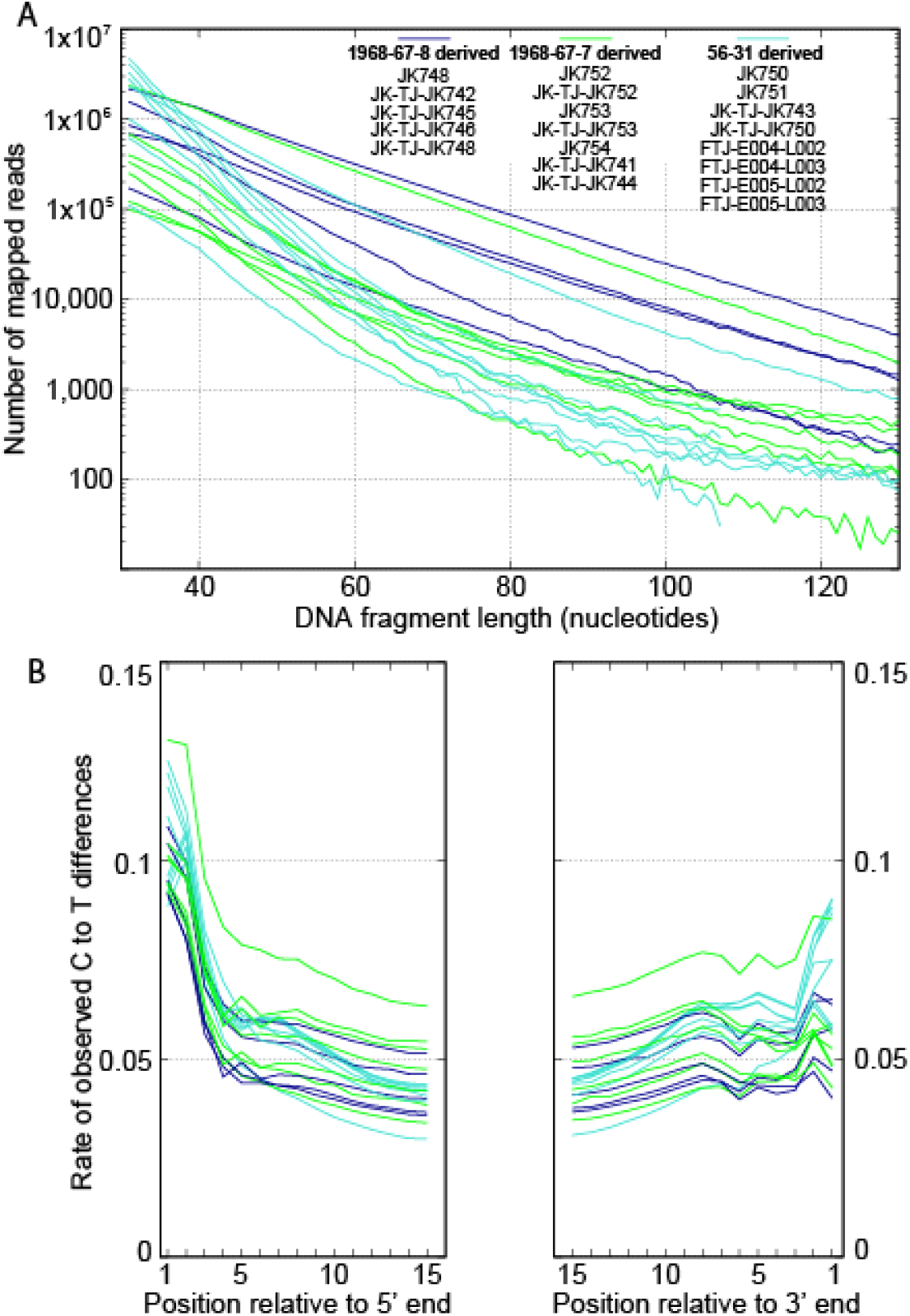
Features of DNA fragments extracted from Thomas Jefferson hairs. (A) DNA fragment length distribution, colored by source Thomas Jefferson sample. (B) Rate of observed C to T differences in aligned DNA data using same color scheme as in B.

In addition to the genomic assessment, historical provenance for these samples is strong. Annotations on the packets containing the identified hairs indicate they were clipped by Nicholas P. Trist following Thomas Jefferson’s death on July 4, 1826. Trist was Jefferson’s grandson-in-law and a co-executor of his estate. A letter, also in Monticello’s collection, from his daughter, Margaret Jefferson Trist Burke, indicates that the locks of Jefferson’s hair saved by her father were passed on to her. In the late 1890s she then divided that hair into packets that she annotated and gave to her children and a cousin. Decades later, descendants of those recipients sent those packets to Monticello (13).

Combining the DNA sequence data from all the verified Thomas Jefferson hair libraries, we achieved roughly 2-fold average coverage (5.9 billion base pairs of unique aligned sequence data) of the Thomas Jefferson nuclear genome and several hundred-fold average coverage of the mitochondrial genome. We note several auxiliary side products to those who may be interested in assessing Thomas Jefferson-related genetic genealogy: (1) a complete mitochondrial genome of Thomas Jefferson based on a high-coverage assembly, (2) a more complete catalog of Jefferson Y-chromosome markers, and (3) a genotype file suitable for genetic genealogy investigation (see Supplement section: A note on Thomas Jefferson DNA data availability).

### DNA samples from Jefferson and Hemings living descendants

To test various paternity scenarios, we worked with a professional family history researcher to identify living descendants in the Jefferson and Hemings families and then networked with members of those families. Because the amount of Jefferson DNA in a living descendant will decrease each generation, we prioritized sampling from individuals who were as few generations removed as possible from Thomas Jefferson and Sally Hemings. We visited with study participants and in almost all cases with close family present to explain the goals of the project. All the descendants and relatives were well-aware of the controversy over the paternity issue and had their own views and perspectives often well discussed in those sessions (for details see SOI). All subjects embraced Thomas Jefferson’s admonition to “follow truth wherever it may lead” (18) and agreed to participate with the aim of scientifically addressing the paternity question and potentially resolving the controversy—one way or the other (13). Volunteers submitted saliva samples for DNA extraction under Smithsonian Review Board Protocols HS 14052, HS 14052-02 and HS 24024 (SOI). In total 16 individuals were sampled, including multiple descendants of Eston and Madison Hemings, two of Sally Hemings’ sons (Fig. 3A). Descendants of her other two recorded children, son Beverly and daughter Harriet, remain unknown.

**Figure 3.**
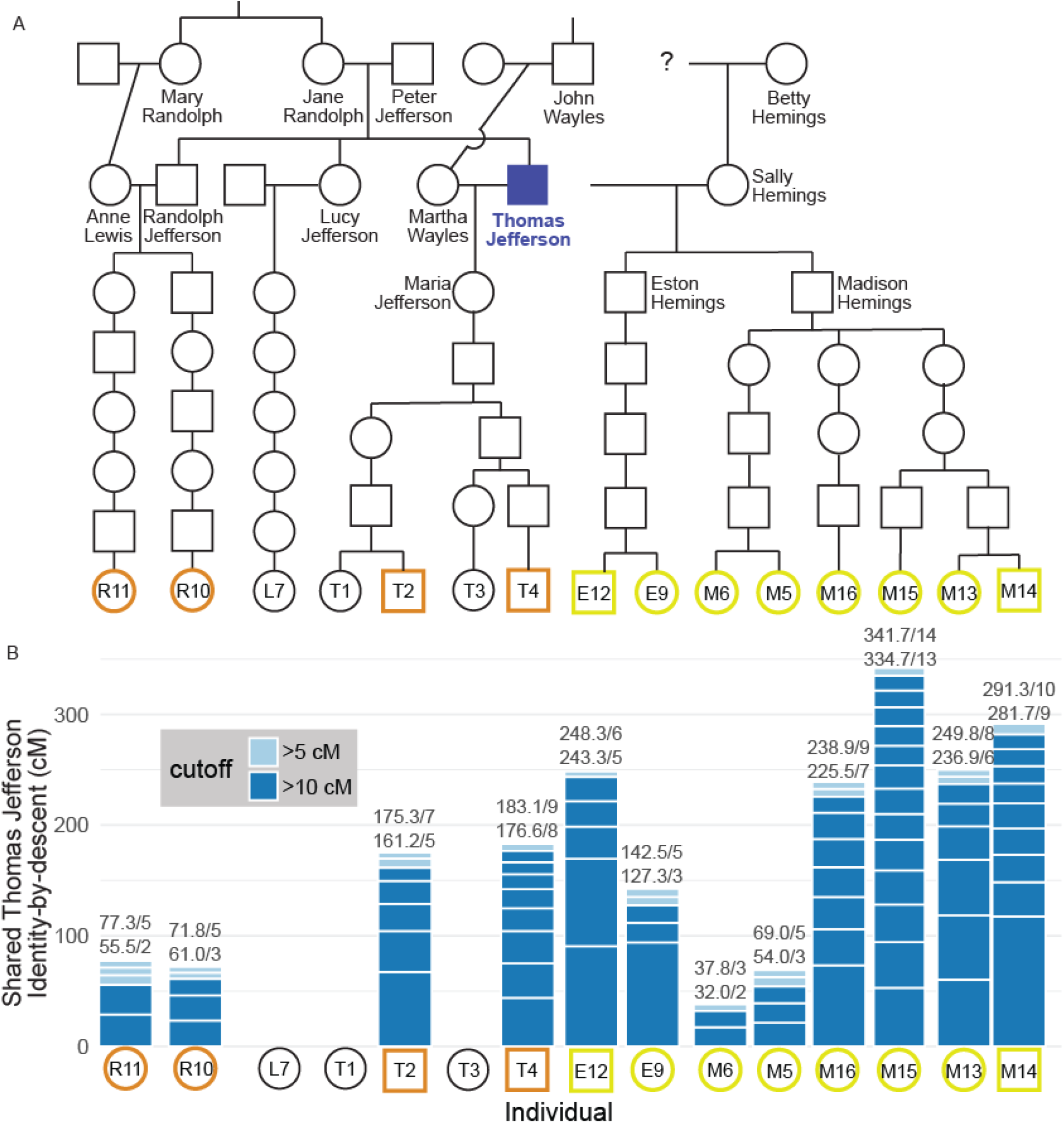
Jefferson and Hemings genealogy and shared DNA with Thomas Jefferson. (A) Relevant genealogy of the Jefferson and Hemings family. Males are squares. Females are circles. Known Thomas and Randolph Jefferson descendants who contributed DNA samples are in orange. Known Hemings descendants who contributed samples are in yellow. (B) Amount of DNA identified as identical-by-descent (IBD) with Thomas Jefferson (cM). IBD with Thomas Jefferson is not necessarily inherited directly from Thomas Jefferson. IBD segments may be co-inherited in Thomas Jefferson from a common ancestor with Thomas Jefferson such as those through his brother, Randolph Jefferson (R10, R11). Note that R10 is further related to the Jefferson lineage through the marriage of Lucy Jefferson’s granddaughter and Randolph Jefferson’s son.

We extracted DNA from the saliva samples and sequenced this DNA for genotype inference (SO). Samples for two of the descendants, T1 and T3 did not yield useful DNA and one of the participants later opted out of the study and their data was removed. For L7, the Thomas Jefferson matriline relative, we sequenced much less of the genome as the goal was only to assemble a full mitochondrial genome for comparison to the Jefferson hair samples. We called genotypes from the sequence data aligned to GRCh38 using GATK v4.5.0 (19).

### A Hidden Markov Model approach to finding Identity by Descent segments

We implemented a computational approach to infer genomic regions that are identical-by-descent (IBD) between the panel individuals and Thomas Jefferson. For the present-day descendants panel, we have diploid genotypes determined from high-coverage sequencing for most reference positions. In contrast, for the low-coverage Thomas Jefferson hair DNA data, we can only observe genomic positions with overlapping reads. To overcome these limitations, we developed a hidden Markov model-based (HMM) scan designed to compare high-quality genotypes with sparse observations from low-coverage sequence data (SOI). The HMM operates on SNP sites learned from a reference population that are observed in both the fully-genotyped individual and the low-coverage sample. The observation pairs for each SNP site serve as the model’s observed states. The HMM uses two hidden states representing the absence of IBD or the presence of any IBD between the two individuals. Cases of multiple overlapping IBD segments, such as those found between siblings or by coincidence, are handled by the IBD presence state. The model uses a high-resolution sex-average genetic map to set the hidden state transition probabilities between adjacent SNP sites (20).

We set the model’s emission probabilities using allele frequencies from a reference population to approximate the probability of matching Thomas Jefferson by chance. If no IBD exists between the individual and Thomas Jefferson, the probability of the observed base should follow its frequency in the population and be independent of the individual’s genotype. On the other hand, where IBD is present, matches between the DNA base observed in Thomas Jefferson hair and the individual’s genotype should occur more often. Note, the reference population frequencies are used only as a proxy for the probability of observing an allele from Thomas Jefferson, and we make no assumptions about the ancestry of the genotyped panel individuals. We chose the British in England and Scotland (GBR) subpopulation from the 1000 Genomes Project (21) as the reference population for the SNP sites and allele frequencies used in the scan but obtained similar results using a panel of Utah residents (CEPH) with Northern and Western European ancestry (CEU) (SOI, Table S3, Fig. S20-21). We used the forward-backward algorithm to decode the posterior probabilities and extracted Thomas Jefferson IBD segments for all panel individuals (SOI, Fig. S2). We separately applied 5 and 10 centimorgan (cM) minimum segment-match cutoffs to identify matching segments. Full implementation details and results of calibrating the method are available in the SOI (Fig. S3–S6).

### Thomas Jefferson shares DNA segments with descendants of Martha Wayles Jefferson and Sally Hemings

We ran the HMM scan comparing Thomas Jefferson DNA and the genotype of each of the panel individuals. We found extensive segments of DNA sharing in all panel individuals compared to unrelated control individuals from the 1000 Genomes Project (members of GBR and African Ancestry in Southwest US (ASW) populations) (SOI, Fig. S19, Table S2). In this section, we present the results of the first scan for each panel individual (Fig. 3, Fig. S7–18). Because the HMM only considers one low-coverage observation per site but we have ∼2-fold coverage from the hair samples, scan results will vary depending on the pseudo-random seed used to sample observations. We produced four additional independent scan results using the GBR reference panel to account for this variation (Table S3, Data Table S1). Although the analysis was run independently for each comparison, the genealogical and recombination histories are not independent for many of these individuals. For example, T2 and T4 are both five generations removed from their ancestors, Thomas and Martha Wayles Jefferson. However, T2 and T4 could only inherit Thomas Jefferson DNA that was present in their more recent common ancestor two generations prior.

Amongst the established Jefferson pedigree descendants, the number of IBD segments and total amount of DNA within them broadly follow expectations from the genealogy with variation due to recombination. The two descendants of Thomas Jefferson and his wife Martha Wayles through their daughter Maria Eppes have 175 cM (individual T2) and 183 cM (individual T4) of Thomas Jefferson IBD DNA—somewhat less than the 213 cM or 3.125% expected of a fifth-generation lineal descendant. The amount of detected Thomas Jefferson IBD DNA for R10 and R11 is also close to the expected amount, given the known pedigree. Individuals R10 and R11 are lineal descendants of Thomas Jefferson’s brother Randolph and Anne Lewis (his maternal first cousin). (Note, there is an additional known intrafamilial marriage for one of R10’s ancestors not depicted in the simplified pedigree in Figure 3). The amount of identifiable Thomas Jefferson IBD DNA in individuals with known Thomas Jefferson ancestry and relatedness is further evidence that the hair DNA used in this comparison derives from Thomas Jefferson and that the scan is well-calibrated. In addition to the IBD detection method implemented here, we ran ancIBD (22) on genotypes called from the Thomas Jefferson genome data using GLIMPSE2 (23). We find overall high concordance between Thomas Jefferson IBD segments called (Fig. S38) with our method generally spanning gaps between ancIBD segments.

We find the highest total amount of identifiable Thomas Jefferson IBD DNA within individual M15 (341.7 cM), a descendant of Sally Hemings through her son Madison. At 5.1%, this is considerably higher than the average expected IBD. Several other descendants of Madison Hemings, M13, M14 and M16 exceed the average expected amount. The highest amount of Thomas Jefferson IBD DNA for a descendant of Sally Hemings’ son, Eston, is in E12 (248.3 cM) and also higher than expected. Two of the smallest Thomas Jefferson IBD totals are found in Madison Hemings descendants M5 and M6. Considering the IBD totals found in the sampled descendants of Madison’s other two children, a parsimonious explanation is that Thomas Jefferson IBD was lost through the stochasticity of recombination and inheritance in the generations leading to M5 and M6. In all, these IBD totals reinforce the previous assertion that the father of Eston Hemings, and now Madison Hemings, must have been a close relative of Thomas Jefferson if not Thomas Jefferson himself.

### Jefferson paternity scenario comparisons

We evaluated three conceptually plausible paternity scenarios for Eston and Madison Hemings that are consistent with the Jefferson-Hemings Y chromosome match. These three scenarios are: (1) TJ - Thomas Jefferson is the father; (2) RJ - Randolph Jefferson, the brother of Thomas Jefferson, is the father; and (3) RJO - a male offspring of Randolph and thus nephew of Thomas Jefferson, is the father. Paternity scenarios with more distant patriline relatives are implausible given the levels of Thomas Jefferson IBD found among Hemings’ descendants. Given those levels, also implausible is a scenario where one of the Carr brothers – sons of Thomas Jefferson’s sister, are the ancestors of Madison Hemings’ descendants. Note that all three paternity scenarios would predict some amount of DNA to be shared between Thomas Jefferson and the descendants of Eston and Madison Hemings because the alternate fathers of Eston and Madison Hemings also share DNA with Thomas Jefferson. Specifically, Thomas Jefferson DNA sharing would be 50% for Randolph Jefferson (a full sibling) and 31.25% for a son of Randolph Jefferson, Thomas Jefferson’s nephew, since Randoph married the first cousin of Randolph and Thomas Jefferson, Anne Lewis. Thus, it is not the *presence* but the *quantity*, in both number and size, of Thomas Jefferson-matching DNA segments that distinguishes between these scenarios.

For statistical evaluation, it is possible that Eston and Madison Hemings had different fathers, although both fathers in the considered scenarios necessarily would have shared the same Jefferson Y chromosome. Therefore, comparison of paternity scenarios was done independently for the descendants of Eston Hemings and Madison Hemings. For each Hemings lineage (Eston and Madison), our panel includes multiple descendant individuals. These descendants share ancestors with each other more recently than their known or putative Jefferson ancestor. Thus, we cannot treat these individual observations as independent since they will share some historical recombination events that affect the number and size of Thomas Jefferson-matching (IBD) segments.

We pooled the Thomas Jefferson IBD observations by the two sampled Hemings branches, EH for Eston Hemings and MH for Madison Hemings, and merged any overlapping segments. This aggregate set represents the total unique IBD that remains in the descendants we sampled and the minimum Thomas Jefferson IBD present in Eston and Madison Hemings. Through simulations of the specific sampled genealogies, we examined the expected distributions for the number of Thomas Jefferson IBD segments and the sizes of these segments that would persist until the present generation (Fig. S27). We used the same simulation framework to evaluate the degree of Thomas Jefferson-matching DNA in the known Thomas Jefferson descendants in our panel (Fig. S22 & S23). We then compared the likelihoods of the observed Thomas Jefferson IBD DNA in the Hemings descendants under each paternity scenario.

We simulated the generations of recombination leading to the present-day Hemings descendants under each of the three paternity scenarios using ped-sim (24) (Fig. S24–S26). We specified the sex of each member of the genealogy and used a sex-specific high-resolution genetic map (20) and interference parameters to model recombination (25). We produced 20,000 simulation replicates for each scenario and performed the same grouping and merging of IBD segments described above. We used a likelihood function similar to the one described by Huff et al. (26) for their Estimation of Recent Shared Ancestry (ERSA) method (SOI). This method models the number of IBD segments shared between a pair of relatives and the distribution of their individual sizes independently and uses the combination to determine the likelihood. We modified the ERSA likelihood to use a generalized Poisson distribution to model the number of merged segments shared with one individual and several relatives. We assume all matches are due to recent ancestry and use the minimum size threshold to reduce the effect of background matches. We obtained maximum likelihood estimates of the model parameters from the simulation for each Hemings branch, scenario, and minimum IBD segment size combination and computed the likelihood of the observations under each scenario (Table S4–S5, Fig. S28–S31). We found that the generalized Poisson distribution provided a better fit to the observed data but also obtained similar results using the standard Poisson distribution (Table S6 & S7). We also implemented an alternative likelihood that modeled only the total shared IBD using a gamma distribution and therefore did not require accurate estimates of the number of segments (Table S10, Fig. S36). We obtained results that were similar to those from the ERSA likelihood (Table S11). Here we present likelihood ratios and posterior probabilities derived from the ERSA likelihood using the generalized Poisson distribution because they are less reliant on estimated probabilities for extreme values with little support from simulations and the ERSA likelihood is slightly more effective at distinguishing between paternity scenarios in the Eston Hemings genealogy (Fig. S37).

To assess the appropriateness of this simulation framework, we first used it to compare the observed number and total length of Thomas Jefferson IBD segments amongst panel individuals of uncontested pedigree, i.e., the descendants of Thomas Jefferson and Martha Wayles and of Randolph Jefferson and Anne Lewis (Fig. 4A). We find that the observed, shared Thomas Jefferson DNA is well-modeled by the simulation framework. Similarly, we compared the observed Thomas Jefferson-shared DNA from Eston and Madison Hemings descendants using the simulation framework under each of the three paternity scenarios (Fig. 4B). The TJ scenario simulations model well the observed amounts of shared Thomas Jefferson DNA amongst descendants of Sally Hemings. Conversely, under the RJ and RJO paternity models, some individuals are extreme outliers.

**Figure 4.**
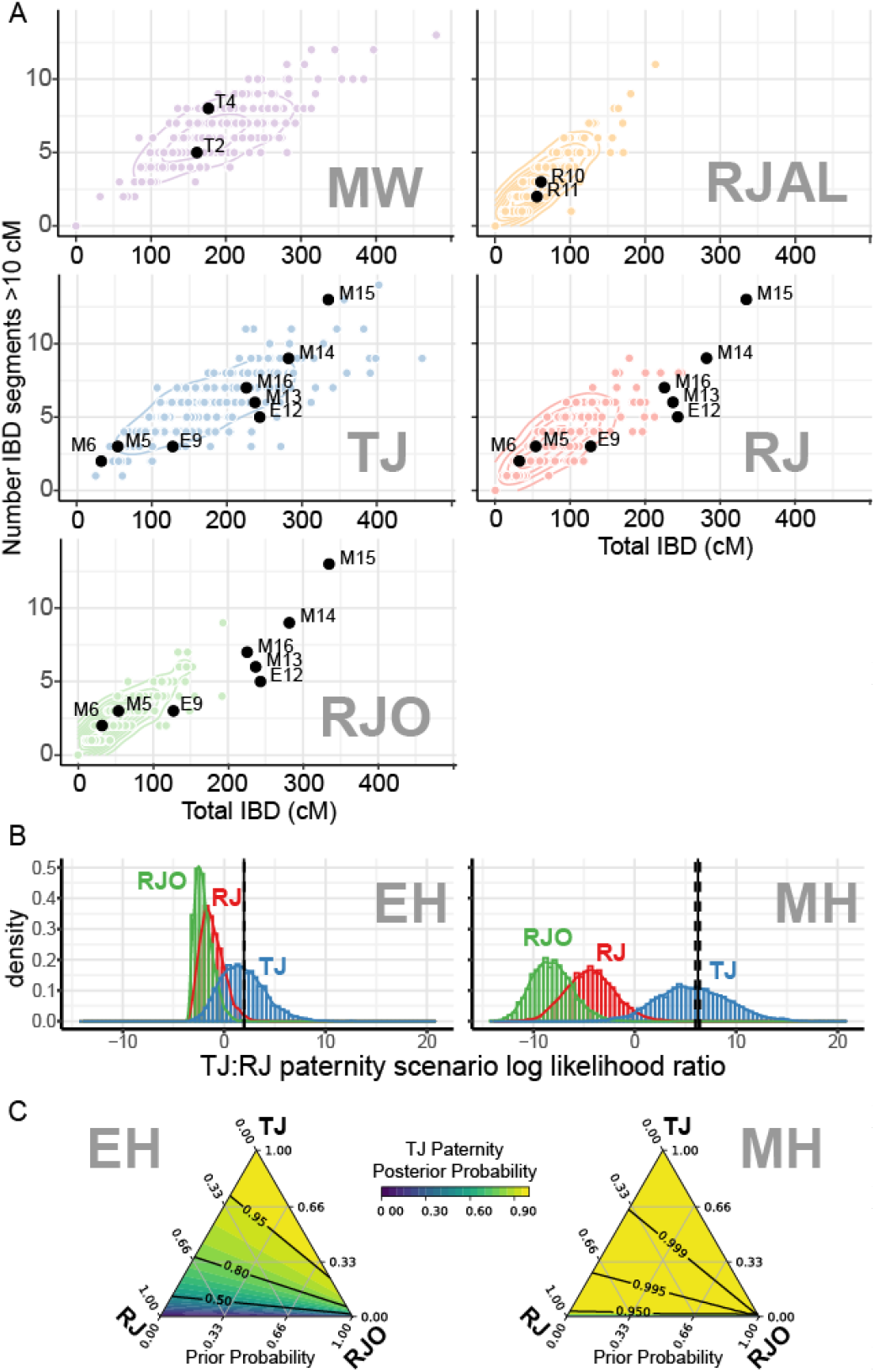
Statistical evaluation of paternity scenarios A, B. Number and size of Thomas Jefferson IBD segments found in panel individuals (labeled black dots) along with 200 pedigree simulations specific to each branch’s genealogy (colored points) and 2-dimensional kernel density estimate to illustrate the range and tendency of IBD sharing. **A**. Descendants of Thomas Jefferson and Martha Wayles (MW) and descendants of Randolph Jefferson and Anne Lewis (RJAL). **B**. Descendants of Sally Hemings through sons Eston (EH) and Madison (MH). Observations for the Hemings descendants are presented with simulations for 3 possible paternity scenarios: Thomas Jefferson (TJ), Randolph Jefferson (RJ), and a son of Randolph Jefferson and Anne Lewis (RJO). **C**. Likelihood ratios comparing TJ paternity versus RJ paternity for Eston Hemings (EH, left) and Madison Hemings (MH, right) based on Thomas Jefferson IBD segments merged across sampled descendants (10 cM minimum segment size) in each lineage. Observed likelihood ratios are shown as a vertical line with the central 95^th^ percentile interval (dashed lines) to mark their position on the log-scaled axis. The distributions of TJ:RJ likelihood ratios computed from pedigree simulations for each possible paternity scenario (TJ, RJ, and RJO) (4,000 simulation replicates each, independent of the set used to fit the likelihood parameters) are shown as histograms along with a smoothed density line. **D**. Ternary heat maps showing posterior probability of TJ paternity for Eston Hemings (EH, left) and Madison Hemings (MH, right) across the range of possible prior probabilities for 3 paternity scenarios considered (TJ, RJ, and RJO). Posterior probabilities were obtained using Thomas Jefferson IBD segments merged across sampled descendants (10 cM minimum segment size) and likelihood models for each paternity scenario based on pedigree simulations. Each axis on the triangle plot indicates the proportion of the prior probability assigned to a scenario and every point within the triangle represents a distinct set of prior probabilities that sum to 1.

Next, we computed the likelihood of the observed, shared Thomas Jefferson DNA in the Eston and Madison Hemings descendants under the three paternity scenarios using 5 cM and 10 cM minimum size thresholds. We used a likelihood ratio, or Bayes factor, to compare the likelihood of the Thomas Jefferson IBD observations under the TJ model with the other two models, RJ and RJO (Table 1). We implemented a resampling strategy to combine results from five HMM scans using the GBR reference panel with different pseudo-random seeds to determine the mean likelihood ratio and estimate central 95^th^-percentile intervals (Fig. S32 & S33, Data Table S2).

**Table 1.** Mean likelihood ratios and posterior probabilities comparing the likelihood of the three paternity scenario models for Eston Hemings (EH) and Madison Hemings (MH) using the 5 cM and 10 cM minimum segment size thresholds. Central 95^th^ percentile intervals are shown in parentheses. Likelihoods were estimated using the ERSA approach. Posterior probabilities were computed assuming equal prior probabilities for all three models. Results from 5 independent IBD scans were combined using a resampling strategy (N=2,000) to estimate the mean values and central 95^th^ percentile intervals.

|  |  | Likelihood Ratio |  | Posterior Probability |  |  |
| --- | --- | --- | --- | --- | --- | --- |
| Branch | Threshold (cM) | TJ:RJ | TJ:RJO | TJ | RJ | RJO |
| EH | 5 | 5.21<br>(4.98–5.39) | 53.77<br>(50.25–56.48) | 82.60%<br>(81.93%–83.1%) | 15.86%<br>(15.43%–16.44%) | 1.54%<br>(1.47%–1.64%) |
|  | 10 | 7.16<br>(7.01–7.29) | 62.26<br>(60.35–63.91) | 86.52%<br>(86.26%–86.74%) | 12.09%<br>(11.9%–12.31%) | 1.39%<br>(1.35%–1.43%) |
| MH | 5 | 424.74<br>(314.1–537.99) | 9.6e+05<br>(6.3e+05–1.3e+06) | 99.76%<br>(99.68%–99.82%) | 0.24%<br>(0.18%–0.32%) | 1.1e-04%<br>(7.4e-05%–1.6e-04%) |
|  | 10 | 512.82<br>(391.8–635.63) | 1.2e+06<br>(9.0e+05–1.6e+06) | 99.80%<br>(99.74%–99.85%) | 0.20%<br>(0.15%–0.26%) | 8.3e-05%<br>(6.1e-05%–1.1e-04%) |

For the EH descendants, we estimate the likelihood ratio comparing the TJ and RJ scenarios to be 5.21 (4.98–5.39 central 95^th^ percentile interval) and 7.16 (7.01–7.29) using the 5 cM and 10 cM thresholds, respectively, which can be interpreted as “substantial” or “positive” evidence favoring the TJ scenario over the RJ scenario (27). The ratios comparing the TJ vs RJO scenarios (53.77 and 62.26 for 5 and 10 cM respectively) indicate “strong” evidence and greater support for the TJ scenario over the RJO scenario. In the deeper sampled MH genealogy, the mean likelihood ratios comparing the TJ scenario to the RJ scenario exceeded 200 for both thresholds: 424.74 (314.1– 537.99) using the 5 cM threshold and 512.82 (391.8–635.63) with the 10 cM threshold. These values are often interpreted as “very strong” or “decisive”. The likelihood ratios comparing the TJ and RJO scenarios provide the strongest evidence to reject the RJO scenario for the MH genealogy.

We compared our observed likelihood ratios with the distributions of ratios obtained by computing the likelihoods for an additional 4,000 simulation replicates (Fig. 4C, S35), as it is possible for the RJ and RJO genealogies to produce likelihood ratios that favor the TJ scenario. We found in all cases that the observed log-transformed likelihood ratio falls close to the median of the distribution for the simulations representing the TJ paternity scenario. In contrast, the observed ratios represent extreme values for the distributions associated with the RJ and RJO genealogies.

We next computed the posterior probabilities of the three paternity scenario models given the observed matches, assuming equal prior probabilities for the scenarios. For the MH descendants we obtained mean posterior probabilities >99.7% for the TJ scenario with both match length thresholds: 99.76% (99.68%–99.82% central 95^th^ percentile interval) using the 5 cM threshold and 99.80% (99.74%–99.85%) with the 10 cM threshold. For the EH descendants, the TJ scenario is also associated with the highest mean posterior probabilities (>82%), while the RJ scenario retains a smaller but considerable proportion of the posterior probability. Finally, because it is difficult to set prior probabilities for these models, we computed the posterior probability of the TJ scenario across the range of possible priors (Fig. 4D, S34) and note that, after accounting for the new evidence, the TJ scenario is favored across the parameter space, except for the most extreme priors.

Finally, we estimated joint likelihood ratios and posterior probabilities under the assumption that Eston and Madison Hemings had the same father (Table 2). Under this assumption, we estimate that TJ paternity is 2,213 (1,614–2,814 central 95^th^ percentile interval) times more likely than RJ paternity using the 5 cM threshold and 3,672 (2,811–4,559) times more likely using the 10 cM threshold. We obtained similar posterior probabilities (∼99.95%) from both thresholds. When we allow for Eston and Madison Hemings to have different fathers and use a uniform prior where all combinations of paternity are equally probable, we estimate the posterior probability of TJ paternity for both Eston and Madison to be 82.4% (81.73%–82.9%) and 86.4% (86.09%–86.57%) using the 5 and 10 cM thresholds, respectively, and obtain a posterior probability of ∼99.96% that Thomas Jefferson was the father of a least one of Eston and Madison (SI, Tables S8 & S9) and therefore a posterior probability <0.1% that he was not the father of either one.

**Table 2.** Mean joint likelihood ratios and posterity probabilities assuming Eston Hemings and Madison Hemings shared the same father using the 5 cM and 10 cM minimum segment size thresholds. Central 95^th^ percentile intervals are shown in parentheses. Likelihoods were estimated using the ERSA approach. Posterior probabilities were computed assuming equal prior probabilities for all three models. Results from 5 independent IBD scans were combined using a resampling strategy (N=2,000) to estimate the mean values and central 95^th^ percentile intervals.

| Scenario | 5 cM |  | 10 cM |  |
| --- | --- | --- | --- | --- |
|  | TJ:X Likelihood Ratio | Posterior Probability | TJ:X Likelihood Ratio | Posterior Probability |
| TJ | 1.0 | 99.95%<br>(99.93%-99.97%) | 1.0 | 99.97%<br>(99.96%-99.98%) |
| RJ | 2,213.25<br>(1614.01-2813.39) | 0.05%<br>(0.03%-0.07%) | 3,671.52<br>(2811.25-4559.46) | 0.03%<br>(0.02%-0.04%) |
| RJO | 5.2e+07<br>(3.4e+07-7.3e+07) | 2.0e-06%<br>(1.4e-06%-3.0e-06%) | 7.7e+07<br>(5.6e+07-1.0e+08) | 1.3e-06%<br>(9.8e-07%-1.8e-06%) |

## Discussion

We describe analysis of DNA from hairs of Thomas Jefferson and comparison to known and putative descendants. The recovered DNA yielded nearly 2-fold average coverage of Thomas Jefferson’s genome. Several lines of evidence indicate that this DNA sequence is from Thomas Jefferson, including that it is a high-resolution match to the expected Thomas Jefferson patriline and matriline, and multiple samples with established provenance derive from the same individuals.

We also generated complete genome sequences from known descendants of Thomas Jefferson and his wife, Martha Wayles Jefferson, and putative descendants of Thomas Jefferson and Sally Hemings. In each case these descendants are five generations removed from the Thomas/Martha/Sally generation. In these individuals we identified regions of the genome that are inferred to be identical-by-descent with the DNA from Thomas Jefferson. Of note, while oral accounts assert that Martha Wayles and Sally Hemings were half-sisters through their father John Wayles, such a relationship, if correct, would not affect the direct comparisons of contemporary descendants with Thomas Jefferson or the calculation of IBD DNA shared with Thomas Jefferson. The amount of this Thomas Jefferson IBD is most consistent with Thomas Jefferson paternity of Eston and Madison Hemings compared to two conceptually plausible alternative scenarios that have been suggested because they are consistent with the Jefferson patriline match.

Over the course of this project, the technology for extracting and sequencing DNA from rootless hair shafts has advanced dramatically. It is now routine to generate multi-fold genome coverage from a few centimeters of a single rootless hair shaft (14). While the DNA in hair shafts does further degrade over time, hair has proven to be a remarkably stable storage vessel for DNA. Given past cultural practices of collecting and gifting locks of hair and their presence in numerous museum collections, the technology described here may potentially be deployed for many genealogical and historical forensic investigations (SOI).

Comparison of DNA to past relatives is particularly powerful in a genealogical context. For illustration, roughly 12.5% of one’s DNA is identical-by-descent with a great-grandparent. But second-cousins, who share this great-grandparent, will only share about 1.6% of their DNA through this common great-grandparent. We note that the Thomas Jefferson IBD segments identified in this study amongst Thomas Jefferson descendants are nearly completely unique amongst the panel individuals who are not close relatives. While they all have identifiable DNA-relatedness to Thomas Jefferson, this signal is small and scattered throughout their genomes, with little or none overlapping between panel individuals. Thus, the power to identify DNA ancestors by comparing *directly to DNA from the putative ancestor* is much higher than by comparing DNA amongst related descendants.

The data and analysis presented here help address a controversy that has existed for more than 200 years and often divided descendants, scholars, institutions, and the American people. Having recovered Thomas Jefferson’s DNA, the genomic evidence further reinforces strong historical scholarship indicative of Thomas Jefferson’s paternity of Sally Hemings children. The study thus advances methods of genomic science and provides greater certainty about early U.S. history. It also has personal and familial consequences for all the progeny of Thomas Jefferson—for those whose descent has historically been accepted, affirmed, or denied. Indeed, while results of this study were being shared among participants and their families, the Monticello Association formed a reconciliation committee engaging the children of several of the descendants (13). The findings may enable descendants of “founding father” Thomas Jefferson, whether born to Martha Wayles Jefferson or to Sally Hemings, and whether their ancestral legacy is one of slavery or freedom, to consider themselves as part of one and the same, complicated, American family.

## Acknowledgments

The loan of historical materials and analysis was made possible with the agreement and approval of the Smithsonian and the Thomas Jefferson Foundation, Monticello.

## Funding

The Smithsonian provided its own trust funds to support the research.

REG is co-founder and paid consultant of Astrea Forensics.

SHV is employed by Embark Veterinary, Inc.

BS is the chief science officer of Colossal Biosciences.

## Credit

Richard E. Green, Conceptualization, Investigation, Formal Analysis, Methodology, Resources, Writing—Original Draft Preparation, Writing-Review & Editing, Supervision, Project Administration

Samuel H. Vohr, Conceptualization, Investigation, Formal Analysis, Methodology, Software, Data Curation, Resources, Writing-Review & Editing, Visualization

Joshua D. Kapp, Investigation, Formal Analysis, Methodology, Resources Jane Ailes, Investigation

Samuel Sacco, Investigation, Formal Analysis, Methodology, Resources

Remy Nguyen, Investigation, Formal Analysis, Methodology, Resources

James Cahill, Investigation, Formal Analysis, Methodology, Resources

Peter D. Heintzman, Investigation, Formal Analysis, Methodology, Resources

Joanne Flores, Investigation

Logan Kistler, Investigation, Resources

Courtney A. Hofman, Investigation, Resources

,Robert C. Fleischer, Conceptualization, Investigation, Resources, Writing-Review & Editing, Supervision

Beth Shapiro, Conceptualization, Investigation, Formal Analysis, Methodology, Resources, Writing—Original Draft Preparation, Writing-Review & Editing, Supervision, Project Administration

Richard Kurin: Conceptualization, Investigation, Resources, Writing-Review & Editing, Supervision, Project Administration, Funding Acquisition

