## Supplemental Online Information for "The DNA legacy of Thomas Jefferson"

**Statistical evaluation of Jefferson paternity scenarios**……………………………... 40

**Human subjects information for DNA collection**……………………………………... 68

**A note on the use of ancient DNA from museum collections**…………………….. 71

**A Note on Thomas Jefferson DNA data availability**………………………………… 72

#### DNA extraction, library preparation, and sequencing

For this study, we processed hair strands over six years and updated our processing approach depending on the best available methods at the time. Table S1 contains a summary of methods used to generate each sequencing library included in the final dataset of Thomas Jefferson hair DNA.

For DNA extraction and isolation, we first selected 3 to 20 hairs to process for each extract. We either proceeded directly to digestion (Table S1, Pretreatment Column, No Treatment) or decontaminated the surface of the hair with a wash pretreatment step (Table S1, Pretreatment Column, Bleach). To decontaminate the surface, we submerged the hair in 0.5% sodium hypochlorite for 10 seconds, followed by submersion in 3 water baths for 10 seconds each. We extracted DNA from the hair with an overnight incubation in 500 µL of hair digest buffer (2% SDS, 10 mM Tris-HCl, 2.5 mM EDTA, 10 mM NaCl, 5 mM CaCl2, 40 mM DTT, and 2 mg/mL Proteinase K), previously described in Gilbert et al., (1). We isolated DNA using either the approach described in Dabney (2) or the column approach using binding buffer D described in Rohland (3). Finally, we quantified the DNA using the Qubit 1X dsDNA HS assay kit and a Qubit 4 fluorometer.

To prepare Illumina libraries, we used a pre-published version (Table S1, Library Method, Pre-SCR), or one of two modified versions (Table S1, Library Method, SCR-Mod1 or SCR-Mod2) of the Santa Cruz Reaction (4). The pre-published version is prepared as described in Kapp et al (4) with the following modifications: a final reaction volume of 80 µL, a 2:1 ratio of adapter to splint oligonucleotide, a 6:1 molar ratio of adapter:ssDNA input, and a 65:1 ratio of ET-SSB:ssDNA input. The first modified SCR version, SCR-Mod1, was performed as described in Nguyen et al (5). The second modified SCR version, SCR-Mod2, was performed as described in Nguyen et al (5) with the following modifications: 0.75 µL of 400,000 U/µL Hi-T4 ligase (NEB) rather than 0.25 µL of 2,000,000 U/µL T4 DNA ligase (NEB), adapter and splint oligonucleotides were trimmed to a 20 bp hybridization region (sequences below), and 15 µL of EBT plus 60 µL of SPRI beads were used during the initial incubation of the post-ligation clean.

SCR-Mod2 Oligonucleotides:
P5 Adapter (5'->3', IDT) = /5AmMC12/ACACGACGCTCTTCCGATCT
P7 Adapter (5'->3', IDT) = /5Phos/AGATCGGAAGAGCACACGTC/3AmMO/
P5 Splint (5'->3', IDT) = /5AmMC6/NNNNNNNAGATCGGAAGAGCGTCGTGT/3AmMO/
P7 Splint (5'->3', IDT) = /5AmMC12/GACGTGTGCTCTTCCGATCTNNNNNNNN/3AmMO/

Following ligation, we quantified the libraries with quantitative PCR using primers IS7 and IS8 (6). For each library, we amplified 1 µL of post-ligation library in 50 µL reactions (1X SYBR Maxima Green Master Mix, 1 µM IS7, and 1 µM IS8). Reactions were cycled with the following conditions: 95C for 10 minutes, followed by 40 cycles of 95C for 30s, 60C for 30s, and 72C for 30s. The fluorescence was measured at the end of each extension step.

We amplified and double-indexed each library using Amplitaq Gold 360 and the primers described in Kircher et al (7). For each library, we prepared 50 µL indexing reactions (1X Amplitaq Gold 360

Master Mix, 1 µM i5 primer, 1 µM i7 primer) and amplified for a library specific number of cycles according to the qPCR results. Reactions were cycled with the following conditions: 95°C for 10 minutes, followed by library-specific cycles of 95°C for 30s, 60°C for 30s, and 72°C for 60s, and a final extension of 72°C for 7 minutes.

After indexing PCR, we purified each amplified library using a SPRI ratio of 1.2X, quantified with a Qubit 1X dsDNA HS assay, and visualized with a Fragment Analyzer and the associated HSNGS kit.

**Table S1:** Summary of wet lab methods used to extract and convert hair DNA into sequencing libraries.

| Library Name | Extract Name | Sample Name | Hair Input strands | Pretreatment | Extraction Method | Library Method |
|---|---|---|---|---|---|---|
| JK748 | JK748 | TJ1968-67-8 | 3 | No Treatment | Dabney | Pre-SCR |
| JK750 | JK750 | TJ56-31 | 3 | Bleach | Dabney | Pre-SCR |
| JK751 | JK751 | TJ56-31 | 3 | Bleach | Dabney | Pre-SCR |
| JK752 | JK752 | TJ1968-67-7 | 3 | Bleach | Dabney | Pre-SCR |
| JK753 | JK753 | TJ1968-67-7 | 3 | Bleach | Dabney | Pre-SCR |
| JK754 | JK754 | TJ1968-67-7 | 3 | Bleach | Dabney | Pre-SCR |
| JK-TJ-JK741-L1 | JK741 | TJ-67-7 | 10 | Bleach | Dabney | SCR-Mod1 |
| JK-TJ-JK742-L1 | JK742 | TJ-67-8 | 10 | Bleach | Dabney | SCR-Mod1 |
| JK-TJ-JK743-L1 | JK743 | TJ-56-31 | 10 | No Treatment | Dabney | SCR-Mod1 |
| JK-TJ-JK744-L1 | JK744 | TJ-67-7 | 10 | No Treatment | Dabney | SCR-Mod1 |
| JK-TJ-JK745-L1 | JK745 | TJ-67-8 | 10 | No Treatment | Dabney | SCR-Mod1 |
| JK-TJ-JK746-L1 | JK746 | TJ1968-67-8 | 3 | Bleach | Dabney | SCR-Mod1 |
| JK-TJ-JK748-L1 | JK748 | TJ1968-67-8 | 3 | No Treatment | Dabney | SCR-Mod1 |
| JK-TJ-JK750-L1 | JK750 | TJ-56-31 | 3 | Bleach | Dabney | SCR-Mod1 |
| JK-TJ-JK752-L1 | JK752 | TJ1968-67-7 | 3 | Bleach | Dabney | SCR-Mod1 |
| JK-TJ-JK753-L1 | JK753 | TJ1968-67-7 | 3 | Bleach | Dabney | SCR-Mod1 |
| FTJ-E004-L002 | FTJ-E004 | TJ-56-31 | 20 | Bleach | Rohland | SCR-Mod1 |
| FTJ-E004-L003 | | | | | | SCR-Mod2 |
| FTJ-E005-L002 | FTJ-E005 | TJ-56-31 | 20 | Bleach | Rohland | SCR-Mod1 |
| FTJ-E005-L003 | | | | | | SCR-Mod2 |

#### Mitochondrial genome analysis

Saliva DNA was collected from panel member L7, a matriline relative of Thomas Jefferson. DNA from this sample was prepared for shotgun sequencing as described. We generated two independent libraries, Rel7_1b and Rel7_2b. The goal for this sample was to learn a complete mitochondrial genome. The library was sequenced on a 2x300 run Illumina run to ensure coverage across the mitochondrial genome.

Using `mia` (8), we assembled a consensus mitochondrial genome from the data from each library. The Rel7_1b library produced an assembly with 47.19 fold average coverage. The Rel7_2b library produced an assembly with 43.45 fold average coverage. Both assemblies were identical except for the length of a poly-cytosine stretch around rCRS position 300. Visual inspection of the aligned sequence data revealed likely heteroplasmy at this position. The consensus of CCCCCCCCTCCCCCC was included in the consensus.

The L7 consensus mitochondrial genome was analyzed using Haplogrep v3.2.1 (9) and determined to belong to haplogroup H13a2b2.

For each hair library, we assembled the mitochondrial genome using the same approach. We compared the Haplogrep call of the consensus mitochondrial genome to determine if it was the same as the L7 haplotype call.

All libraries of DNA derived from samples in 1956-31, 1968-67-7, and 1968-67-8 that generated mitochondrial consensus sequences with confident Haplogrep calls had the H13a2b2 call matching L7.

We further compared the mitochondrial consensus sequence from library JK754 (derived from 1968-67-7) to the mitochondrial consensus sequence from L7 and found that they were a complete match at all consensus positions.

#### Y-chromosome analysis of Thomas Jefferson and Eston Hemings descendant E12

We used the program `yhaplo` (10) to characterize the Y-chromosome from sequence data from Thomas Jefferson hairs and from individual E12 of the panel. E12 is a patriline descendant of Eston Hemings. Absent false-paternity in his genealogy, E12 will carry the Y-chromosome of the father of Eston Hemings.

We used the following bioinformatic procedure to prepare data for `yhaplo` analysis and to run yhaplo.

Because the `yhaplo` analysis requires data on the hg19 coordinate system, we first converted the hg38 aligned data to hg19 aligned data.

**Generate fastq file of reads aligned to hg38 Y-chromosome**

```
samtools view -h TJBAM chrY | \
      samtools view -Sb -o TJ.chrY.hg38.bam -
java -jar PIC SamTOFastq \
      I=TJ.chrY.hg38.bam \
      F=TJ.chrY.bam \
```

**Remap these reads to hg19 Y-chromosome**

```
bwa aln -t 36 HG19 TJ.chrY.fq.gz > TJ.chrY.hg19.sai
bwa samse HG19 TJ.chrY.hg19.sai TJ.chrY.fq.gz | \
    samtools view -Sb -o TJ.chrY.hg19.bam -
samtools sort -@ 4 -o TJ.chrY.hg19.s.bam TJ.chrY.hg19.bam
samtools index -@ 4 TJ.chrY.hg19.s.bam
```

648,552 non-duplicate reads mapped to hg19 with map-quality >= 20.
582,030 (89.7%) align to the Y-chromosome.

We assessed the depth-of-coverage in the Thomas Jefferson data at the ISOGG informative sites:

```
samtools mpileup -f /data/genomes/hs37d5.fa \
      -l isogg.2016.01.04.chr.pos.txt -q 20 -Q 20 \
      -a TJ.chrY.hg19.s.bam | cut -f 4 | sort -g | uniq -c
7376 0
3750 1
2024 2
1048 3
 594 4
 386 5
 259 6
 164 7
 140 8
```

```
 122 9
  81 10
…
```

**Generating yhaplo format genotype file from aligned data**
We downloaded ISOGG variant data (isogg.2016.01.04.txt) from the yhaplo repository data directory. Then, we converted this file to a format suitable for samtools mpileup -l option, i.e., a file with chromosome (Y) and position for each Y-chromosome variant position, generating `isogg.2016.01.04.chr.pos.txt.`

We generated an mpileup file for Y-chromosomse genotype calling from the hg19 Y-chromosome aligned data, filtering only base calls from mapped reads with map-quality >= 20 and base-quality >= 20

```
samtools mpileup -f HG19 -l isogg.2016.01.04.chr.pos.txt \
        -q 20 -Q 20 -s -a TJ.chrY.hg19.s.bam > TJ.chrY.hg19.isogg.mp
```

We then ran a custom script for consensus genotype calling from this mpileup data. The script outputs a file suitable for input to the yhaplo program implementing the following rule set:

1. There must be unanimous consensus among aligned bases.
2. To call a T allele requires >=1 observations in minus-strand aligned data.
3. To call an A allele requires >=1 observations in the plus-strand aligned data.
4. There must be no flanking gaps in any called base.
5. The coverage at a site must be >=1 and <=9.
6. Only observations with map-quality and base-quality >=20 are considered.

Note that rules 2 and 3 are designed to avoid calling Y-chromosome alleles that may be due to ancient-DNA associated cytosine deamination.

For the E12 data, we ran a similar routine to extract read data from the hg38 bam file and remapped to hg19 using bwa mem. The runs for converting to a `yhaplo` genotype file were similar except that the two rules specific for ancient DNA were not used and the coverage limits for adjusted (minimum of 4, maximum of 40) to account for the higher depth of coverage in the E12 data.

Finally, we ran `yhaplo` on both Thomas Jefferson and E12 genotype files using standard options. Data in Figure 1C was extracted from the `yhaplo` output files.

#### Bioinformatics analysis of hair and saliva DNA

##### Alignment of Saliva DNA from descendant panel

We ran the following sequence of steps to generate bam files for the DNA libraries prepared for each descendant.

HG38 = `GRCh38_full_analysis_set_plus_decoy_hla.fa`
INDIVIDUAL = the individual's identifier for this project
FQ1 and FQ2 = the forward and reverse fastq reads for this library
PIC = picard tools

### bwa version 0.7.17-r1188

```
bwa aln -t 22 -R '@RG\t[Read group information]` HG38 FQ1 FQ2 |
samtools sort -o INDIVIDUAL.s.bam

# picard version 2.25.7
java -jar PIC CleanSam \
      I=INDIVIDUAL.s.bam \
      O=INDIVIDUAL.s.C.bam \
      CREATE_INDEX=TRUE

java -jar PIC MarkDuplicates \
          I=bams/INDIVIDUAL.s.C.bam \
          O=bams/INDIVIDUAL.s.C.dm.bam \
          M=bams/INDIVIDUAL.dup.metrics.txt \
          OPTICAL_DUPLICATE_PIXEL_DISTANCE=2500 \
          CREATE_INDEX=TRUE
```

##### Genotype calling of descendant samples

Genotype calls for short variants on autosomes were generated using GATK v 4.5.0 (11) with the following steps and parameters:

1. **BaseRecalibrator**, using known variant sites from **dbSNP build 138** specified with the `--known-sites` option.
2. **ApplyBQSR**
3. **HaplotypeCaller**, with intervals (`--L`) defined using variant sites from the **1000 Genomes Project,** padding (`--ip 20`) , and `--output-mode EMIT_ALL_ACTIVE_SITES`.
4. **CombineGVCFs**
5. **GenotypeGVCFs**, with the `--include-non-variant-sites` option.
6. **VariantRecalibrator** for SNPs (`-mode SNP`), using `--truth-sensitivity-tranche` values of 100.0, 99.9, 98.9, and 90.0, and the following annotations (-

an): QD, MQ, MQRankSum, ReadPosRankSum, FS, SOR, DP, and InbreedingCoeff.

7. **ApplyVQSR** for SNPs (`-mode SNP`) with a `-ts-filter-level` of 99.9.
8. **VariantRecalibrator** for indels (`-mode INDEL`), with `--max-gaussians 4` and the following annotations: QD, DP, FS, SOR, ReadPosRankSum, and MQRankSum.
9. **ApplyVQSR** for indels (`-mode INDEL`) with a `-ts-filter-level` of 99.0.

The following procedures were used to call genotypes on sex chromosomes:

For **female X chromosomes** and **male X PAR regions**, similar pipeline as above.

For **male X chromosome non-PAR region** and **Y chromosome**:

1. **HaplotypeCaller**, using the same parameters as above but with `--sample-ploidy 1`.
2. **CombineGVCFs**
3. **GenotypeGVCFs**, same parameters as above.
4. **VariantFiltration**, applying the following filters:

```
--filter-expression "QD < 2.0" --filter-name "QD_filter" \
--filter-expression "MQ < 40.0" --filter-name "MQ_filter" \
--filter-expression "FS > 60.0" --filter-name "FS_filter" \
--filter-expression "SOR > 3.0" --filter-name "SOR_filter" \
--filter-expression "MQRankSum < -12.5" --filter-name
"MQRS_filter" \
--filter-expression "ReadPosRankSum < -8.0" --filter-name
"RPRS_filter"
```

**Running Tilde on TJ hair DNA libraries**

We performed pairwise comparisons between TJ hair libraries using the program **tilde** to assess contamination with the following procedure:

1. Build the reference panel using the **GBR population** from the **1000 Genomes Project** with `vcftools --IMPUTE --mac 10`.
2. Run `make_obs_table` with options `-q 30 -Q 30`.
3. Run `pairs_in_range` with options `-r 100—40000 -d`.
4. Run `indv_test`.
5. Combine LLRs from chromosomes 1-22 for each comparison, and run `sample_pairs -t 1000`.

**Alignment of DNA sequence data from hair DNA**

DNA sequence data from each hair was aligned to the reference human genome using the following procedure:

We ran the following sequence of steps to generate a single, merged bam file for the DNA libraries prepared from Thomas Jefferson hair DNA.

HG38 = `GRCh38_full_analysis_set_plus_decoy_hla.fa`
INDIVIDUAL = the individual's identifier for this project
FQ1 and FQ2 = the forward and reverse fastq reads for this library
LIB = library name
Merged forward and reverse read pairs using SeqPrep.

### bwa version 0.7.17-r1188

```
bwa aln -t 36 HG38 LIB.M.fq > LIB.sai
bwa samse -r "@RG\t[Read group information]" HG38 LIB.sai LIB.M.fq |
samtools view -Sb - | samtools sort -O BAM -@ 2 -o LIB.hg38.M.s.bam

# picard version 2.25.7
java -jar PIC CleanSam \
      I=LIB.hg38.M.s.bam \
      O=LIB.hg38.M.s.C.bam \
      CREATE_INDEX=TRUE

java -jar PIC MarkDuplicates \
          I=bams/LIB.M.s.C.bam \
          O=bams/LIB.M.s.C.dm.bam \
          M=bams/LIB.dup.metrics.txt \
          OPTICAL_DUPLICATE_PIXEL_DISTANCE=2500 \
          CREATE_INDEX=TRUE
```

Bam files from each library were then merged into a single bam file (TJ-241112-final.bam) for analysis. This file contains @RG (Read Group) tags for each aligned sequence for per-library analyses.

**Pairwise analysis of libraries using tilde**

The aligned sequence data from each library was compared against others using the `tilde` program (12). This program is designed to test the likelihood of the DNA sequence data under two models: (1) the data derive from the same individual and (2) the data derive from unrelated individuals. We used this analysis to test the hypothesis that data derived from different samples/libraries originates from the same individual. The likelihood ratio is dependent on the amount of data and other factors. In Fig. S1 (below), all pairwise library comparisons between libraries with the Jefferson Y chromosome haplotype and the Thomas Jefferson mitochondrial haplotype showed positive log-likelihood ratios, consistent with all libraries from all samples deriving from the same individual.

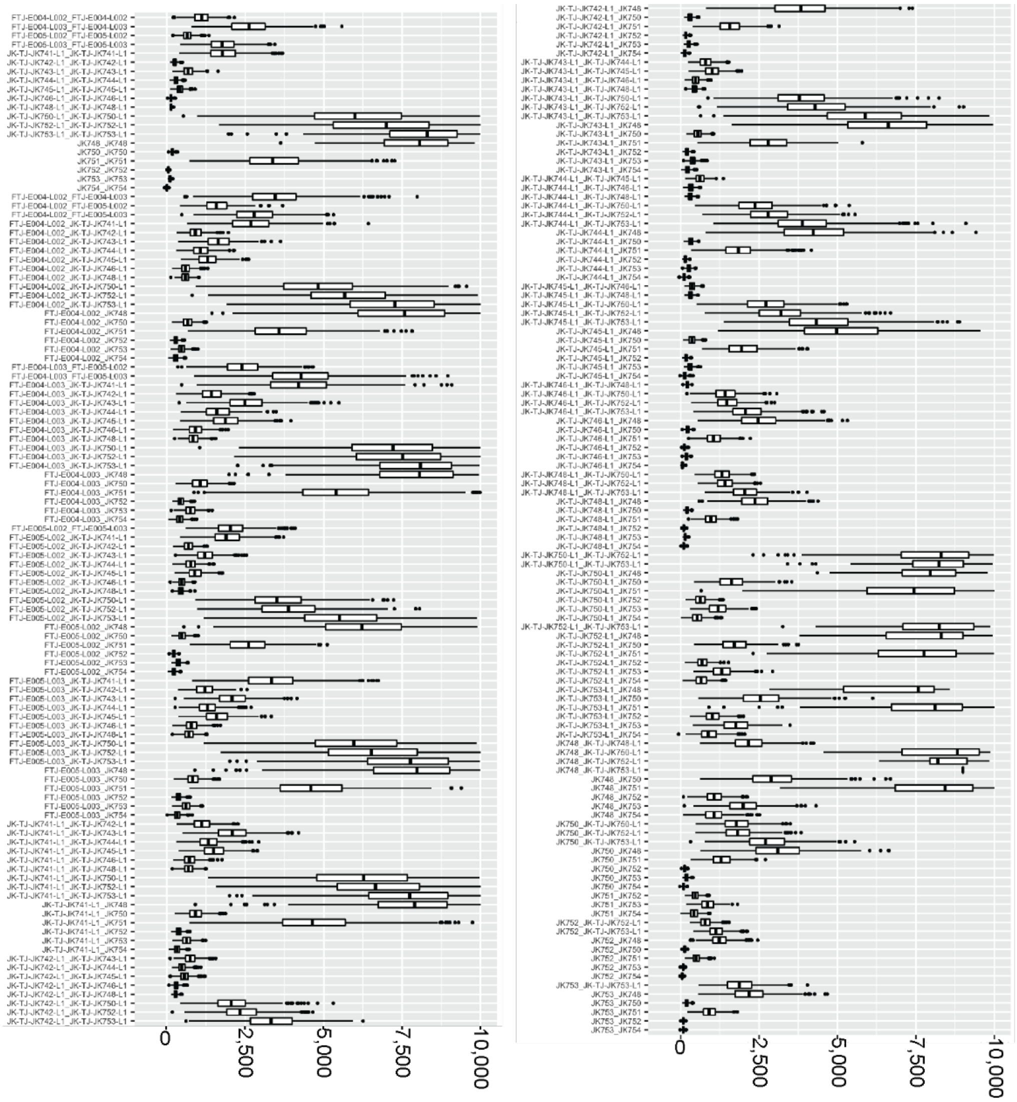

**Figure S1 – tilde comparison of all pairwise libraries.** Positive log-likelihood ratios indicate likelihood of DNA data is higher under the model that they derive from the same individual than under the model of deriving from unrelated individuals.

##### HMM scan for IBD detection

###### Overview

There are many existing tools for detecting IBD segments from high-quality genotype data (13, 14). Many of these methods were developed to detect distant relatedness within data sets with large numbers of individuals and assume a similar level of data quality for each sample. For ancient DNA, imputation strategies have been used to identify IBD from ancient DNA using existing tools (15-19). Ringbauer et al. developed *ancIBD* to detect IBD from ancient samples using genotype likelihoods (20) and Li et al. developed *clusIBD* for IBD detection in ancient and other poor-quality DNA samples (21).

Our comparisons required a method capable of detecting long, contiguous IBD segments (up to >100 cM) while being robust to missing data and sequence error in the hair samples while making full use of the high-quality genotypes from the descendant panel. With these requirements in mind, we developed a new IBD detection tool, TJBD, designed specifically for fine-scale detection of IBD segments between an individual with high-quality genotype information and a sparse, low-coverage DNA sample (0.1-2 fold coverage). We evaluated the power of TJBD to detect IBD using genotypes generated from pedigree simulations and show it can accurately detect shared IBD greater than 5 cM in size while maintaining a low false positive rate. Finally, we applied the scan to compare the combined Thomas Jefferson hair sequences to the panel of descendants and individuals from the 1000 Genomes Project as controls.

###### Description of IBD Scan

TJBD is a tool for fine-scale detection of IBD segments shared between two individuals, one with high-confidence genotype calls (hereafter referred to as the "high-coverage" individual) and one where only low-coverage sequence data is available (the "low-coverage" individual). The inputs for the scan are a VCF file containing the genotype calls for the high-coverage individual, a BAM file containing mapped sequences for the low-coverage individual, allele frequencies and linkage disequilibrium statistics for a reference panel, chosen to represent the population from which the low-coverage individual originates, and a genetic map to estimate recombination probabilities and measure the size of detected segments. This information informs the emission and transition probabilities of a Hidden Markov Model (HMM) that is used to estimate the probability of shared IBD across the length of a chromosome. Finally, discrete IBD segments are extracted from these posterior probabilities.

TJBD uses a simplified view of identity by descent to detect long, contiguous IBD segments. For two diploid individuals, there are 4 possible combinations of haplotype phases that may share IBD and it is possible for more than 1 combination to share IBD in the same region. However, multiple IBD is considered rare outside of full siblings (20). Additionally, for our comparisons, our genealogical research did not uncover any additional relationships that could produce overlapping TJ IBD in any of the descendants in our panel. Instead of modelling all possible IBD configurations, we designed TJBD to distinguish between only two cases, one where no IBD exists between the two individuals and one where any IBD is shared. This model allows us to use unphased genotypes for the high-coverage individual and to avoid phase-switch errors which may artificially fragment the detected IBD segments. TJBD may conflate overlapping IBD, either due to the population background or false positive signals, on opposite haplotype phases as larger segments (22) but we reduce the impact of these cases by applying a minimum segment size threshold.

Our scan focuses on the 22 autosomal chromosomes (chromosomes 1-22) of the human reference genome where all individuals in our comparison have two copies of each chromosome. We exclude the X chromosome from our scan for two reasons. First, scanning the X chromosome would require extending the HMM to handle the cases where the X chromosome is in a haploid state when low-coverage or the high-coverage individual is a genetic male. Second, statistical interpretation of the results of an X chromosome scan would require special consideration of its unique demographic properties (23). Finally, IBD segments on the X cannot be transmitted through two successive male ancestors (23). Our genealogy research indicates that only R10 and R11 could have possibly inherited IBD segments due to recent ancestry shared with Thomas Jefferson. T1 and L7 may also share X IBD, but we do not have genotype data for those individuals.

TJBD uses single nucleotide polymorphisms (SNPs) as markers to detect Identity by descent segments. The SNPs used in the scan are selected using a pre-existing reference panel of individuals, which is treated as a proxy for the population from which the low-coverage sample derives. We also use this panel to obtain allele frequencies for each SNP and linkage disequilibrium statistics. For the high-coverage individual, we extract the called genotypes at the marker sites from the VCF files described in the previous section. For the low-coverage individual, we examine the sequences aligned to the reference that overlap the marker sites and make a single base observation where possible. If there are multiple reads that overlap a marker position, we choose 1 read at random to make the base observation, unless the number of overlapping reads exceeds a fixed maximum coverage parameter. We include only SNPs markers that have a high-coverage genotype call and a low-coverage base observation in the scan.

Before running the scan, we apply a linkage disequilibrium filter similar to the one described in (24) to remove redundant markers that may produce false positives driven by linkage disequilibrium rather than recent shared ancestry. We apply this filter by first calculating the haplotype $r^2$ statistic between each marker and each of the 500 surrounding markers in the reference panel using VCFtools (25). After the base pair observations have been made from the low-coverage sample and before the HMM scan is run, we examine the $r^2$ values between pairs of observed markers, in order, along the chromosome. If the $r^2$ between the two markers exceeds a threshold value (0.30) and neither marker has been previously excluded, the marker with the higher minor allele frequency is excluded from the HMM scan.

#### Description of Hidden Markov Model

##### Hidden States

TJBD uses a Hidden Markov Model (HMM) to decode the posterior probability that an IBD tract overlaps a SNP marker. The model has two hidden states, 0 and 1, which indicate the absence of IBD (no-IBD) and the presence of any shared IBD (IBD). For a set of $N$ SNP markers distributed across the length of a chromosome, the hidden state is represented using $s_{1..N} \in \{0,1\}$.

##### Emission Probabilities

Let $o_{i\in 1..N} \in \{A,C,G,T\}$ be the base observed from the low-coverage sample at marker $i$ and $f_{o_i}$ represent the frequency of the observed base $o_i$ in the reference population. For the high-coverage individual, $g_{i,j\in\{0,1\}} \in \{A,C,G,T\}$ is a vector of pairs containing the two alleles called at marker $i$. We define $m_i \in \{0,1,2\} = \sum_{j=0,1} o_i = g_{i,j}$ as the number of times the low-coverage base is found in the genotype at the $i$-th marker.

If there is no IBD shared between the two individuals, the probability of observing the low coverage base is independent of the high-coverage genotype and depends only on the frequency of the allele in the population and the probability that the base was read erroneously ($\varepsilon$). For the scan, we set the error rate to 1%.

$$e_i(o_i|s_i = 0) = f_{o_i} + (1 - f_{o_i})\varepsilon$$

If a marker is in an IBD segment, the probability of observing the low coverage base depends on the number of times it is found in the high-coverage genotype $\{0,1,2\}$ and the probability of observing it by chance on the opposite haplotype phase (i.e., the one that does not share IBD).

$$e_i(o_i|s_i = 1) = \frac{1}{2}[\frac{1}{2}m_i + (f_{o_i} + (1 - f_{o_i})\varepsilon)]$$

##### Transition Probabilities

The transition probabilities of our HMM are set using a recombination map that describes the distance between adjacent SNP markers in centimorgans ($d_{i,i+1}$). By definition, if two markers are 1 centimorgan apart, there is a 1% chance of a recombination occurring between the two markers with each generation on average. Using a constant representing the expected number of generations separating the two individuals ($g$), we obtain the probability of any recombination occurring between the two SNP markers over $g$ generations.

$$r_{i,i+1} = 1 - \left(e^{-0.01d_{i,i+1}}\right)^{g-1}$$

If the previous marker was in the IBD state ($s_i = 1$), then any historical recombination would disrupt the IBD segment and cause a transition to the no-IBD state ($s_{i+1} = 0$). Conversely, if no historical recombination occurs, the next marker will remain in the IBD state ($s_{i+1} = 1$)

$$t_{i,i+1}(s_{i+1} = 0|s_i = 1) = r_{i,i+1}$$
$$t_{i,i+1}(s_{i+1} = 1|s_i = 1) = 1 - r_{i,i+1}$$

If the previous marker was in the no-IBD state ($s_i = 0$), a historical recombination may lead to a transition to the IBD state. However, this transition probability also depends on the number of ancestors $g$ generations in the past. For example, if $g = 3$ (great grandparent), there are $2^{3-1} = 4$ great grandparents contributing to the parental chromosome phase with IBD. If a historical recombination occurred between SNPs $i$ and $i + 1$, there are 3 ancestors from which SNP $i + 1$ may originate, but only 1 ancestor would produce the IBD state (the low-coverage individual).

$$t_{i,i+1}(s_{i+1} = 1 | s_i = 0) = \frac{1}{2^{g-1} - 1} r_{i,i+1}$$
$$t_{i,i+1}(s_{i+1} = 0 | s_i = 0) = 1 - \frac{1}{2^{g-1} - 1} r_{i,i+1}$$

We found the value for $g$ did not have a noticeable effect on the results of our scans. After several generations have passed, IBD segments tend to become small enough that they are likely to be passed on completely intact or not at all. As a result, there is considerable overlap in the IBD segment size distributions for ancestors a few generations apart and many chromosomes will share no IBD with a given ancestor.

#### Calling IBD Segments

We use the forward-backward algorithm to decode the posterior probability of the IBD state for each SNP marker from the HMM and extract discrete IBD segments using the following procedure. First, we identify runs where the posterior probability exceeds a predetermined threshold value (referred to as the “upper” threshold). Second, any markers surrounding these runs where the posterior probability is greater than a lower, secondary cutoff are included in the called segment (the “lower” threshold). If two regions that satisfy the upper threshold are separated by a region where the lower threshold is always met, the combined region will be called as a single segment. Third, and optionally, a minimum length in centimorgans can be applied to remove short segments. The final called IBD segments are the ranges greater than the minimum length wherein all SNP posterior probabilities are greater than the lower threshold and the posterior probability for at least 1 SNP exceeds the upper threshold (See Figure S2).

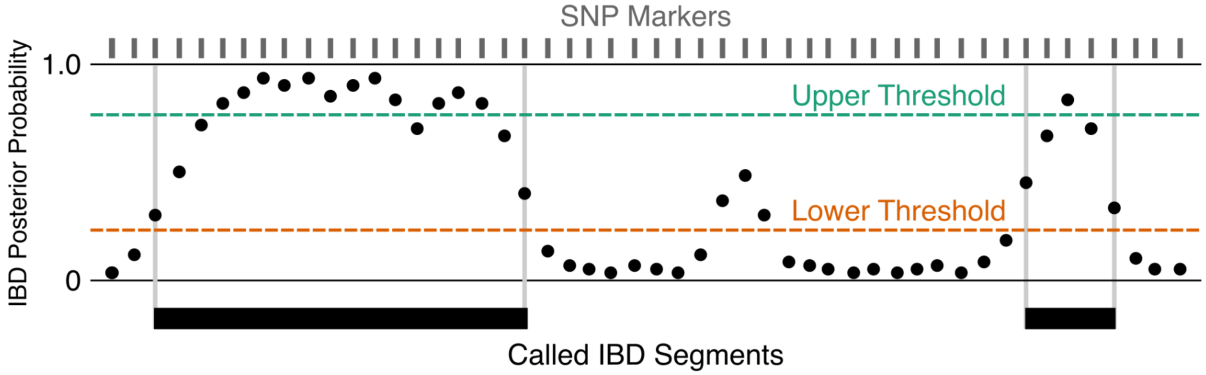

**Figure S2 - Schematic of IBD segment calling criteria.**

#### Software Implementation and Availability

TJBD is written in Python 3. It uses HTSlib (25, 26)   through pysam (https://github.com/pysam-developers/pysam) for handling BAM and VCF file input, numpy (27) for vector operations and pandas (DOI: 10.25080/Majora-92bf1922-00a) to manipulate tabular data. The source code is available under the MIT License at https://github.com/svohr/tjbd.

##### Evaluating Scan Power & Accuracy

#### Pedigree Simulations

We evaluated the power and accuracy of TJBD to detect identity-by-descent segments using genotype data from pedigree simulations. We used ped-sim (28) to simulate the transmission of genetic segments from a single ancestor through 5 generations to a direct descendant. This pedigree is analogous to the genealogy relating Thomas Jefferson to descendants T2 and T4 and the descendants of Sally Hemings under the TJ paternity scenario. Although we consider other possible pedigrees in this study, we chose to use this single pedigree to evaluate TJBD because it covers the widest range of expected segment sizes and simulating additional generations of recombination would produce more chromosomes devoid of IBD segments.

```
# Ancestor and descendant separated by 5 generations
#
# Relevant comparison:
#   *_g1_b1_i1 (ancestor)
#   *_g6_b1_i1 (present day descendant)
#
def pedigree1 1 6
1 1
2 0 1
3 0 1
4 0 1
5 0 1
6 1
```

**Figure S3 - Ped-sim def file**. The input file describes a simple pedigree with five generations of descent from an ancestor to a descendant.

For each pedigree, we ran ped-sim using the cross-over interference model with the parameters inferred by Campbell et al.. Recombination rates were set using the sex-specific genetic map from (29) and the sex of each simulated individual was randomly assigned. To produce genotype data from the pedigree simulations, we provided ped-sim with composite individuals to use as founders. We generated these composite individuals using the method described in (Rodriguez et al. Parente2 (30) to avoid introducing latent IBD to our experiments. Briefly, we made composite individuals from the 1000 Genomes British in England and Scotland (GBR) population by copying genotypes from each source individual along tiled segments of 0.2 cM across each chromosome. We disabled genotype error simulation in ped-sim and instead simulated base pair sequencing error in the low-coverage individual separately. We generated 200 ancestor/descendant pairs with simulated genotypes.

To evaluate the power and accuracy of the IBD scan and to determine the threshold values to use in the Thomas Jefferson and descendant panel comparisons, we applied the TJBD scan to the set of 200 simulated ancestor/descendant pairs using SNP markers and a reference panel based on the GBR population (see below). We scanned each pair, simulating ~1.5-fold coverage of the ancestor individual, and calling segments based on the upper and lower thresholds described above. To assess the effects of different values for these parameters, we ran each scan multiple times, sweeping through the range of upper probability thresholds, from 0 to 1, exclusive, in 0.1 steps, in combination with all applicable lower thresholds (0 to the upper threshold value, exclusive, in 0.1 steps). We did not apply a minimum segment size requirement.

Figure S4 shows true positive, false positive, and false discovery rates for detection of IBD segments and correct assignment of IBD state by marker position using each combination of upper and lower threshold values. With no minimum size enforced, the detection of IBD segments is entirely determined by the upper threshold. For segment detection, we found that the upper thresholds among the parameters we explored yielded false positive rates between 87.6% and 92.7%, and false discovery rates between 0.3% and 9.2% (note: we quantified error using FDR for segments because the definition of a true negative segment is ambiguous in this case. The FDR is dependent on the prevalence of IBD segments in the scenario of two directly-related individuals separated by 5 generations). Positional accuracy, how often TJBD correctly infers that a marker is in an IBD segment, is primarily determined by the lower threshold. To choose the parameters that best balance the positional true positive and false positive rates, we computed the root-mean-squared error as;

$$RMSE = \sqrt{\frac{\sum_i (t_i - o_i)^2}{n}}$$

Where $t$ is the true state of the marker, $o$ is the observed/called state and $n$ is the total number of markers (lower panel). We found that the scan performed similarly across a wide range of thresholds, and the lowest RMSEs were found when the lower threshold was set around 0.3-0.4 and the upper threshold was set around 0.4-0.5. From these observations we fixed the upper detection threshold to 0.4 and the lower to 0.3 for the TJ/Rel scans.

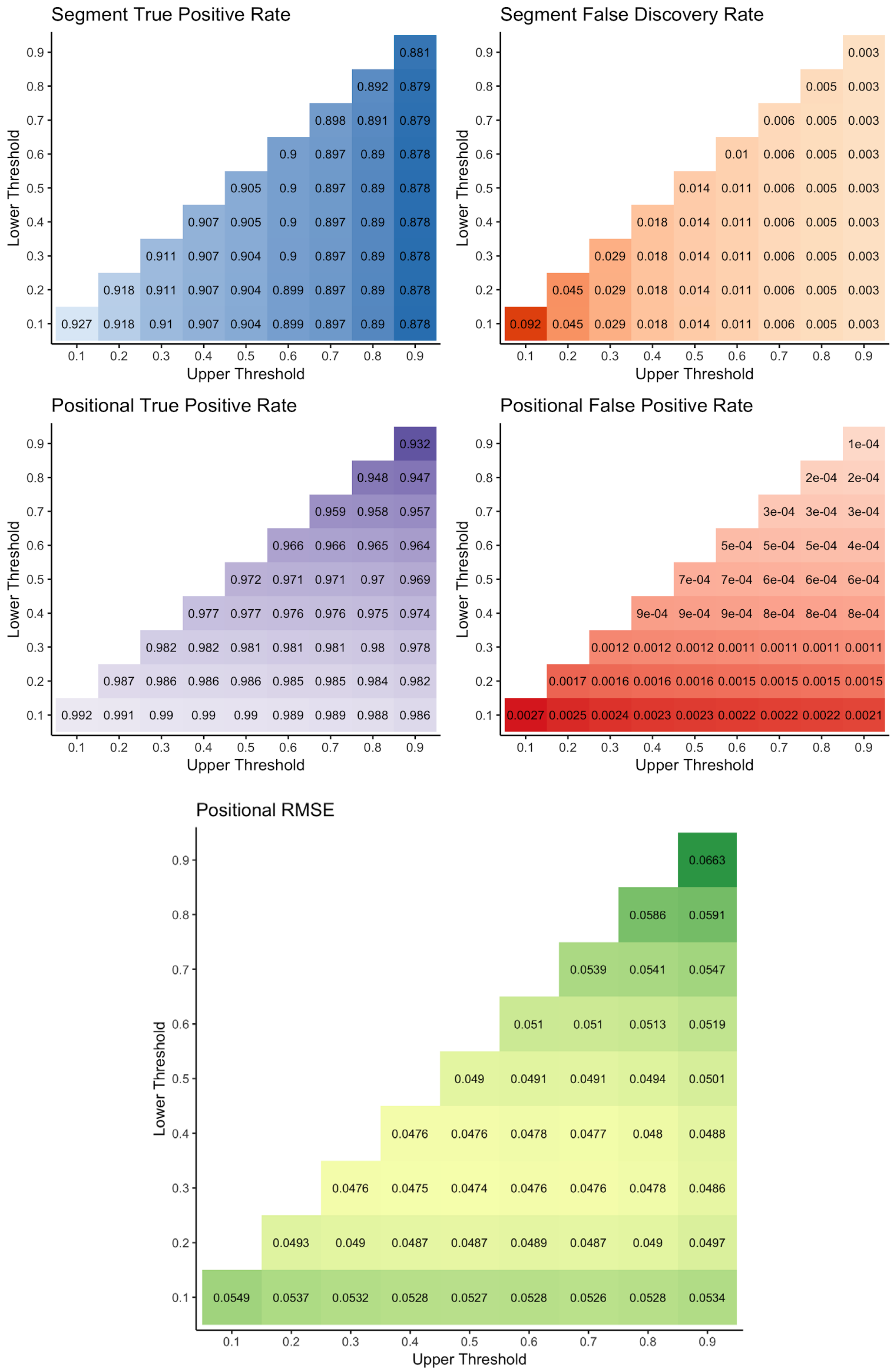

**Figure S4 – Sensitivity and false positive/discovery rates of TJBD.** Top panel: Rates are shown by segment and position for the TJBD scan across various upper and lower thresholds. Lower panel: root-mean-squared error for positional IBD calls.

We next examined the effect the size of the true IBD segment had on the power of the scan to detect it. We found that TJBD had power to detect nearly all segments 6 cM or greater in size (99.6% using the strictest threshold, 1,645 out of 1,652 total segments), regardless of the upper threshold applied. For smaller segments, we found that power dropped depending on the size of the segment and the choice of threshold, where lower thresholds had more power to detect segments of each size group. Segments less than 1 cM size were virtually undetectable using any threshold. To assess the accuracy of the scan's results, we examined the proportion of called IBD segments that overlap a true IBD segment (at least 50% of the called segment overlapping a true segment). We found our IBD scan had consistently high accuracy for segments longer than 5 cM (99.3% using the most permissive threshold, 1,739 out of 1,752 detected segments). We observed a drop in accuracy for segments smaller than 5 cM and further reduced accuracy associated with less stringent detection thresholds. Finally, we examined the relationship between the lower probability threshold and differences between the sizes of the detected segment and the true IBD segment. Fixing the upper probability threshold to 0.6, most detected segments are within 2 cM of the true segment size. We observed that more stringent lower probability thresholds tend to underestimate the size of the underlying segment, while less stringent tend to overestimate the size. Setting the lower threshold to 0.30 yielded the average difference closest to 0 (-0.2 cM).

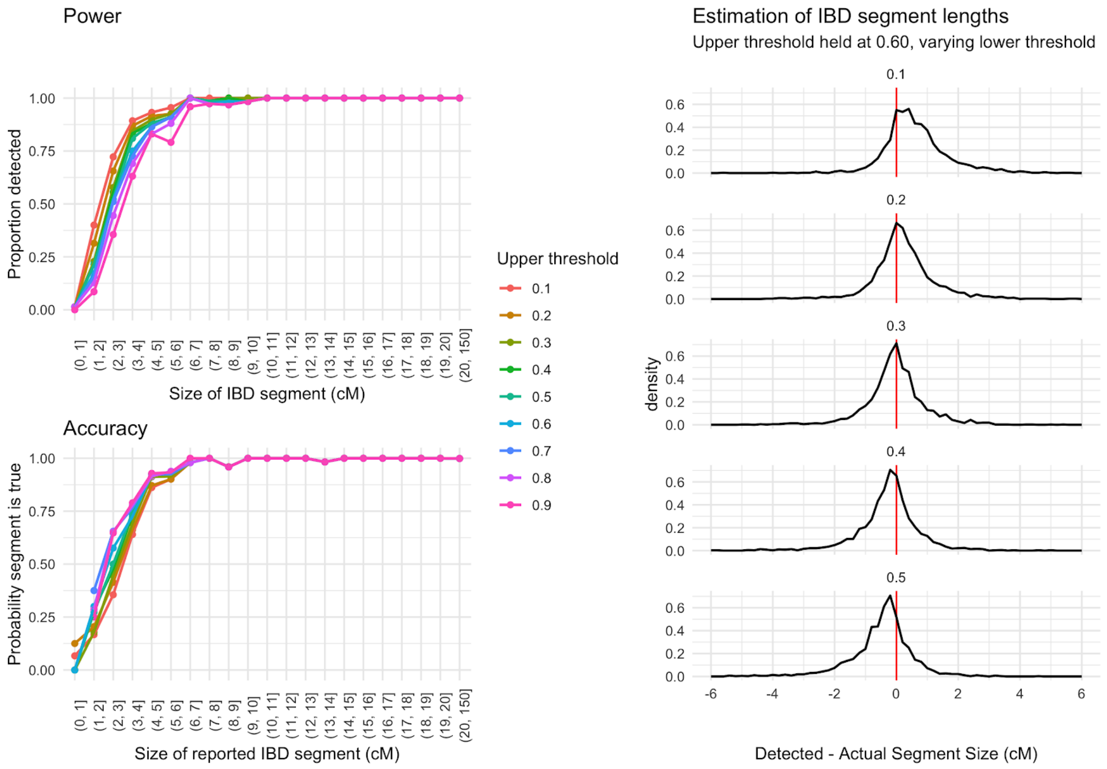

**Figure S5 – Power and Accuracy of TJBD HMM.** Left panel: Analysis of 200 TJBD HMM scans of simulated ancestor/descendant pairs. TJBD consistently detects segments larger than 6 centimorgans in size, with few false positives, regardless of probability thresholds used. For segments smaller than 6 cM, both accuracy and power to detect segments decreases with segment size. Lowering the upper threshold increases power to detect while decreasing accuracy. Right panel: Decreasing the lower threshold widens the flanking region included in the detected segment. We found that lower thresholds around 0.3 resulted in estimated sizes within ± 2 cM of the actual segment size without significant bias towards over or underestimation of segment size.

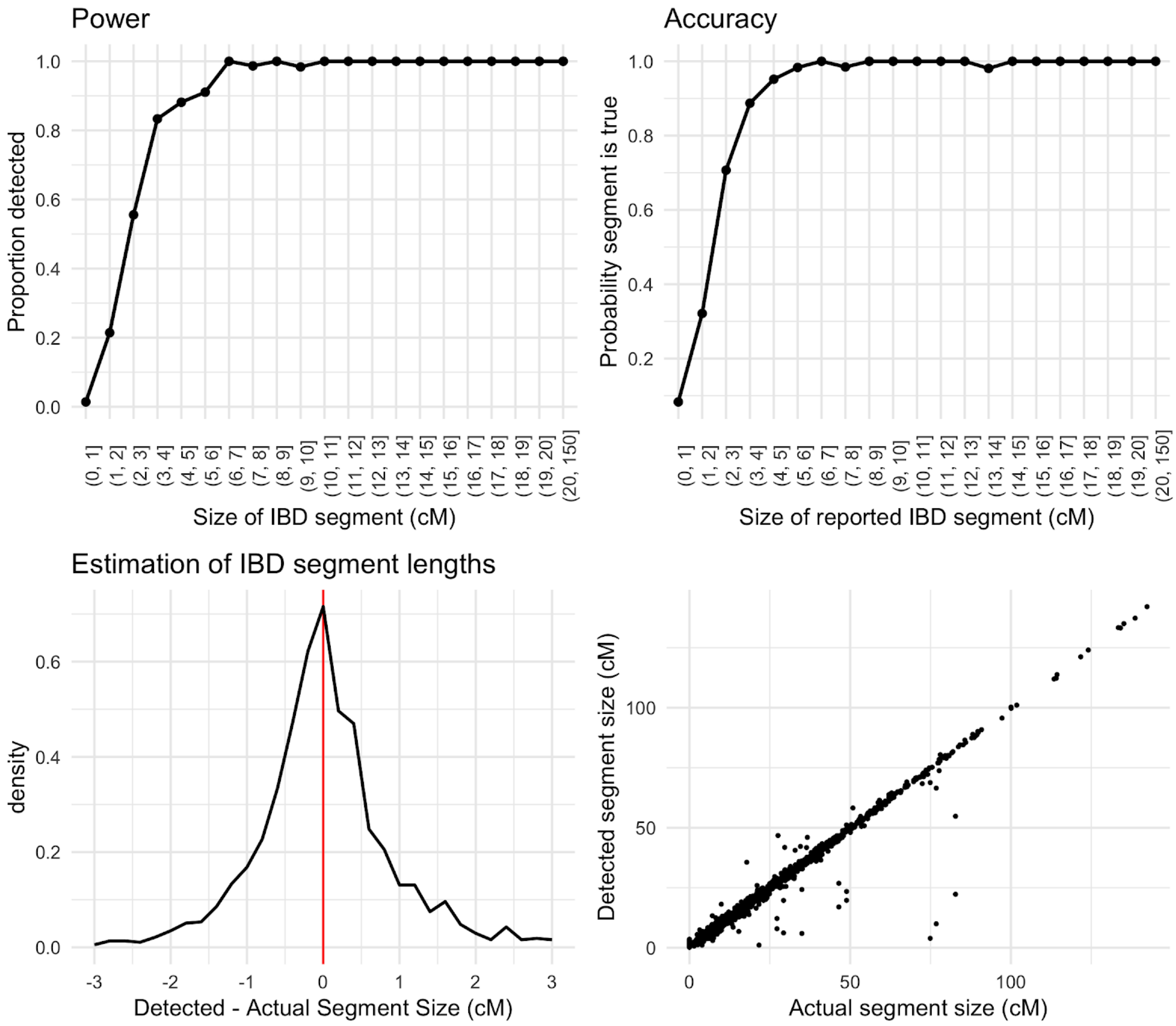

**Figure S6 - Power and accuracy of TJBD scan.** Scan results are shown under parameter ranges selected for TJ and Relative panel comparisons (upper threshold = 0.4, lower threshold = 0.3). **Top:** Power, probability a segment is detected, and accuracy, probability a detected segment is true, are consistently high for segments larger than 5 cM. **Bottom:** Sizes of detected segments closely match sizes of underlying actual segments, usually within ± 2 cM. In a few cases, some segments are detected as multiple segments (most points found below the diagonal).

### IBD Scan Results

#### Reference Panel and SNP Filtering

We used data from the 1000 Genomes Project's British in England and Scotland population (GBR) to select the single-nucleotide polymorphism (SNP) markers used in the scan (91 individuals). We used VCFTools to filter the phase 3 GRCh38 VCF files to include only biallelic SNPs with a minor allele count of 2 or greater in the GBR population. To minimize the effect of spurious mappings of the short sequences from the hair samples, we further filtered SNP sites to exclude regions listed in the UCSC Genome Browser's simpleRepeat track (31) https://hgdownload.soe.ucsc.edu/goldenPath/hg38/database/simpleRepeat.txt.gz, and applied a mappability filter created for GRCh38 using the seqbility tool (https://github.com/lh3/misc/tree/master/seq/seqbility) to include only sites where all overlapping 30-mers map uniquely in the reference and no single mismatch would cause a 30-mer to map elsewhere as determined by BWA (32). After applying these filtering criteria, we retained 5,250,333 autosomal SNP markers for the scan. In addition to the GBR panel, we constructed a second reference panel based on the 1000 Genomes Utah residents (CEPH) with Northern and Western European ancestry (CEU) population using the same method but increased the minimum minor allele count of 4 to correspond with the larger sample size (N=178 in the CEU panel versus 91 in the GBR panel).

We applied additional filtering steps through TJBD for each panel individual independently before running the scans for IBD segments. Our model requires every SNP marker used in the scan to have an observed base from the low-coverage sample. Additionally, we applied a maximum coverage limit to exclude markers with more than 4 overlapping reads (set by assuming under a Poisson distribution and 2-fold average coverage ~95% of sites would be covered by 4 or fewer reads). We also required a minimum mapping quality of 30 and a minimum base quality of 30 for observations made from the low-coverage BAM file. After filtering based on the low-coverage sample, the linkage disequilibrium filter was applied to reduce the density of markers so that no pair of markers had a haplotype $r^2$ greater than 0.3, as calculated from the reference panel haplotypes. The final number of markers used to scan the 22 autosomes varied for each individual due to differences in called genotypes and ranged from 598,318 to 605,141 SNPs.

#### Descendants

We ran the TJBD scan comparing the sequences from the combined Thomas Jefferson libraries with each individual in our panel of descendants. IBD segments were called using an upper posterior probability threshold of 0.4 and a lower threshold of 0.3. We applied two minimum segment sizes, 5 and 10 centimorgans, to produce two sets of segments for further analysis. Figures S7–S19 show the results of the IBD scan for each descendant.

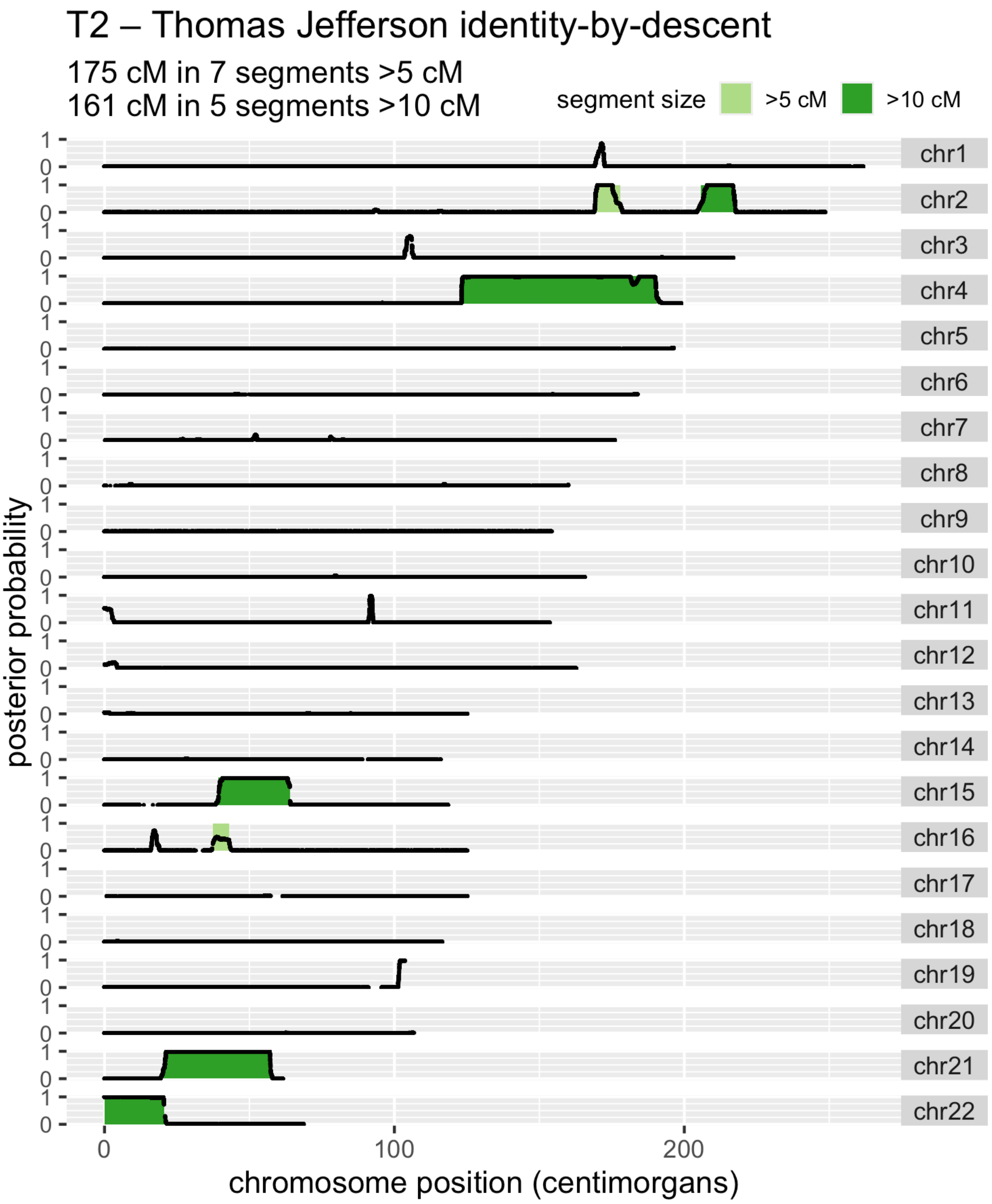

**Figure S7 - TJBD scan against T2 individual.** T2 is a known descendant of Thomas Jefferson and Martha Wayles. The posterior probabilities of IBD for each SNP are shown as black points. Detected IBD segments are shaded in light green if greater than 5 centimorgans in size and dark green if greater than 10 centimorgans in size.

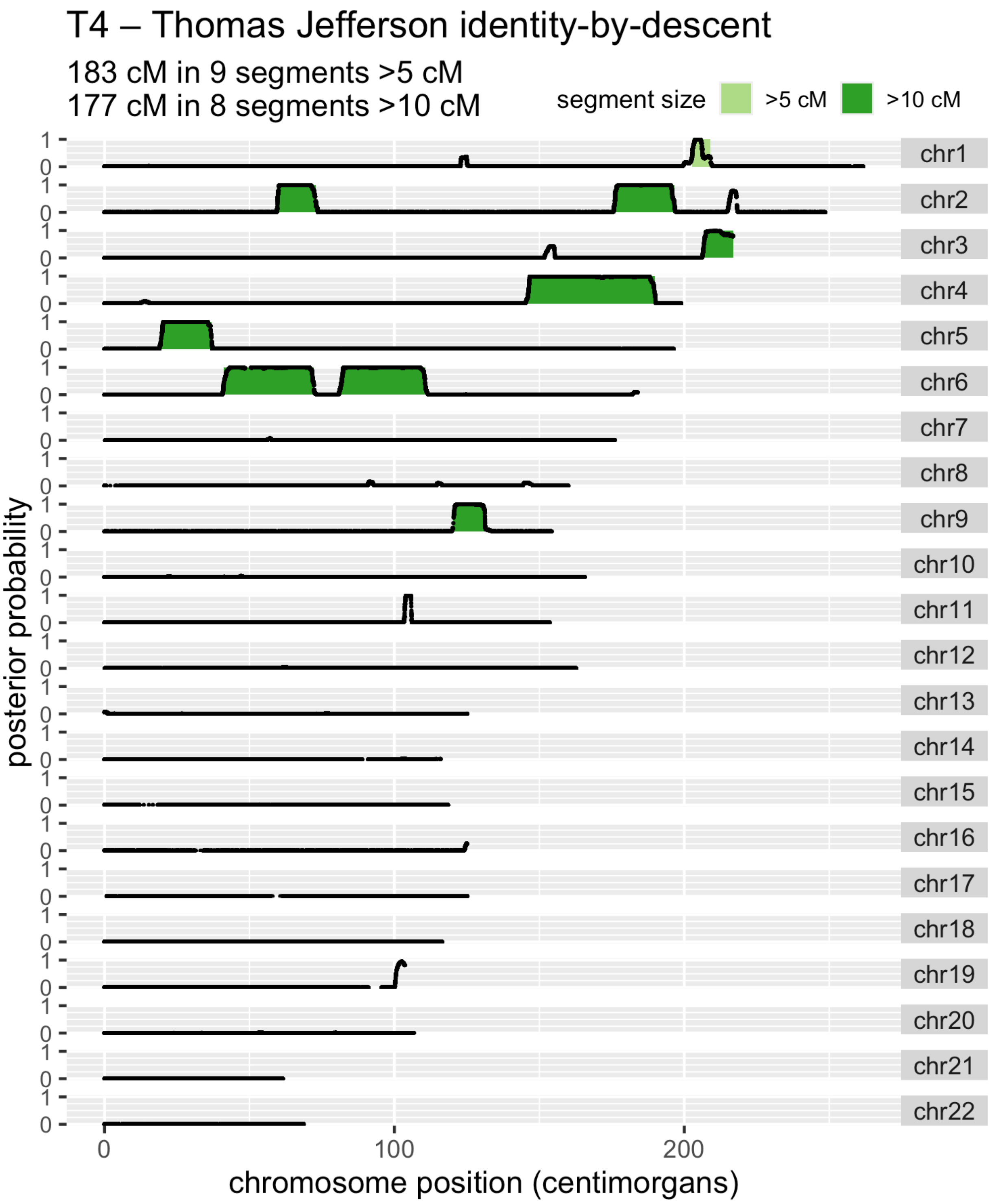

**Figure S8 - TJBD scan against T4 individual.** T4 is a known descendant of Thomas Jefferson and Martha Wayles. The posterior probabilities of IBD for each SNP are shown as black points. Detected IBD segments are shaded in light green if greater than 5 centimorgans in size and dark green if greater than 10 centimorgans in size.

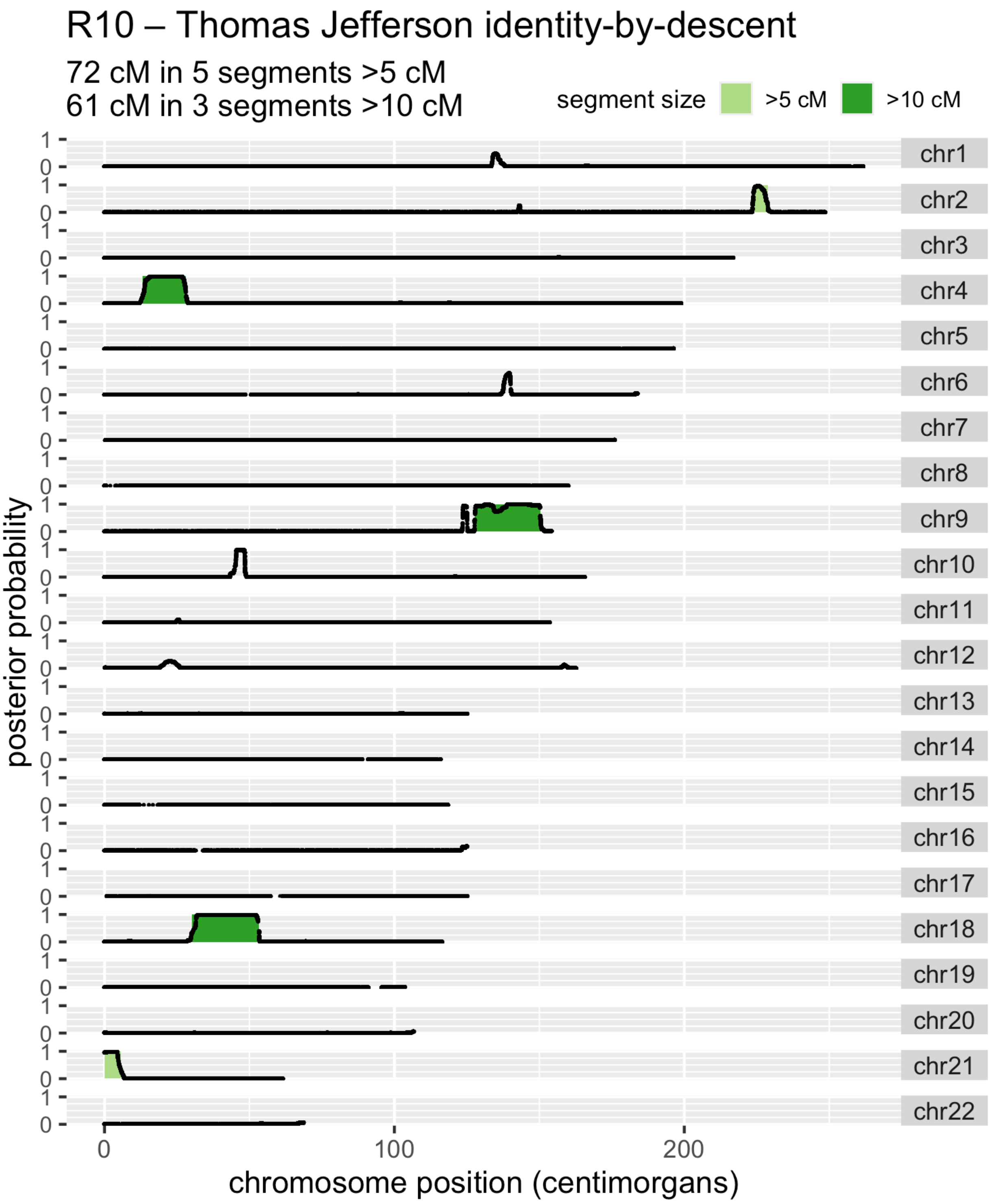

**Figure S9 – TJBD scan against individual R10**. R10 is a known descendant of Randolph Jefferson and Anne Lewis. The posterior probabilities of IBD for each SNP are shown as black points. Detected IBD segments are shaded in light green if greater than 5 centimorgans in size and dark green if greater than 10 centimorgans in size.

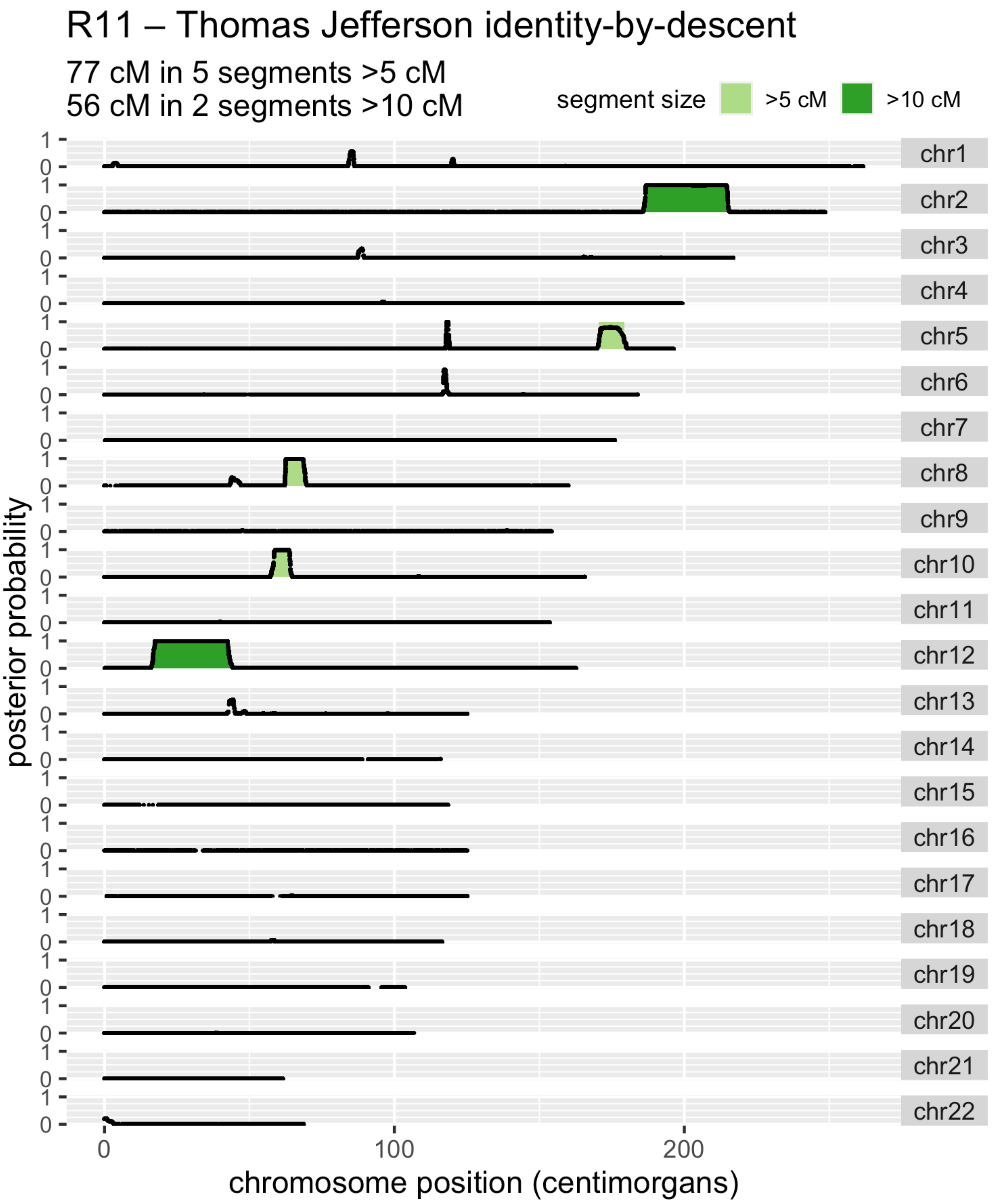

**Figure S10 - TJBD scan against individual R11**. R11 is a known descendant of Randolph Jefferson and Anne Lewis. The posterior probabilities of IBD for each SNP are shown as black points. Detected IBD segments are shaded in light green if greater than 5 centimorgans in size and dark green if greater than 10 centimorgans in size.

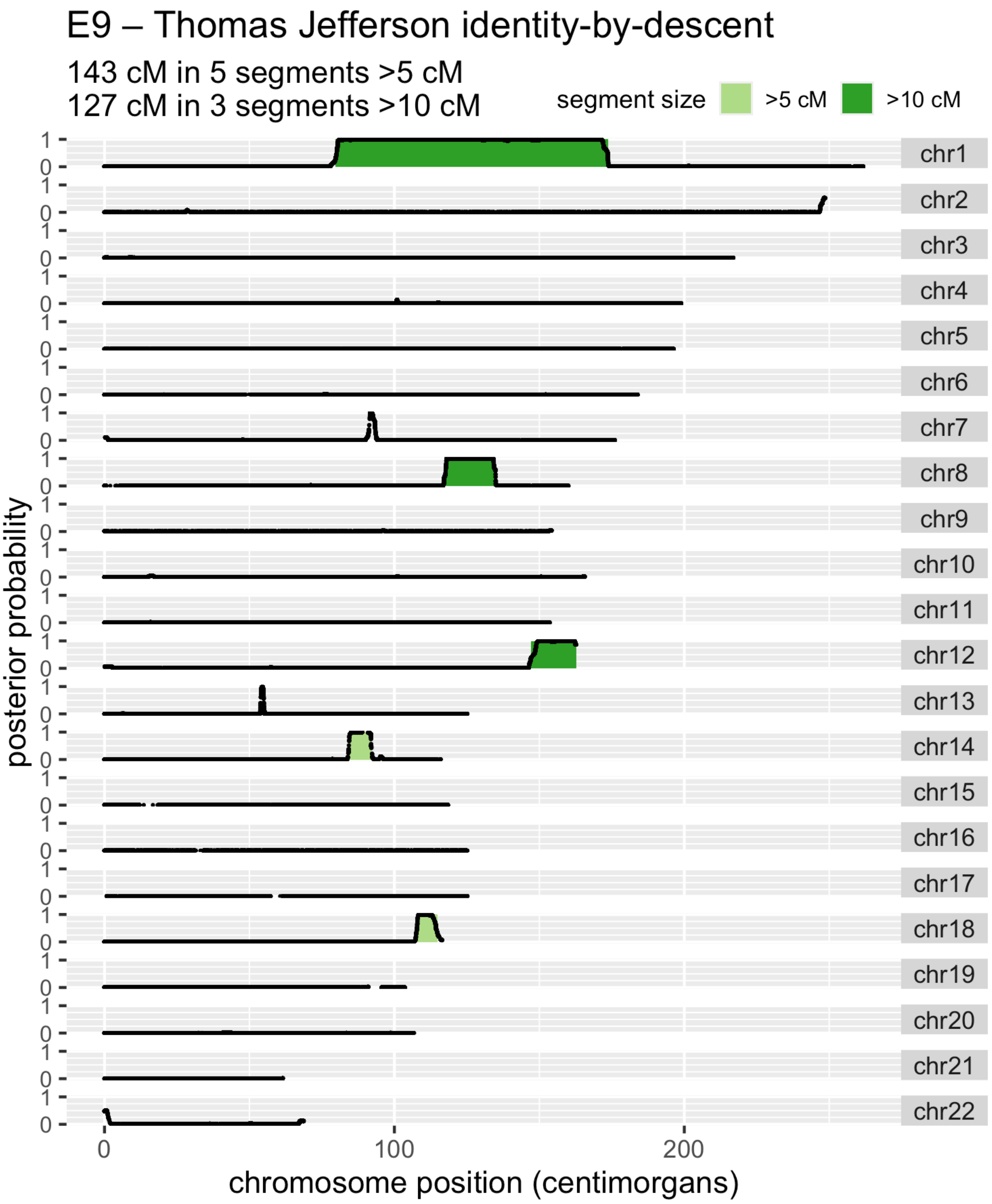

**Figure S11 – TJBD scan against individual E9.** E9 is a known descendant of Eston Hemings. The posterior probabilities of IBD for each SNP are shown as black points. Detected IBD segments are shaded in light green if greater than 5 centimorgans in size and dark green if greater than 10 centimorgans in size.

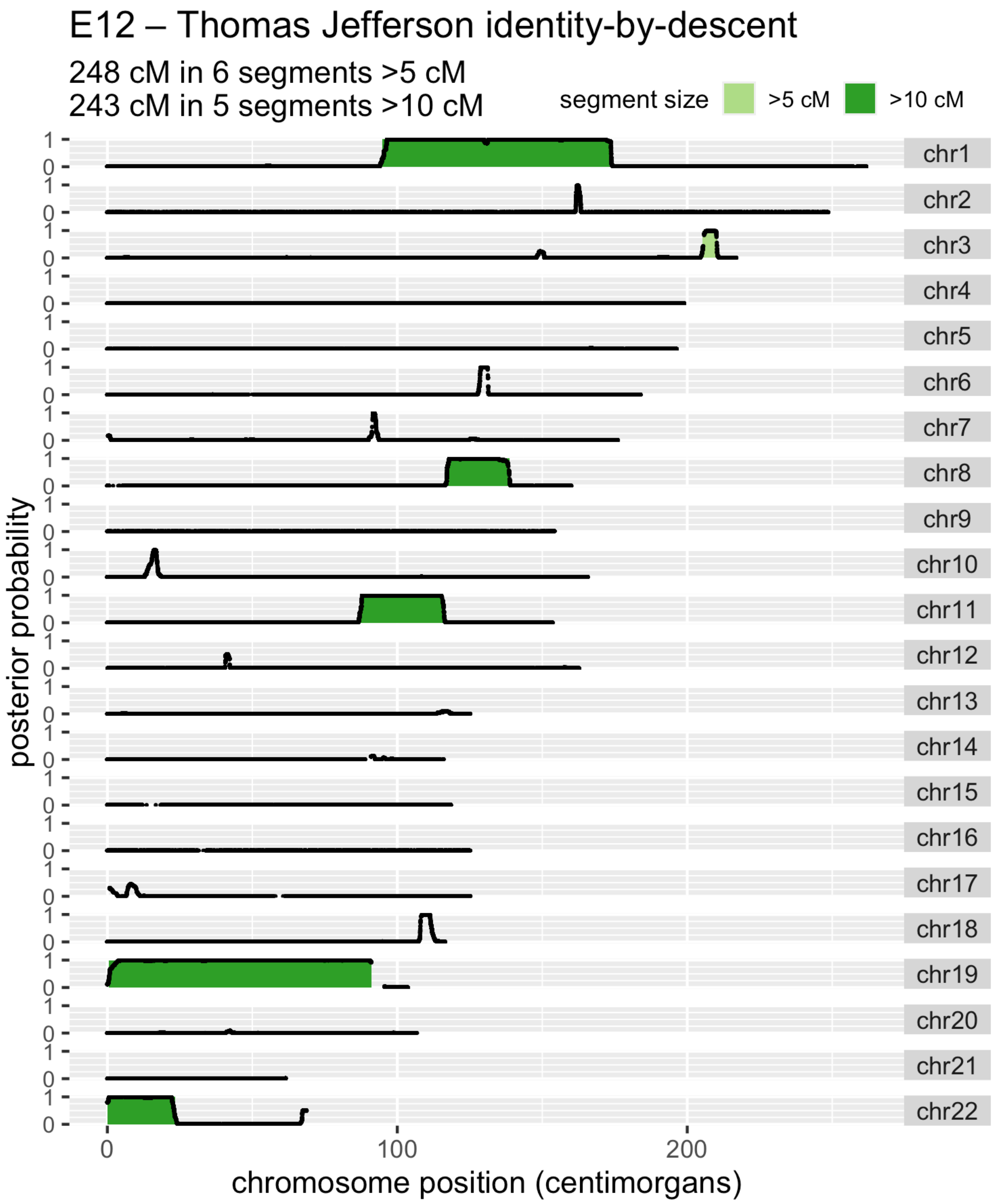

**Figure S12 – TJBD scan against individual E12.** E12 is a known descendant of Eston Hemings. The posterior probabilities of IBD for each SNP are shown as black points. Detected IBD segments are shaded in light green if greater than 5 centimorgans in size and dark green if greater than 10 centimorgans in size.

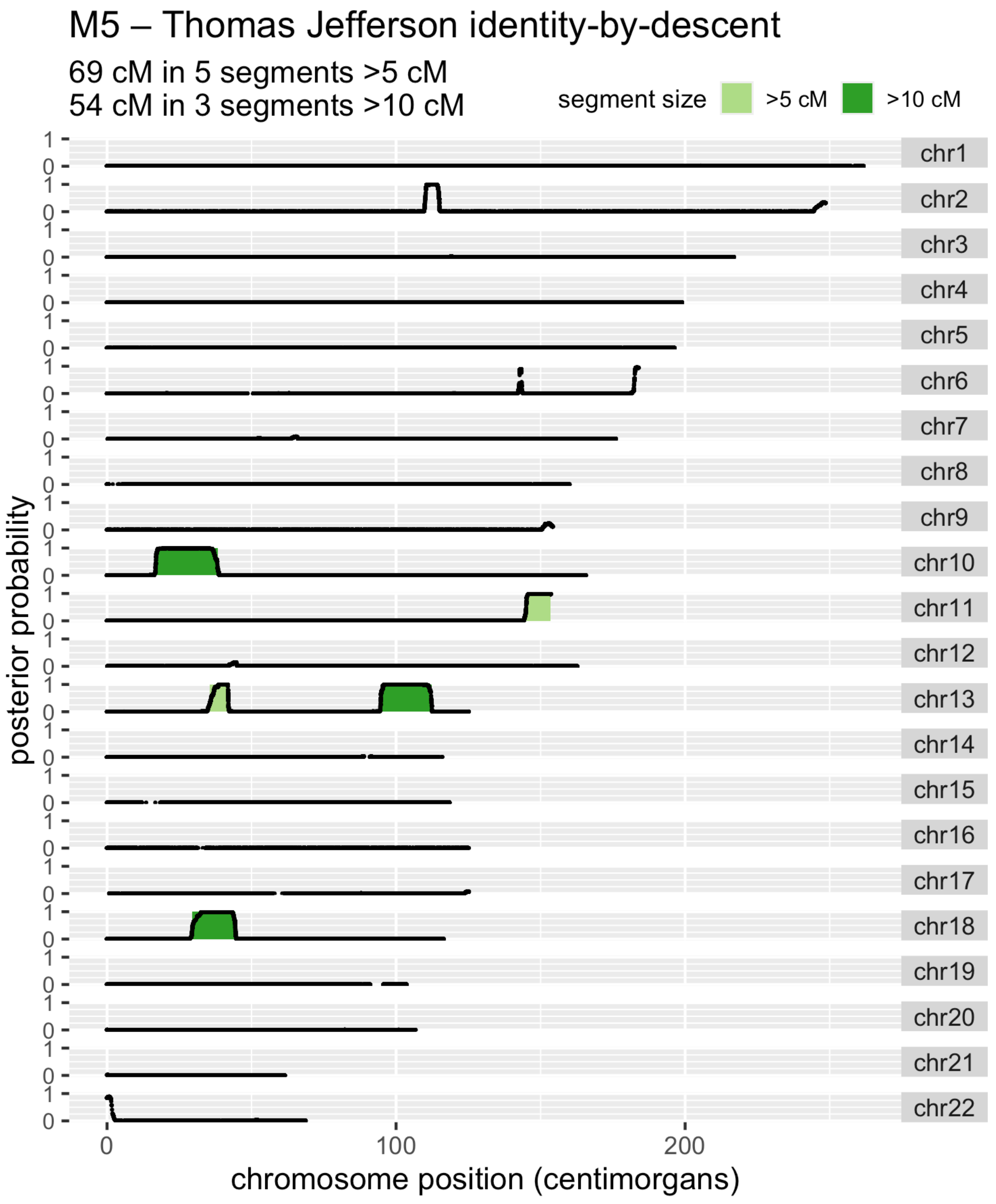

**Figure S13 – TJBD scan against individual M5.** M5 is a known descendant of Madison Hemings. The posterior probabilities of IBD for each SNP are shown as black points. Detected IBD segments are shaded in light green if greater than 5 centimorgans in size and dark green if greater than 10 centimorgans in size.

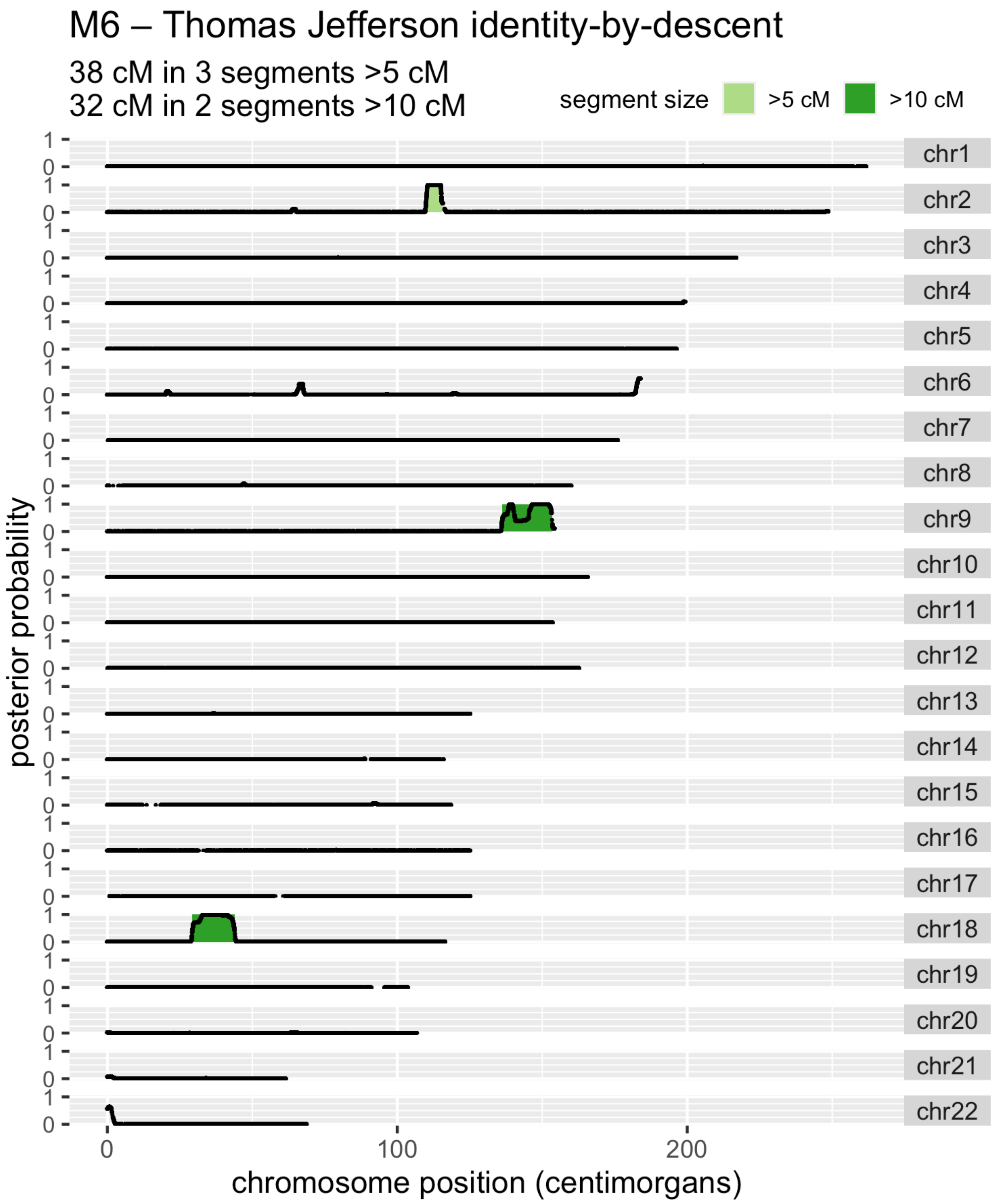

**Figure S14 - TJBD scan against individual M6.** M6 is a known descendant of Madison Hemings. The posterior probabilities of IBD for each SNP are shown as black points. Detected IBD segments are shaded in light green if greater than 5 centimorgans in size and dark green if greater than 10 centimorgans in size.

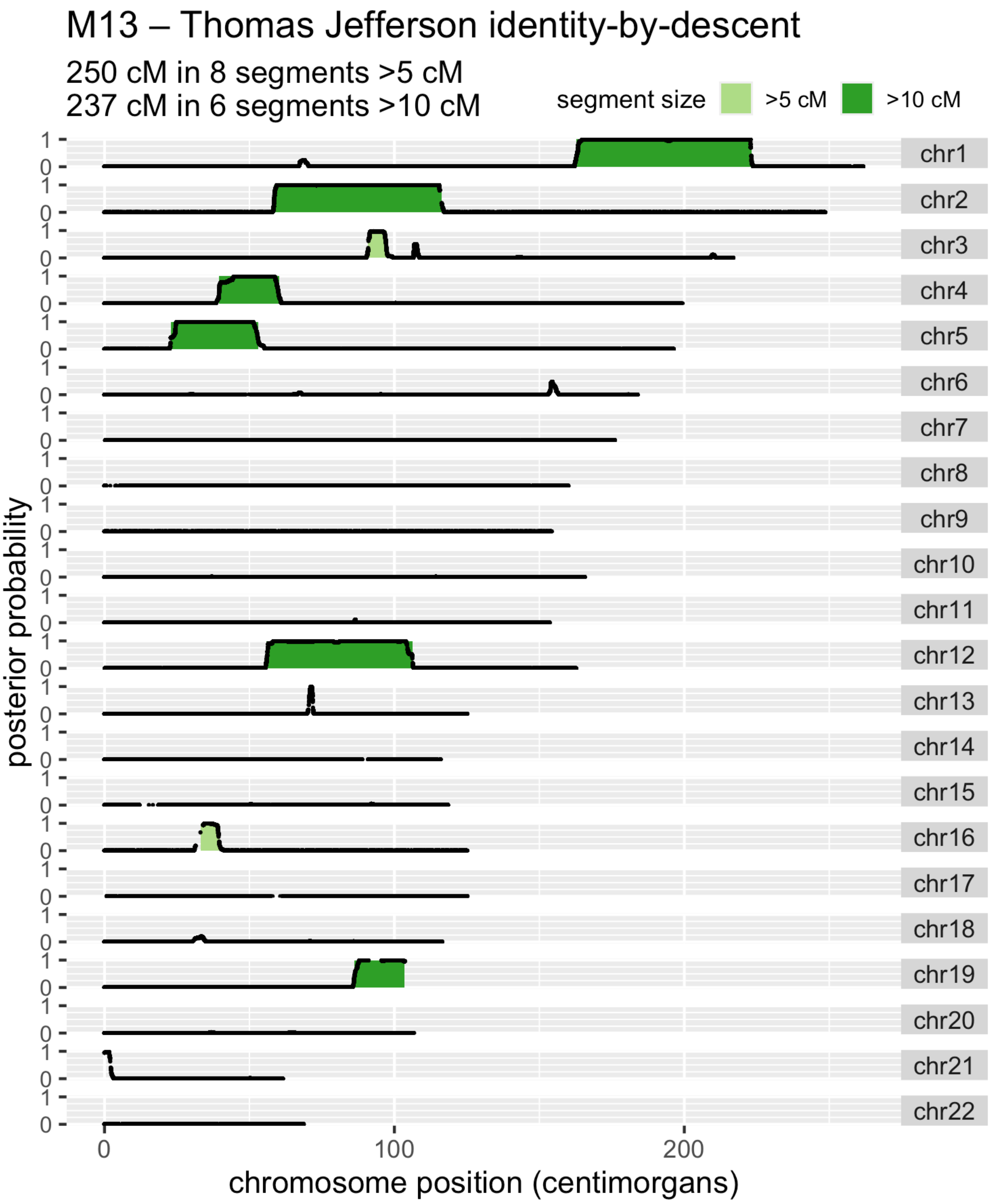

**Figure S15 - TJBD scan against individual M13.** M13 is a known descendant of Madison Hemings. The posterior probabilities of IBD for each SNP are shown as black points. Detected IBD segments are shaded in light green if greater than 5 centimorgans in size and dark green if greater than 10 centimorgans in size.

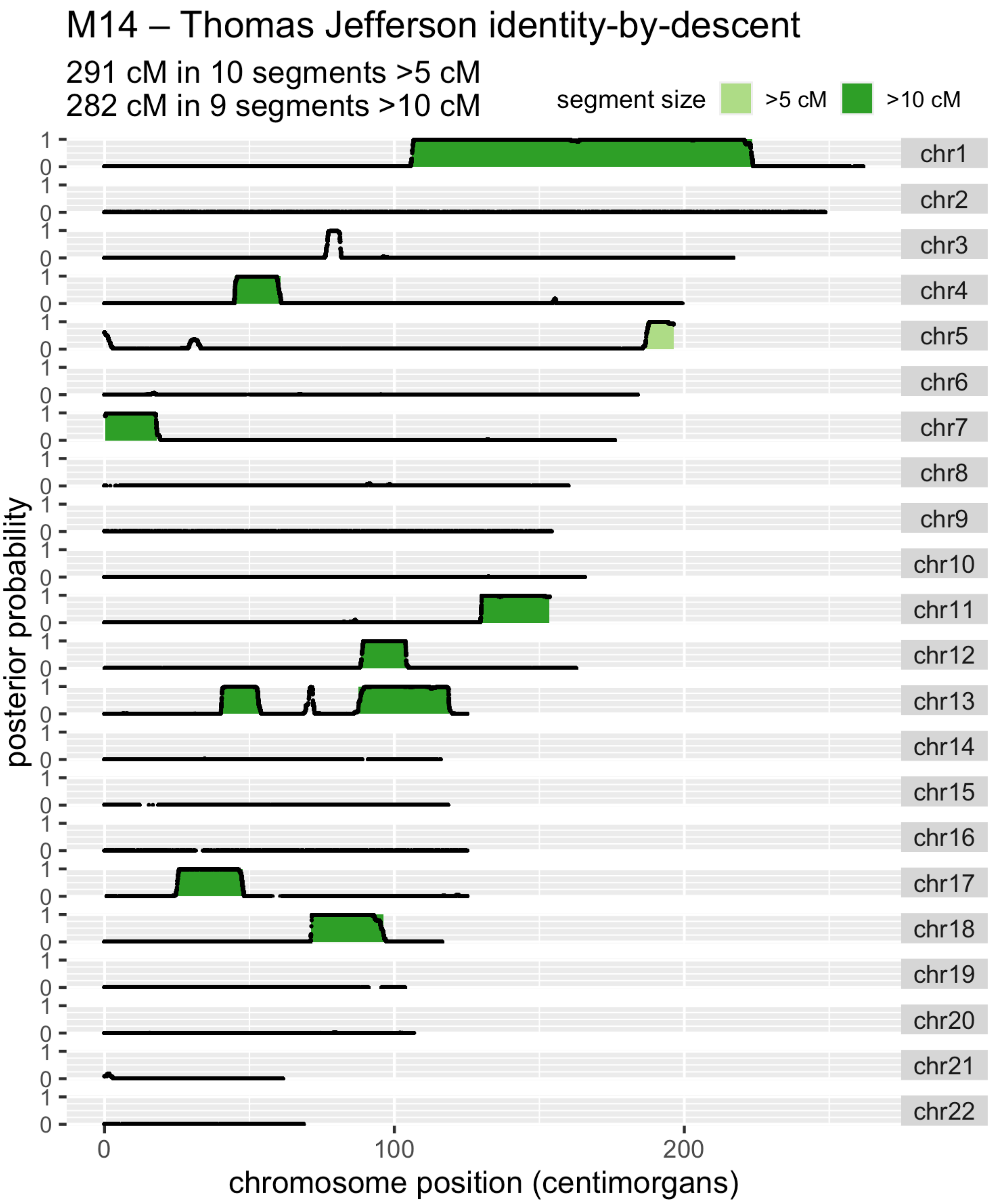

**Figure S15 - TJBD scan against individual M14.** M14 is a known descendant of Madison Hemings. The posterior probabilities of IBD for each SNP are shown as black points. Detected IBD segments are shaded in light green if greater than 5 centimorgans in size and dark green if greater than 10 centimorgans in size.

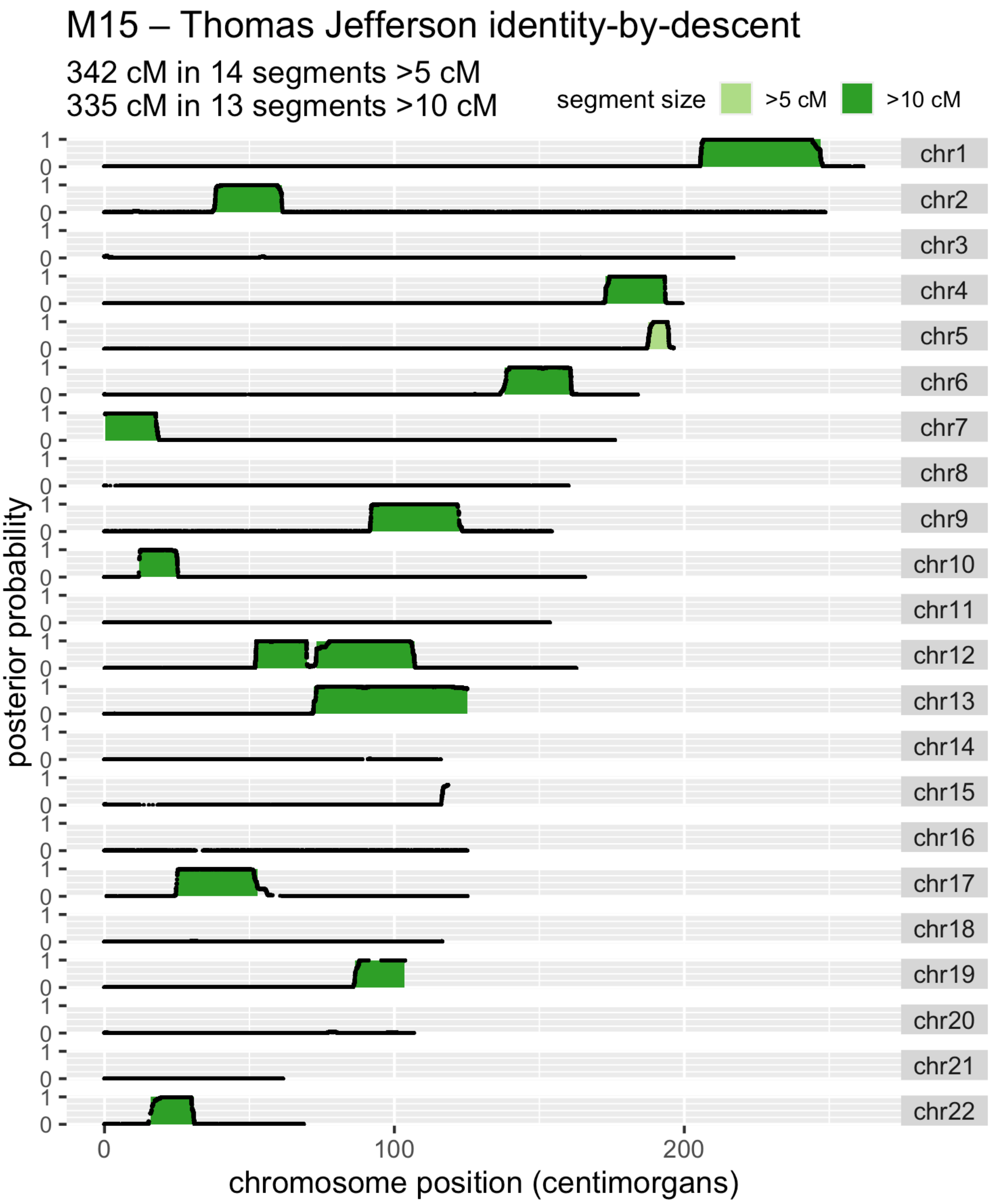

**Figure S17 - TJBD scan against individual M15.** M15 is a known descendant of Madison Hemings. The posterior probabilities of IBD for each SNP are shown as black points. Detected IBD segments are shaded in light green if greater than 5 centimorgans in size and dark green if greater than 10 centimorgans in size.

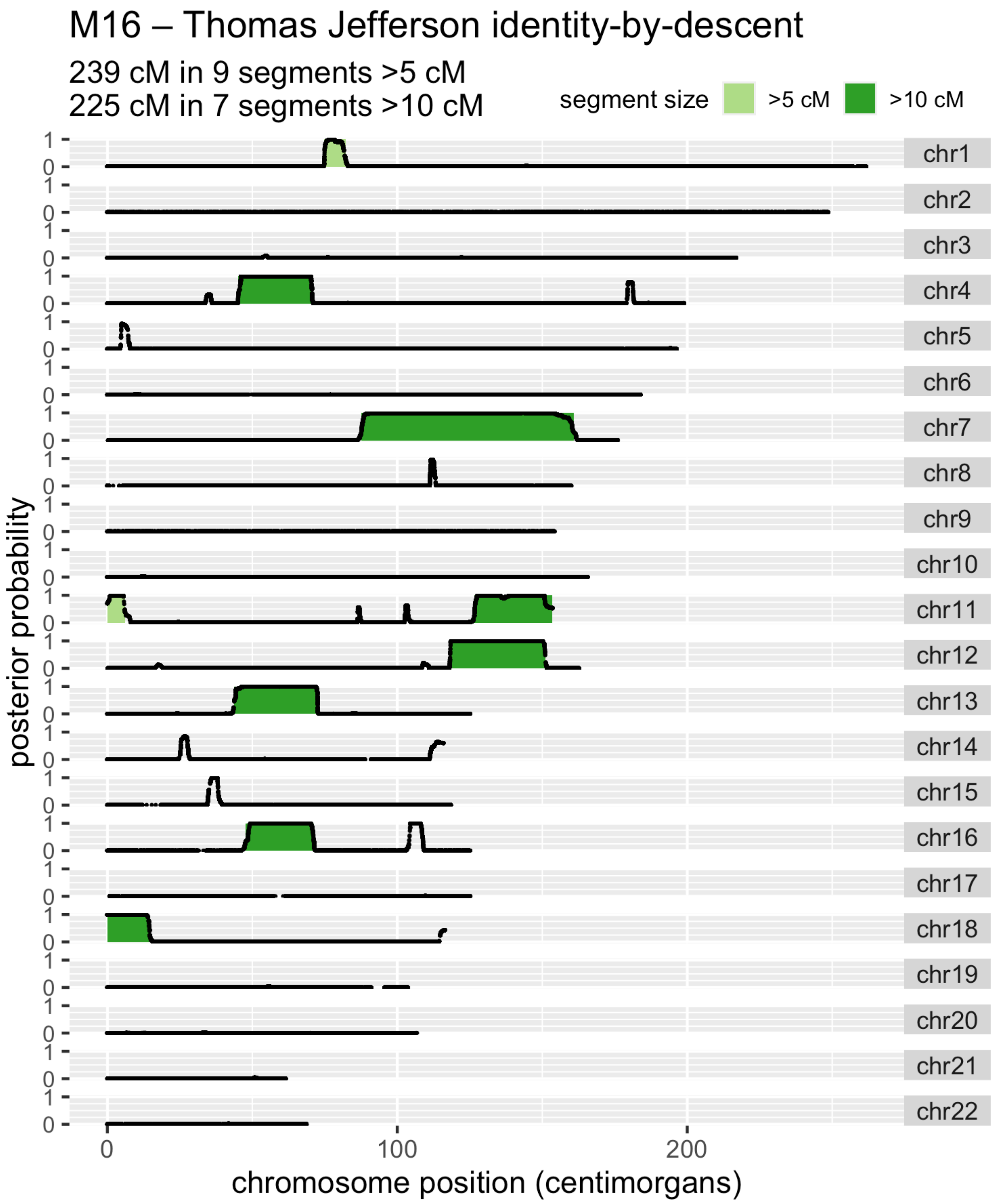

**Figure S18- TJBD scan against individual M16.** M16 is a known descendant of Madison Hemings. The posterior probabilities of IBD for each SNP are shown as black points. Detected IBD segments are shaded in light green if greater than 5 centimorgans in size and dark green if greater than 10 centimorgans in size.

#### Controls

Since our scan does not consider the ancestral composition of the descendant individuals, we used controls to evaluate the effect of ancestry on false positive segments. We ran the TJBD scan with the same parameters described above with no minimum segment size requirement on the individuals from the British in England and Scotland (GBR) and the African Ancestry in Southwest US (ASW) populations from 1000 Genomes Project. Since it is unlikely that any of these individuals share recent ancestry with Thomas Jefferson, any detected segments are likely due to population background matching or other false positives. For the GBR population we found an average of 3.04 segments per individual with a mean size of 2.06 centimorgans (Table S2). For ASW individuals, we found segments with a similar mean segment size (2.01 cM), but only an average of 1.08 segments per individual.

**Table S2 – Summary of TJBD scan controls.** Results are summarized across control individuals from the 1000 Genomes Project, British in England and Scotland (GBR) and African Ancestry in Southwest US (ASW) individuals.

| Population | Segment Count | | Segment Size (cM) | | Total IBD (cM) | |
|---|---|---|---|---|---|---|
| | mean | std. dev | mean | std. dev | mean | std. dev |
| ASW (N=74) | 1.08 | 1.09 | 2.01 | 1.08 | 2.17 | 2.39 |
| GBR (N=91) | 3.04 | 1.84 | 2.06 | 1.69 | 6.28 | 4.98 |

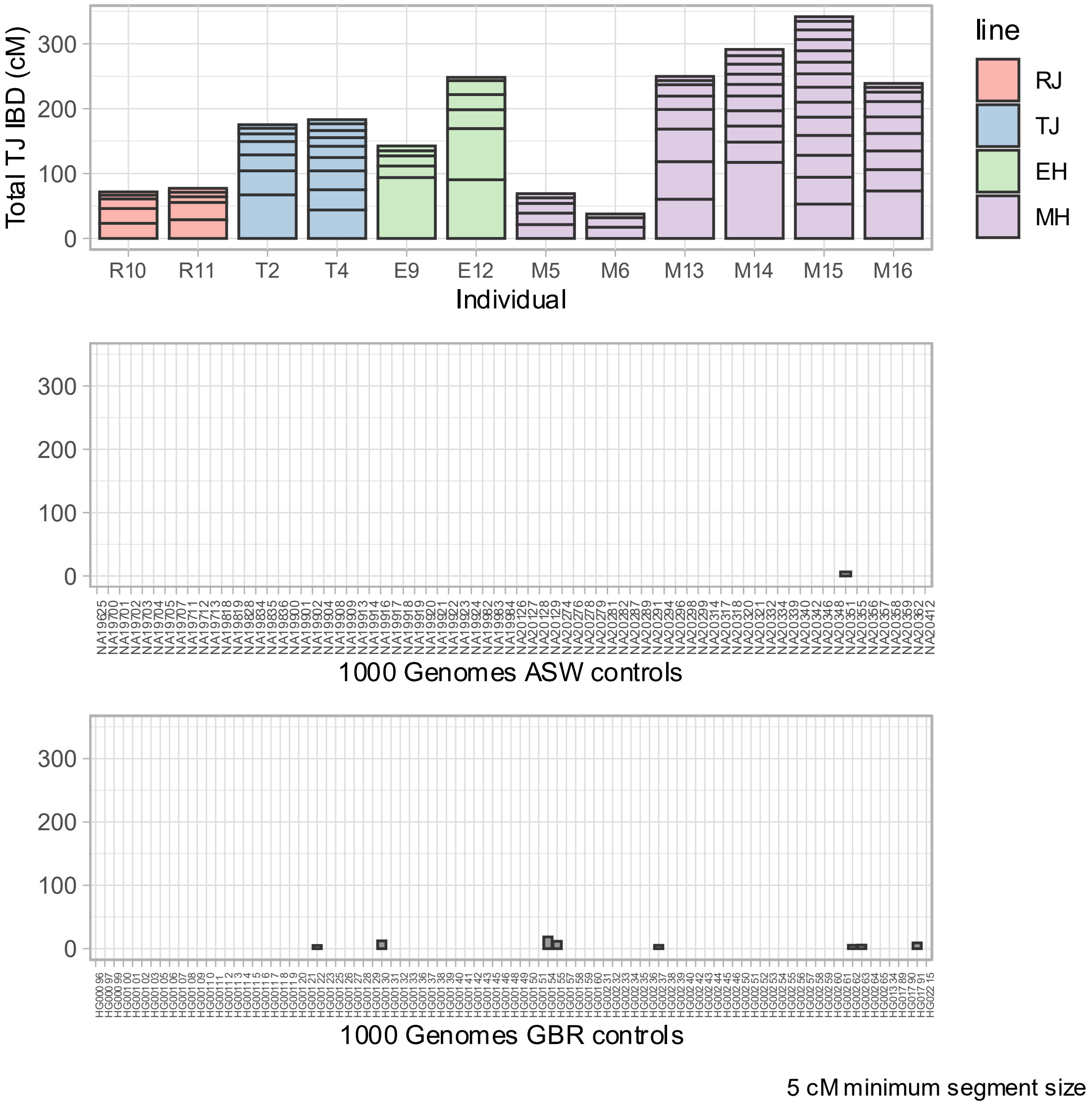

**Figure S19– TJBD control scans**. Comparison of TJ IBD scan results for descendants of Thomas Jefferson and Martha Wayles (T2 and T4), Randolph Jefferson and Anne Lewis (R10 and R11), Eston Hemings (E9 and E12) and Madison Hemings (M5, M6, M13, M14, M15, and M16) with unrelated individuals from the 1000 Genomes Project. Each panel displays the detected TJ IBD segments shown as stacked bars. **Top:** TJ IBD segments greater than 5 cM found in the panel of descendants. **Middle:** TJ IBD segments detected in individuals from the African Ancestry in Southwest US (ASW) subpopulation from the 1000 Genomes Project. **Bottom:** TJ IBD segments detected in individuals from British in England and Scotland (GBR) subpopulation from the 1000 Genomes Project.

#### Scan Replicates & Alternative Reference Panel

We examined the impact of non-deterministic behavior during data preparation for the HMM scan and the choice of reference panel on the detected IBD segments. The HMM only considers only one observed base per marker site from the low-coverage sample but sites may be overlapped by multiple reads. In these instances, a single read is sampled at random. Consequently, scans with different starting pseudo-random seed values may have differences in the reported IBD segments. The choice of reference panel may also have a significant effect on the detected IBD segments. To evaluate these effects, we ran the HMM scan 4 additional times with different random seeds using the GBR reference panel and 3 times using the reference panel based on the 1000 Genomes Utah residents (CEPH) with Northern and Western European ancestry (CEU) population. We then examined the detected segments in the Hemings descendants, grouped and merged according to Hemings ancestor (Eston or Madisons) as these observations are the basis for our statistical analysis.

We found the IBD segments detected by scans with different random seeds and using the GBR and CEU reference panels to be largely concordant (Table S3, Figures S20 and S21). For scans using the GBR reference panel, the most common differences were on the boundaries of reported segments which caused segments to appear slightly longer or shorter between scans. This pattern matches what would be expected given the HMM's one low-coverage observation per site limitation (i.e., edges of IBD segments may be missed by sampling from the non-IBD chromosome phase). We also observed occasional drop out between scans in segments between 5 and 10 centimorgans in size. With larger segments, we observed some instances of segments in one scan being detected as two shorter segments with a small gap in between. We found a similar pattern of differences when using the CEU panel and a slightly wider range of variation between scans. We note that the parameters used to extract IBD segments were tuned based on the GBR reference panel and using parameters specific to the CEU panel may improve consistency. We also observed more consistency between scans in the EH branch compared to the MH scans. The wider variation in the MH results may be explained by a larger number of "degrees of freedom", i.e., the MH branch has more individuals and more segments that can vary between scans compared to the EH branch.

The results of all 8 scans are included in Supplemental Data Table S1: IBD Scan Results.

**Table S3** - Summary of detected IBD segments from scans using GBR and CEU reference panels after merging segments by Hemings branch.

| | | | IBD >5 cM | | IBD >10 cM | |
|---|---|---|---|---|---|---|
| **Branch** | **Reference Panel** | **Replicate** | **Number of Segments** | **Total IBD cM** | **Number of Segments** | **Total IBD cM** |
| EH | GBR | 1 | 9 | 294.3 | 6 | 274.0 |
| EH | GBR | 2 | 9 | 294.5 | 6 | 274.3 |
| EH | GBR | 3 | 9 | 295.8 | 6 | 274.1 |
| EH | GBR | 4 | 9 | 295.4 | 6 | 274.0 |
| EH | GBR | 5 | 9 | 295.2 | 6 | 273.7 |
| EH | CEU | 1 | 10 | 281.6 | 6 | 253.4 |
| EH | CEU | 2 | 9 | 293.5 | 6 | 274.5 |
| EH | CEU | 3 | 8 | 282.0 | 6 | 270.9 |
| MH | GBR | 1 | 27 | 868.1 | 22 | 827.4 |
| MH | GBR | 2 | 28 | 866.7 | 23 | 826.7 |
| MH | GBR | 3 | 27 | 870.6 | 22 | 831.1 |
| MH | GBR | 4 | 26 | 857.7 | 22 | 823.8 |
| MH | GBR | 5 | 28 | 879.5 | 22 | 833.8 |
| MH | CEU | 1 | 29 | 873.5 | 24 | 829.3 |
| MH | CEU | 2 | 28 | 873.8 | 23 | 829 |
| MH | CEU | 3 | 30 | 888.3 | 22 | 821.7 |

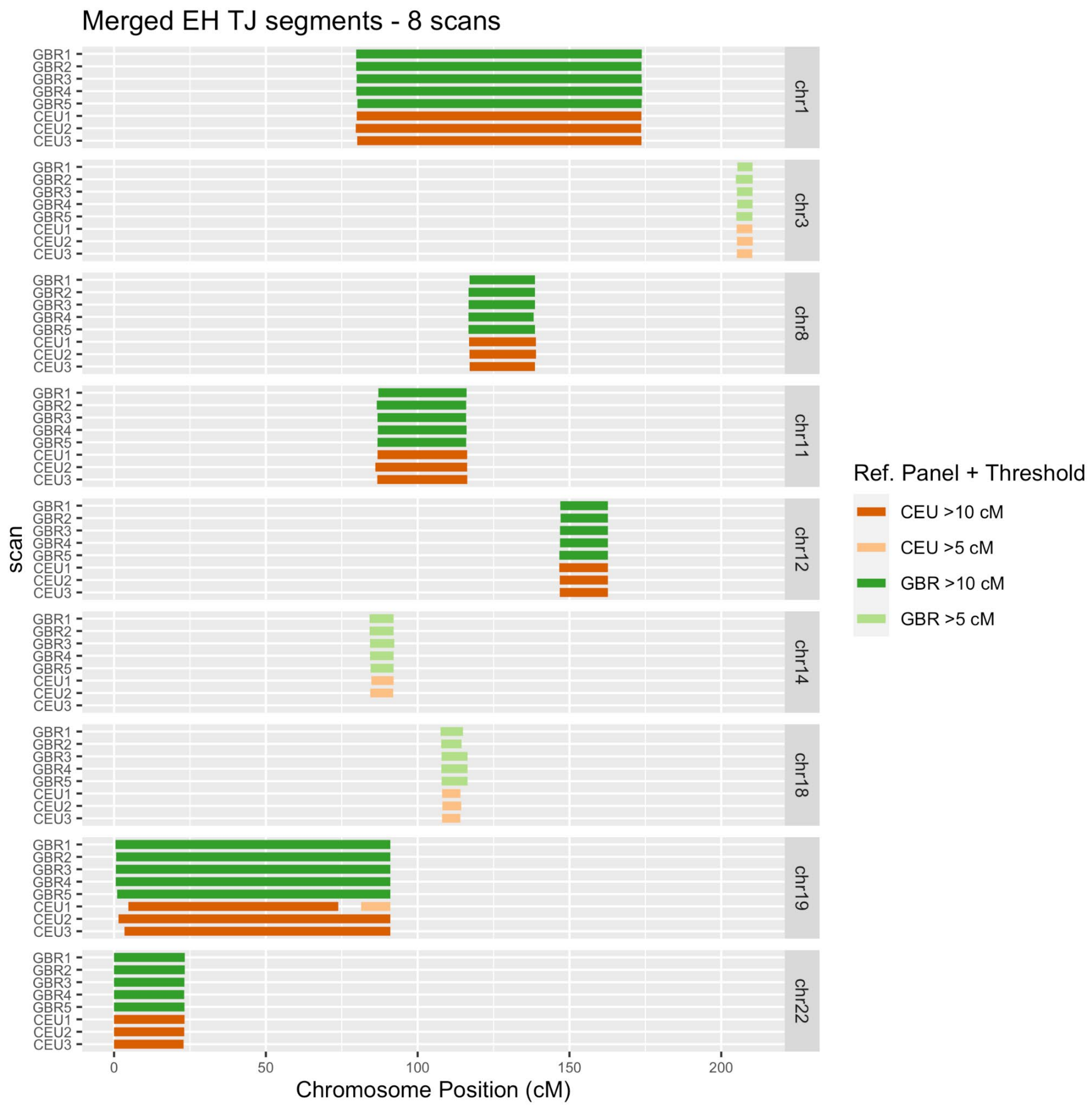

**Figure S20 -** IBD segments detected in descendants of Eston Hemings (EH), E9 and E12, in 8 independent scans. Scans labelled GBR1-5 use the reference panel based on the 1000 Genomes GBR population and scans CEU1-3 use the reference panel based on the CEU population. Only chromosomes with detected segments in any of the scans are shown.

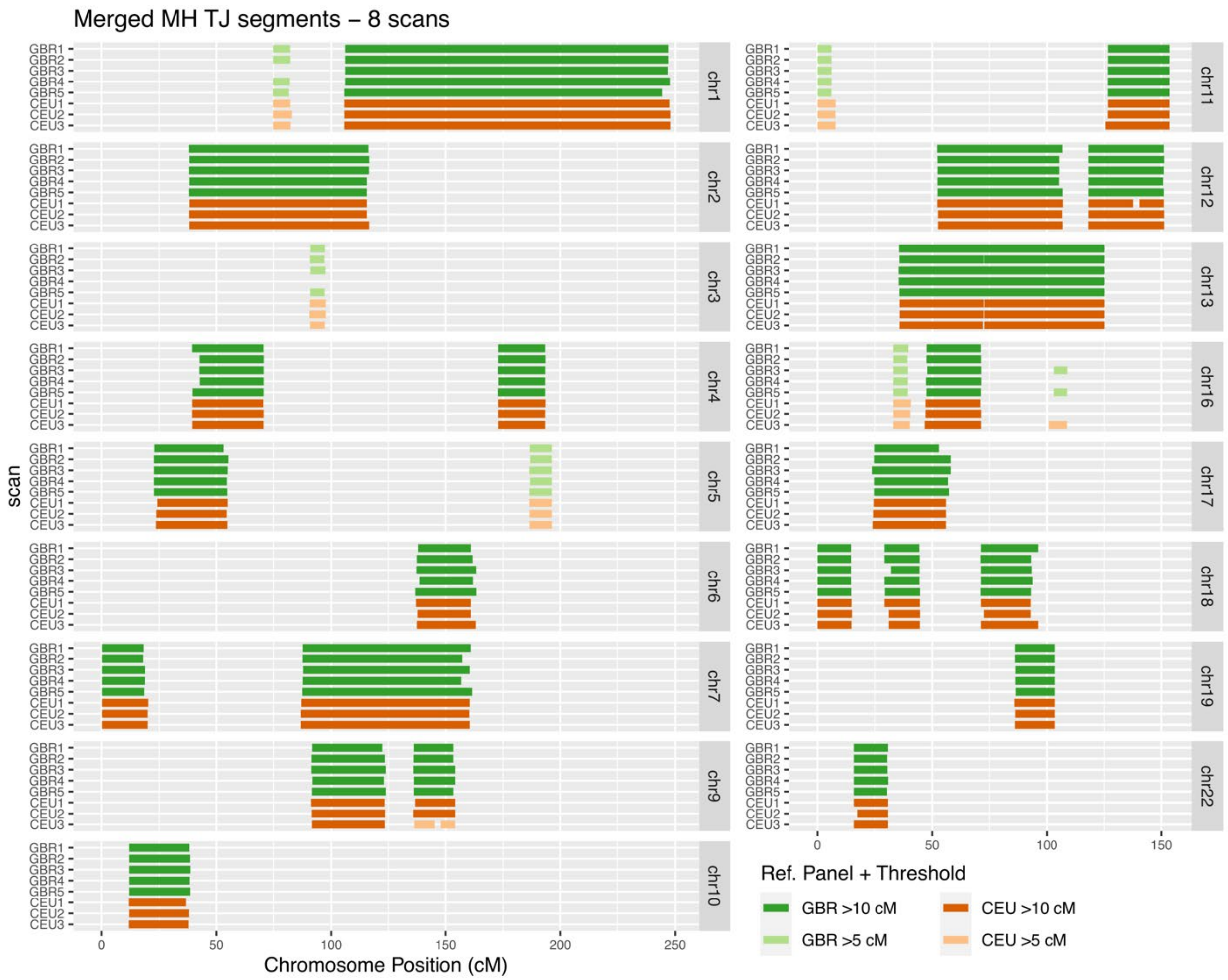

**Figure S21** - TJ IBD segments detected in descendants of Madison Hemings (MH) (M5, M6, M13, M14, M15, and M16) in 8 independent scans. Scans labelled GBR1-5 use the reference panel based on the 1000 Genomes GBR population and scans CEU1-3 use the reference panel based on the CEU population. Only chromosomes with detected segments in any of the scans are shown.

#### Statistical evaluation of Jefferson paternity scenarios

##### Genealogy Simulations

We used ped-sim (28) to simulate the 3 paternity scenarios for Madison and Eston Hemings and the subsequent generations of recombination leading to their present-day descendants as well as the genealogies of the other Jefferson descendants. We chose to include sex-specific recombination in the simulations as several lineages in the genealogy contain unequal numbers of male and female ancestors. Lower recombination rates in males compared to females tend to produce longer IBD segments, but fewer in number, and increase variance in the total shared IBD (28). We specified the sex of every member of the genealogy and used the sex-specific high-resolution genetic map from Bhérer et al. (33) and interference parameters from Campbell et al. (34) to model recombination. We produced 24,000 simulation replicates for each scenario and performed the same grouping and merging of IBD segments described above. We split the simulation results for each scenario into two sets; a 20,000 replicate “training” set that was used to fit the parameters of the likelihood function and a 4,000 replicate “test” set that was used to evaluate the likelihood results. We also produced an additional 1,000 simulation replicates for the genealogy of the descendants of Thomas Jefferson and Martha Wayles (T2 and T4) and the genealogy of the descendants of Randolph Jefferson and Anne Lewis (R10 and R11).

#### Simulated Genealogies

##### Descendants of Thomas Jefferson and Martha Wayles

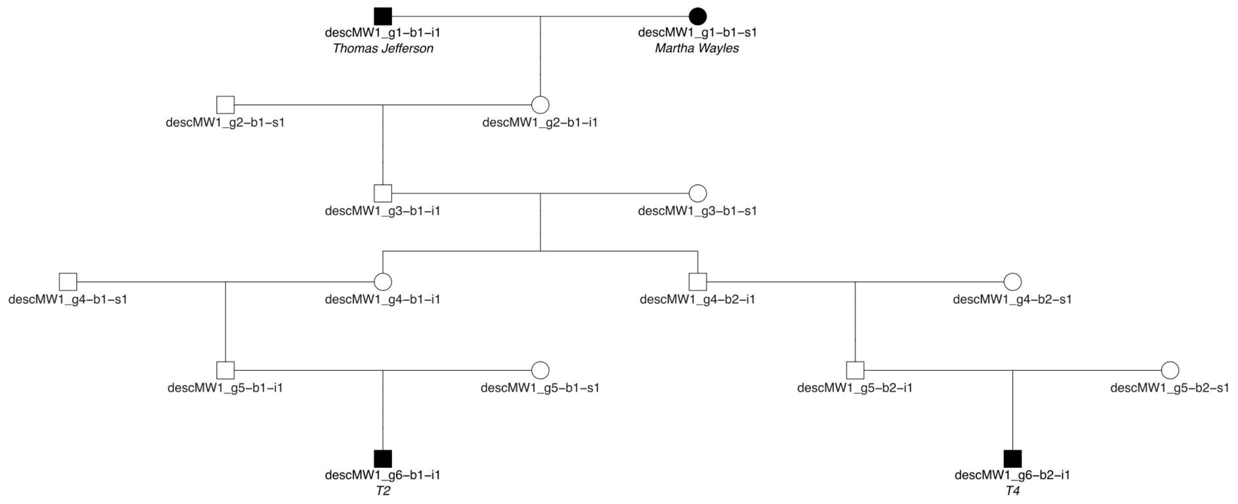

Example ped-sim def file input:

```
def descMW 1000 6
1 1 1 1sM
2 0 1 1sF
3 0 1 1sM
4 0 2 1sF 2sM
5 0 2 1:1 2:2 1,2sM
6 1 2 1:1 2:2 1,2sM
```

**Figure S22– Simulation framework for T2 and T4.** Pedigree structure (top) and Ped-sim definition input (bottom) used in genealogy simulations for T2 and T4, direct descendants of Thomas Jefferson and Martha Wayles Jefferson. Each node is labeled with the identifier from ped-sim. Names of historical individuals and identifiers of sampled descendants are shown in italics.

#### Descendants of Randolph Jefferson and Anne Lewis

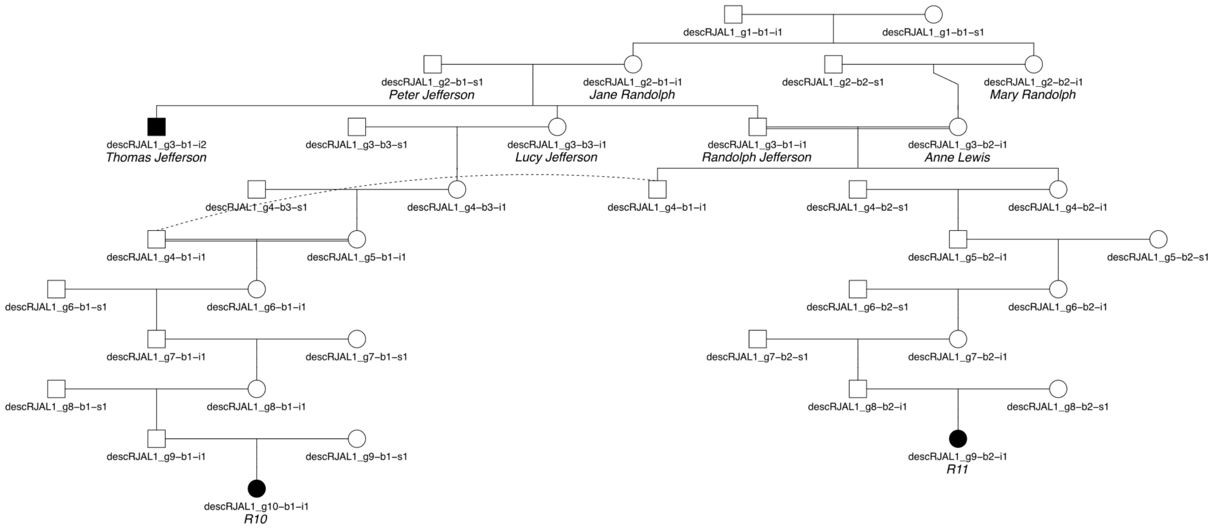

Example ped-sim def file input:

```
def descRJAL 1000 10
2 0 2 1,2sF
3 2 3 1,3:1 1sM 2sF 3sF 2,3n
4 0 3 1,2:1_2 1sM 2,3sF
5 0 2 2:2 1:3 1sF 2sM
6 0 2 1:1_1^4 1sF 2sF
7 0 2 1sM 2sF
8 0 2 1sF 2sM
9 1 2 1sM 2sF 1n
10 1 1 1sF
```

**Figure S23 – Simulation framework for R10 and R11**. Pedigree structure (top) and Ped-sim definition input (bottom) used in genealogy simulations for R10 and R11, direct descendants of Randolph Jefferson (brother of Thomas Jefferson) and Anne Lewis (first cousin of Thomas Jefferson). Each node is labeled with the identifier from ped-sim. Names of historical individuals and identifiers of sampled descendants are shown in italics.

#### Descendants of Sally Hemings - TJ Paternity Scenario

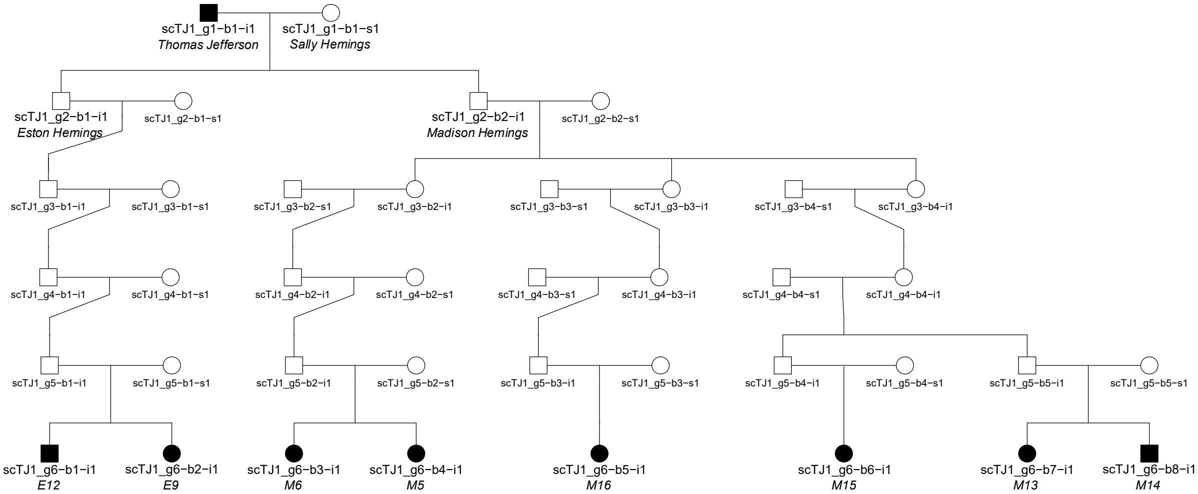

Example ped-sim def file input:

```
def scTJ 4000 6
1 1 1 1sM
2 0 2 1-2sM
3 0 4 1:1 2-4:2 1sM 2-4sF
4 0 4 1,2sM 3,4sF
5 0 5 1:1 2:2 3:3 4,5:4 1-5sM
6 1 8 1,2:1 3,4:2 5:3 6:4 7-8:5 1,8sM 2-7sF
```

**Figure S24 – Simulation framework for descendants of Sally Hemings – Thomas Jefferson paternity scenario.** Pedigree structure (top) and Ped-sim definition input (bottom) used in genealogy simulations for Eston Hemings descendants (E9 and E12) and Madison Hemings descendants (M5, M6, M13, M14, M15, and M16) for the scenario where Thomas Jefferson is the father of both Eston and Madison Hemings (TJ scenario). Each node is labeled with the identifier from ped-sim. Names of historical individuals and identifiers of sampled descendants are shown in italics.

#### Descendants of Sally Hemings - RJ Paternity Scenario

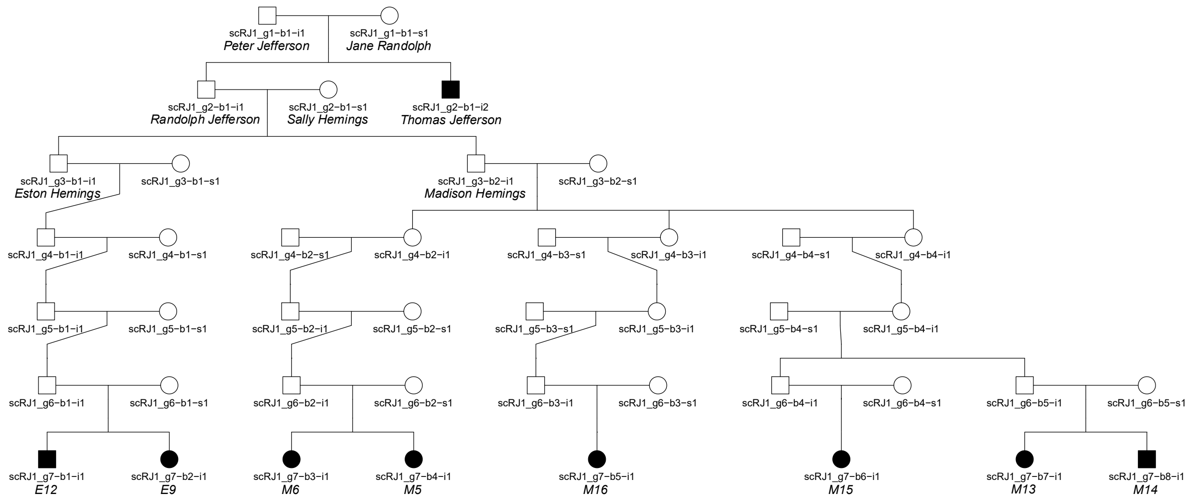

Example ped-sim def file input:

```
def scRJ 4000 7
1 0 1
2 2 1 1sM
3 0 2 1-2sM
4 0 4 1:1 2-4:2 1sM 2-4sF
5 0 4 1,2sM 3,4sF
6 0 5 1:1 2:2 3:3 4,5:4 1-5sM
7 1 8 1,2:1 3,4:2 5:3 6:4 7-8:5 1,8sM 2-7sF
```

**Figure S25- Simulation framework for descendants of Sally Hemings - Randolph Jefferson paternity scenario.** Pedigree structure (top) and Ped-sim definition input (bottom) used in genealogy simulations for Eston Hemings descendants (E9 and E12) and Madison Hemings descendants (M5, M6, M13, M14, M15, and M16) for the scenario where Randolph Jefferson is the father of both Eston and Madison Hemings (RJ scenario). Each node is labeled with the identifier from ped-sim. Names of historical individuals and identifiers of sampled descendants are shown in italics.

#### Descendants of Sally Hemings - RJO Paternity Scenario

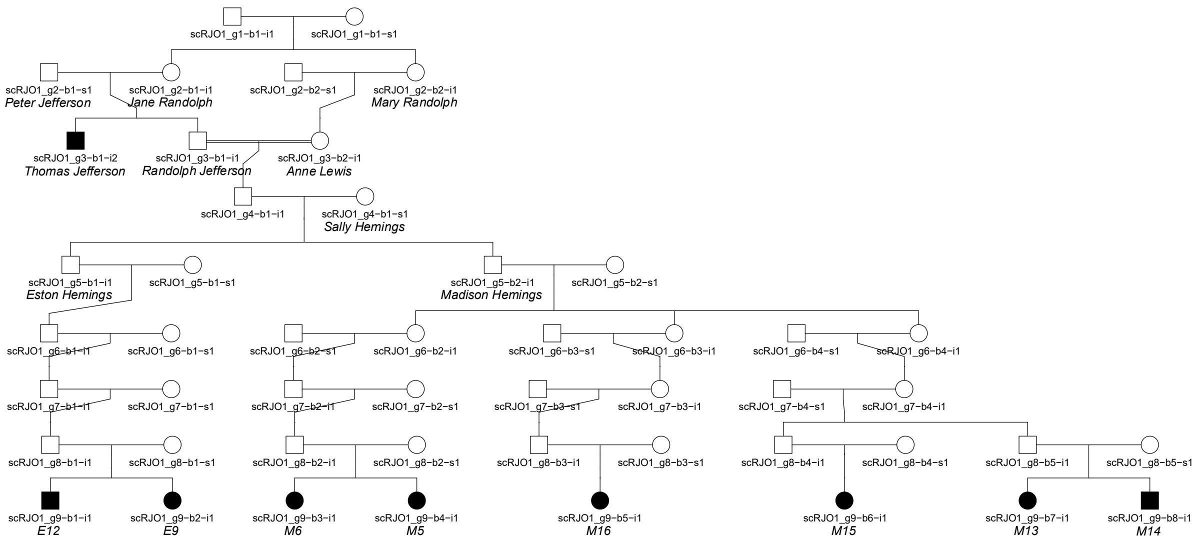

Example ped-sim def file input:

```
def scRJO 4000 9
2 0 2 1,2sF
3 2 2 1sM 2sF 2n
4 0 1 1:1_2 1sM
5 0 2 1-2sM
6 0 4 1:1 2-4:2 1sM 2-4sF
7 0 4 1,2sM 3,4sF
8 0 5 1:1 2:2 3:3 4,5:4 1-5sM
9 1 8 1,2:1 3,4:2 5:3 6:4 7-8:5 1,8sM 2-7sF
```

**FigureS26– Simulation framework for Sally Hemings descendants – Randolph Jefferson offspring paternity scenario.** Pedigree structure (top) and Ped-sim definition input (bottom) used in genealogy simulations for Eston Hemings descendants (E9 and E12) and Madison Hemings descendants (M5, M6, M13, M14, M15, and M16) for the scenario where a son of Randolph Jefferson and Anne Lewis is the father of both Eston and Madison Hemings (RJO scenario). Each node is labeled with the identifier from ped-sim. Names of historical individuals and identifiers of sampled descendants are shown in italics.

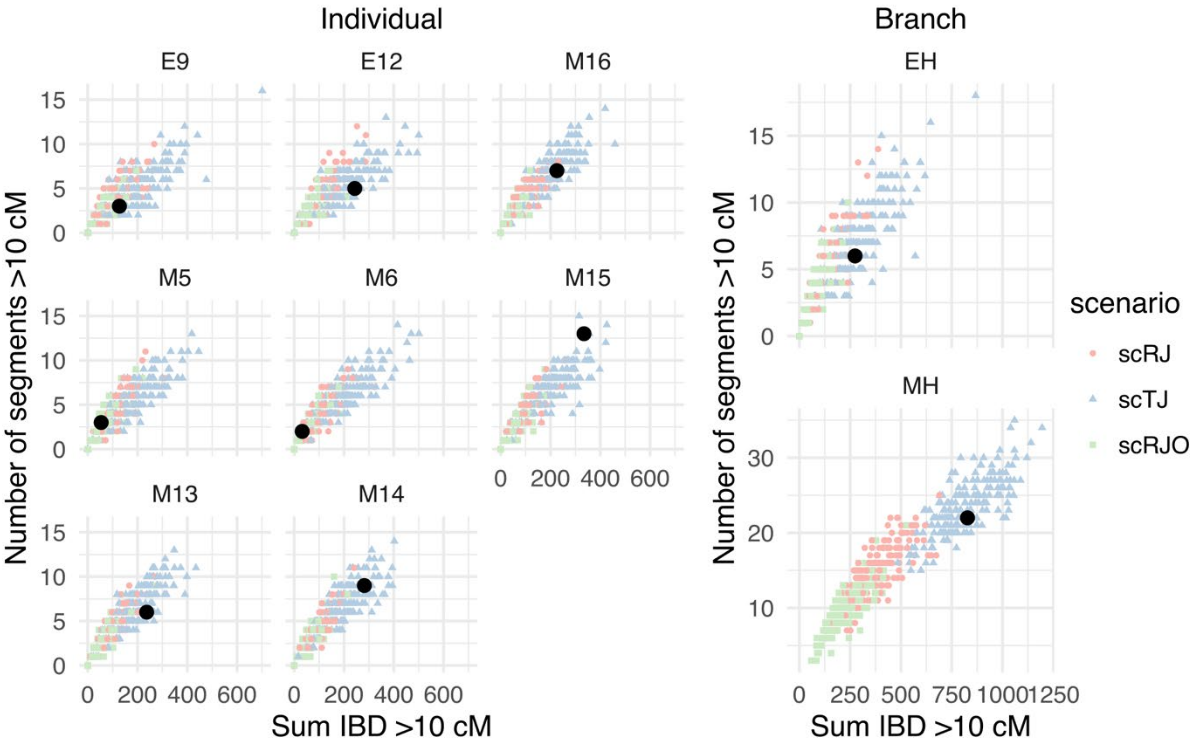

**Figure S27 – Comparison of paternity scenario simulations. Left:** Comparison of observed number and total size of IBD segments greater than 10 cM in each Hemings descendant (black circles) with 200 simulations of each paternity scenario (TJ, RJ, and RJO) (shapes in color). When considering the expected Thomas Jefferson IBD in a single individual, the distributions for each scenario overlap considerably. **Right:** Comparison of observed number and total size of IBD segments greater than 10 cM merged by Hemings ancestor (Eston Hemings (EH) and Madison Hemings (MH)) (black circles) with 200 simulations of each paternity scenario (TJ, RJ, and RJO) (shapes in color). Pooling Thomas Jefferson IBD segments found in each descendant by Hemings ancestor improves the separability of the scenario distributions.

#### Likelihood Model

To estimate the likelihood of our IBD observations under the 3 proposed paternity scenarios, we used a likelihood function like the one proposed by Huff et al. (35) to detect and estimate relatedness between pairs of individuals. Let $n$ equal the number of observed merged match segments found among the sampled descendants of a Hemings offspring and $s_{1..n}$ the sizes of the merged segments. For a paternity scenario $p \in \{TJ, RJ, RJO\}$, genealogy $g \in \{EH, MH\}$, and minimum size threshold $t$ , the likelihood is:

$$L(n, s|p, g, t) = N(n|p, g, t) \cdot S(s|p, g, t)$$

$$S(s|p, g, t) = \prod_{i \in s} F(s_i|p, g, t)$$

Where $N(n|p, g, t)$ is the probability of observing $n$ segments longer than $t$ centimorgans, $S(s|p, g, t)$ is the probability of the set of segment sizes and $F(s_i|p, g, t)$ is the probability of observing a segment of length $s_i$. We use a Poisson distribution (standard or generalized) with mean equal to sample mean segment count from simulations to approximate $N(n|p, g, t)$. For

$F(s_i|p,g,t)$, we use an exponential distribution using the mean segment size of segments greater than threshold $t$ from the simulations ($\theta_{p,g,t}$).

$$F(s_i|p,g,t) = \frac{e^{-(s_i-t)/\theta_{p,g,t}}}{\theta_{p,g,t}}$$

The likelihood includes an assumption that the number of segments and the sizes of the segments are independent. We observed a weak inverse relationship between the number and mean size of merged segments in the simulated data. But, we note that the variation within the simulated scenarios is small when compared to the differences between scenarios (Figure S29).

The original ERSA methodology was designed to distinguish between related and unrelated individuals in a large sample and considers matches due to the population background in addition to recent shared ancestry. For our purposes, we assume all matches are due to recent ancestry and use the minimum size threshold to reduce the effect of background matches.

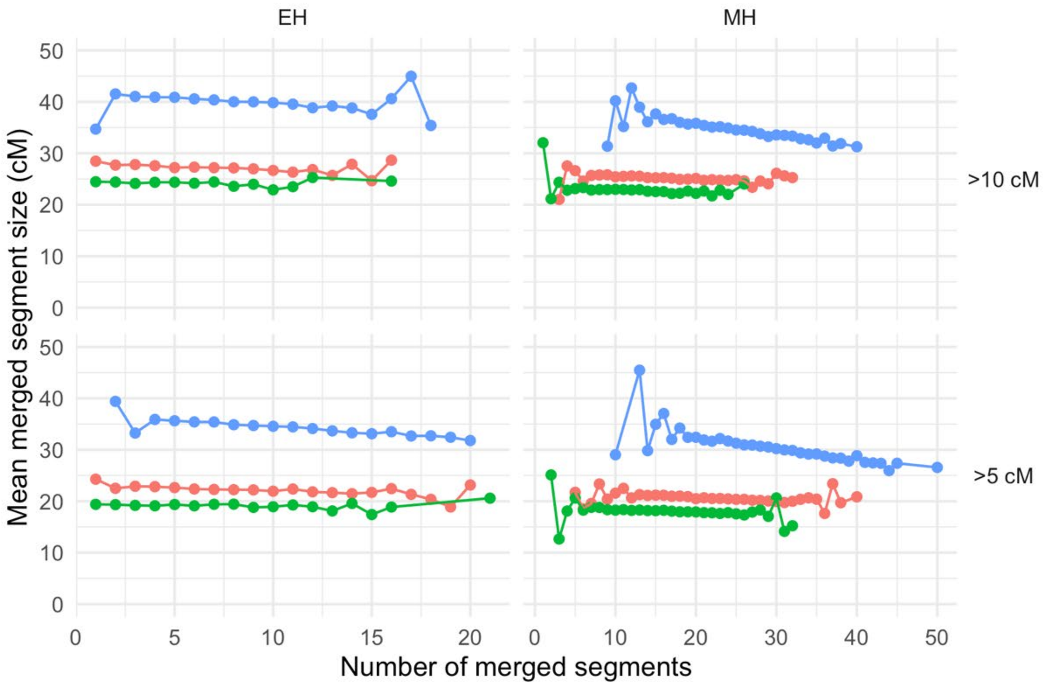

**Figure S28** - Relationship between number of merged segments and mean merged segment size for each genealogy and paternity scenario combination.

We fit the parameters of the Poisson and exponential distributions for each paternity scenario model using the `fitdistplus` R library (10.18637/jss.v064.i04) and evaluated the results using histograms and quantile-quantile plots (Figures S29 and S30). First, we confirmed that the fit parameters matched the sample means of the simulated data (Table S4). For the Poisson distributions, we found that the theoretical density and cumulative distribution functions matched the empirical observations for the paternity scenarios. We noted tendencies in the cases of the TJ scenario and/or the MH genealogy for the empirical distributions to be under dispersed when compared to the theoretical expectation (Figure S29). These differences may be the result of these cases being more affected by the merging of overlapping segments due to the density of matches in the TJ scenario or the number of redundant matches in the larger MH genealogy. To address the underdispersion, we fit a more flexible generalized Poisson distribution using the *statsmodels* python package (Seabold, Skipper, and Josef Perktold. “statsmodels: Econometric and statistical modeling with python.” Proceedings of the 9th Python in Science Conference. 2010). We found the generalized Poisson model provided a better fit for the observed data (Figure S30, Table S5) but note that similar results were obtained using the standard Poisson and would lead to the same conclusions (Table S6). For the exponential distributions we found a general “light tail” pattern where we observe fewer very long segments (>~100 cM) in the empirical distribution compared with what is expected by the theoretical distribution (Figure S30). This pattern can be explained by the fact that the size of match segments are constrained by the sizes of individual chromosomes. We also note that only 1 observed merged segment exceeds 100 cM in size.

| | | | Number of Segments | | Threshold adjusted segment size (cM) | |
|---|---|---|---|---|---|---|
| Branch | Threshold (cM) | Scenario | Mean | Poisson fit | Mean | Exponential fit |
| EH | 5 | TJ | 8.90460 | 8.90460 | 29.58323 | 29.58323 |
| EH | 5 | RJ | 6.77245 | 6.77245 | 17.28486 | 17.28486 |
| EH | 5 | RJO | 4.83875 | 4.83875 | 14.21017 | 14.21017 |
| EH | 10 | TJ | 7.40220 | 7.40220 | 30.09390 | 30.09390 |
| EH | 10 | RJ | 5.07305 | 5.07305 | 17.27346 | 17.27346 |
| EH | 10 | RJO | 3.39040 | 3.39040 | 14.26314 | 14.26314 |
| MH | 5 | TJ | 28.63000 | 28.63000 | 25.37804 | 25.37804 |
| MH | 5 | RJ | 20.49915 | 20.49915 | 15.63307 | 15.63307 |
| MH | 5 | RJO | 14.38955 | 14.38955 | 13.12571 | 13.12571 |
| MH | 10 | TJ | 24.05635 | 24.05635 | 24.66381 | 24.66381 |
| MH | 10 | RJ | 15.17955 | 15.17955 | 15.20786 | 15.20786 |
| MH | 10 | RJO | 9.94430 | 9.94430 | 12.87228 | 12.87228 |

**Table S4** - Parameters of the ERSA likelihood’s Poisson and exponential distributions used to model Thomas Jefferson IBD sharing under the 3 paternity scenarios for each branch and minimum segment size threshold combination. The mean number of segments and mean segment size adjusted by the threshold size from the simulations used to fit the parameters are included for comparison.

| | | | Number of Segments Generalized Poisson Parameters | |
|---|---|---|---|---|
| Branch | Threshold (cM) | Scenario | Mean/location | Dispersion/shape |
| EH | 5 | TJ | 8.90460 | -0.10318 |
| EH | 5 | RJ | 6.77245 | 0.05274 |
| EH | 5 | RJO | 4.83875 | 0.06546 |
| EH | 10 | TJ | 7.40220 | -0.0968 |
| EH | 10 | RJ | 5.07305 | 0.03132 |
| EH | 10 | RJO | 3.39040 | 0.0496 |
| MH | 5 | TJ | 28.63000 | -0.21596 |
| MH | 5 | RJ | 20.49915 | -0.06443 |
| MH | 5 | RJO | 14.38955 | -0.00659 |
| MH | 10 | TJ | 24.05635 | -0.21415 |
| MH | 10 | RJ | 15.17955 | -0.06851 |
| MH | 10 | RJO | 9.94430 | -0.01732 |

**Table S5** - Parameters of the ERSA likelihood's generalized Poisson distribution used to model Thomas Jefferson IBD sharing under the 3 paternity scenarios for each branch and minimum segment size threshold combination.

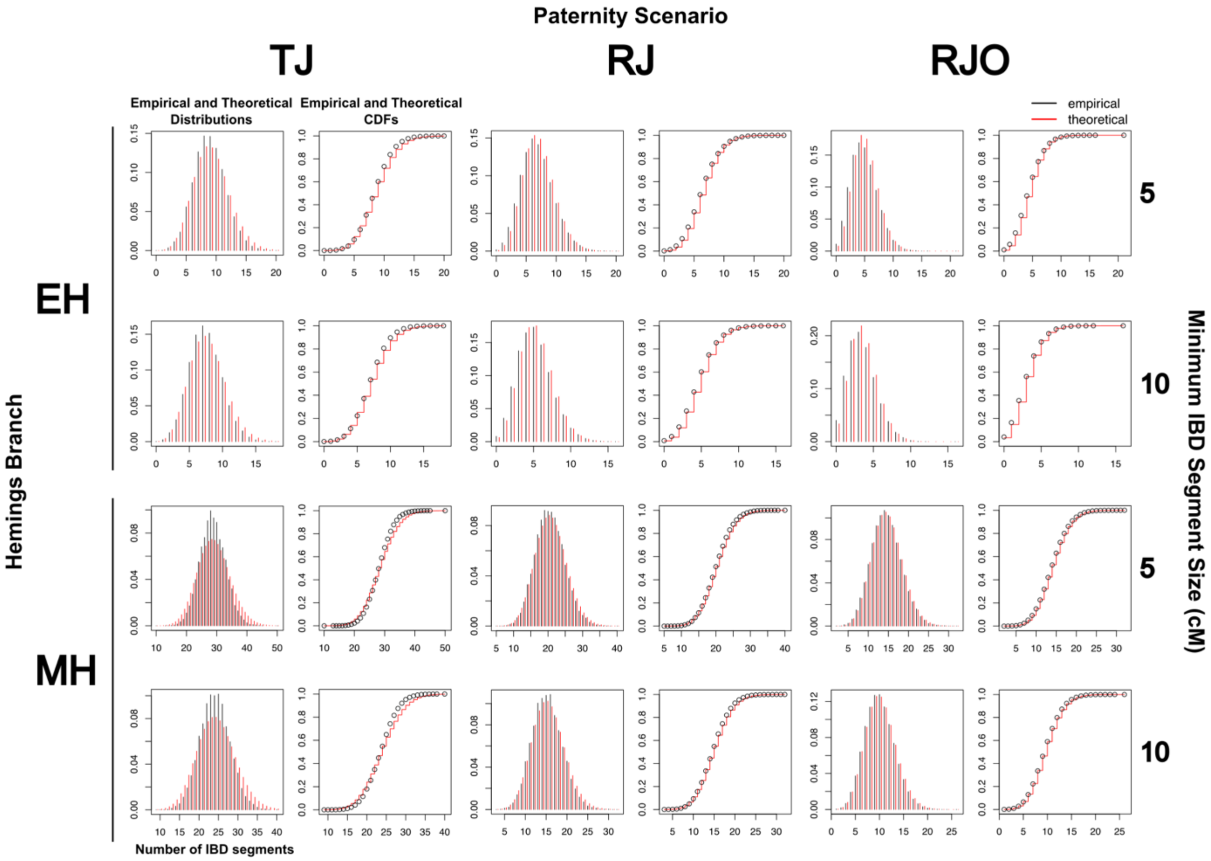

**Figure S29**- Comparisons of the number of merged segments per simulation with the Poisson approximation using the mean number of observed segments for each combination of paternity scenario (TJ, RJ, or RJO), Hemings branch (EH or MH), and minimum IBD segment size (5 cM or 10 cM). A histogram comparing the empirical and theoretical distributions and a plot of the empirical and theoretical cumulative distribution functions (CDF) are shown for each model.

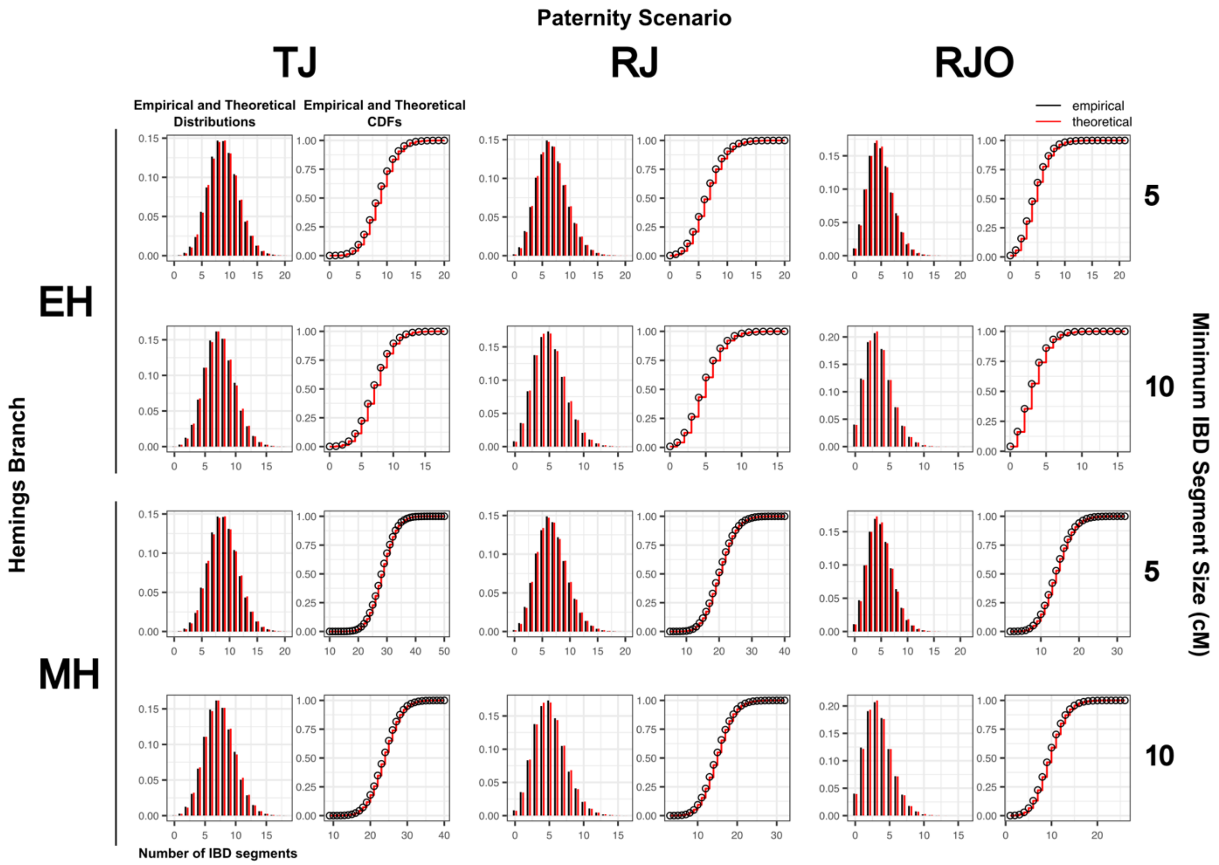

**Figure S30**- Comparisons of the number of merged segments per simulation with the generalized Poisson approximation for each combination of paternity scenario (TJ, RJ, or RJO), Hemings branch (EH or MH), and minimum IBD segment size (5 cM or 10 cM). A histogram comparing the empirical and theoretical distributions and a plot of the empirical and theoretical cumulative distribution functions (CDF) are shown for each model.

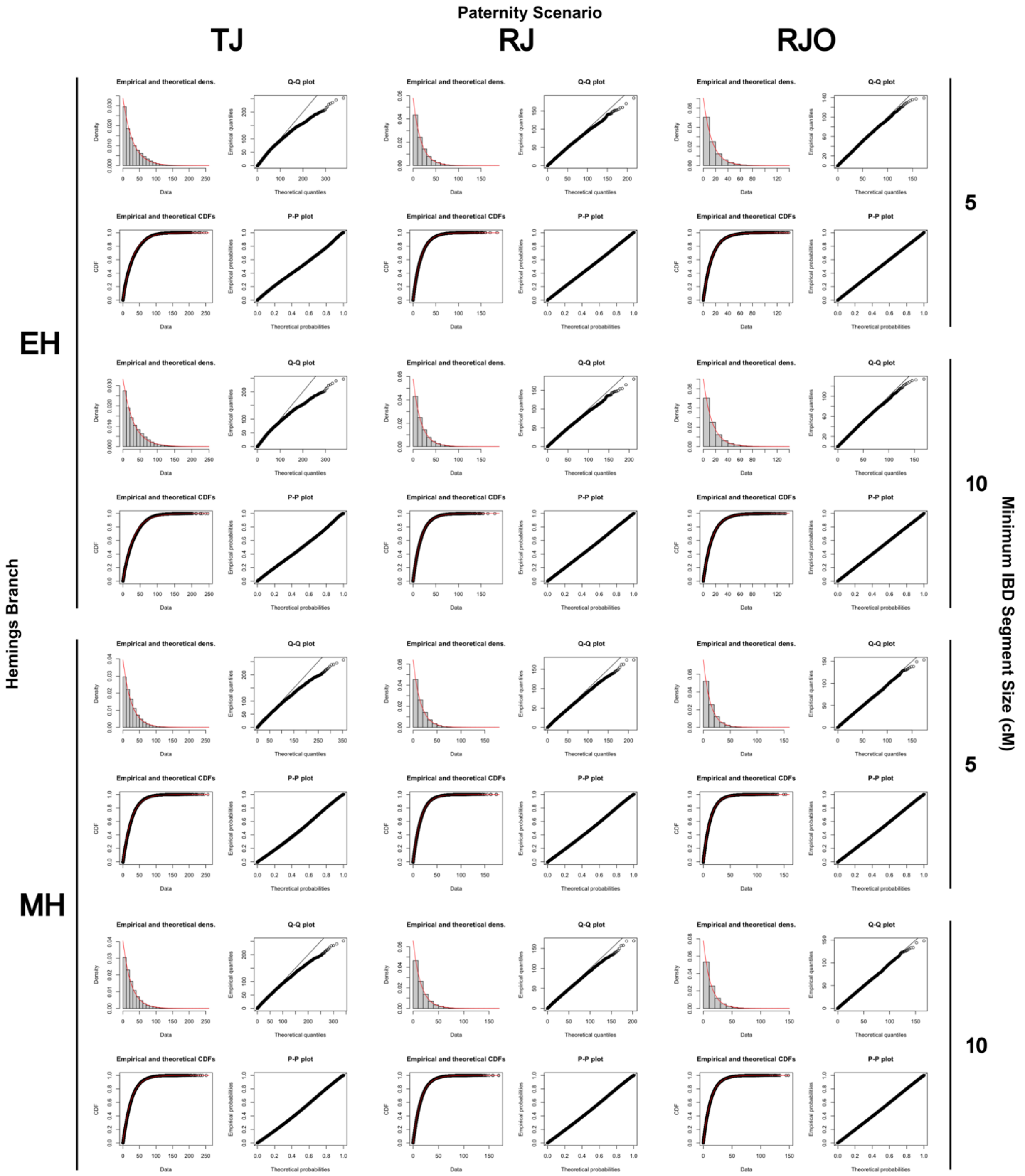

**Figure S31**- Comparisons of the distribution of merged segment sizes with the exponential approximation using the mean merged segment size for each combination of paternity scenario (TJ, RJ, or RJO), Hemings branch (EH or MH), and minimum IBD segment size (5 cM or 10 cM). There are four sub-plots for each model: a histogram comparing the empirical and theoretical distributions (top left), a quantile-quantile plot (top right), a comparison of the empirical and theoretical cumulative distribution functions (CDF) (bottom right), and a probability-probability plot (bottom right).

##### Likelihood Ratios

We computed likelihood ratios (or Bayes factors) from the likelihoods of the observed TJ matches under the 3 paternity scenario models to compare the likelihood of each scenario for each branch and IBD threshold. For paternity scenarios $p_1, p_2 \in \{TJ, RJ, RJO\}$, the likelihood ratio is computed as:

$$LR_{p_1:p_2} = L(n, s|p_1, g, t) \; / \; L(n, s|p_2, g, t)$$

##### Posterior Probabilities

We computed the posterior probabilities of the 3 paternity scenario models given the observed matches, assuming equal prior probabilities for the scenarios. We applied Bayes' theorem to the likelihood described earlier:

$$P(p|n, s, g, t) \; = \frac{L(n, s|\; p = TJ, g, t)P(p)}{P(n, s|g, t)}$$

$$P(n, s|g, t) = \sum_{p \in TJ, RJ, RJO} L(n, s|p, g, t)P(p)$$

Where $P(p \in \{TJ, RJ, RJO\})$ is the prior probability of a paternity scenario.

#### Uncertainty Estimation

A considerable source of uncertainty in our statistical analysis is the variation between IBD scan runs. The IBD HMM only considers one observed base per marker and when more than one sequence overlaps a marker, the base observation is made from one of the sequences drawn at random. Consequently, scans starting from different pseudo-random seeds may result in different detected IBD segments, which may have a significant impact on the final statistics. To examine the variation between scans, we produced 5 independent scans for TJ IBD in the descendant individuals using the GBR reference panel (see previous section). In addition, we devised a resampling strategy that leveraged the fact that chromosome-level results are independent within individuals and individual-level results are similarly independent to explore the range of possible IBD scan results and assess their impact on the final statistics.

We implemented a resampling strategy to construct combinations of scan results from the 5 GBR-based IBD scans (Figure S32). For each descendant individual, we constructed a resampled set of IBD segments by selecting the detected segments from one of the five scans for each chromosome by random draw, producing a combination of independent scan results. We then merged segments by Hemings branch as described above and computed the likelihood under each of the three paternity scenarios with each minimum segment size. We repeated this procedure 2,000 times and obtained the mean, median, and the middle 95% interval for the likelihood ratios and posterior probabilities (See Supplemental Data Table S2: Resampled Statistical Results).

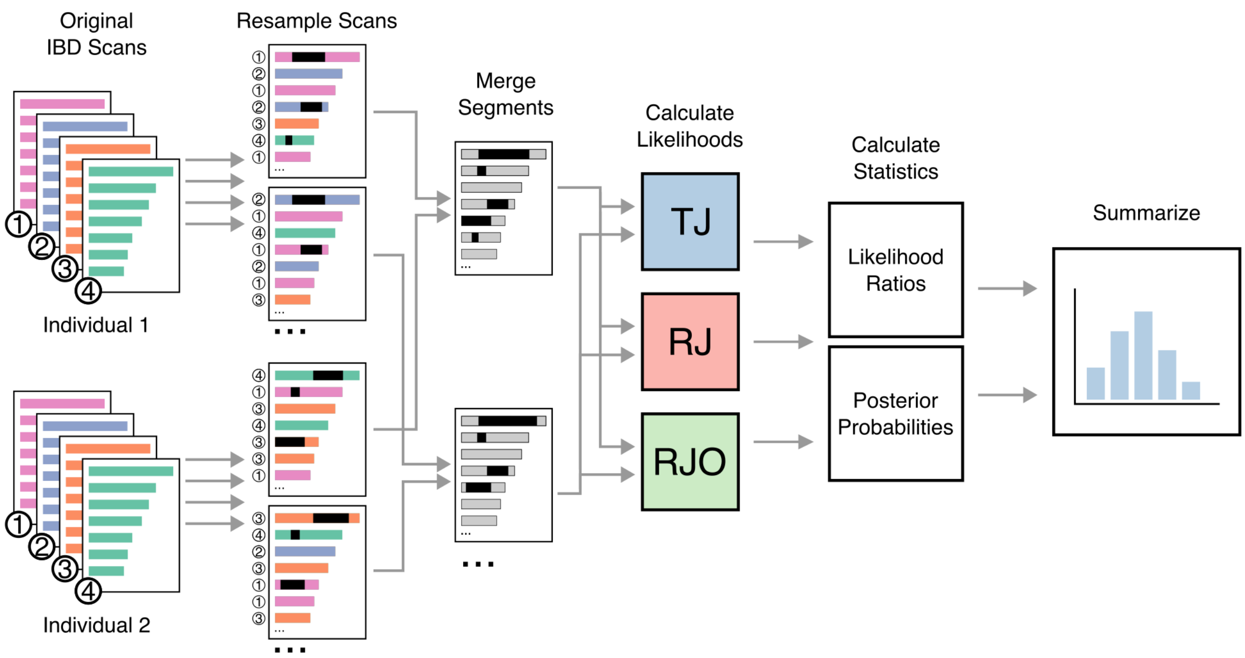

**Figure S32– Schematic of resampling strategy to estimate likelihood ratio and posterior probability uncertainty**. Results from the original IBD scans are sampled by chromosome to produce a new combination of chromosome scans for an individual. Resampled IBD segments from descendants of the same Hemings ancestor are merged and the likelihoods of the resampled segments under the three scenarios (TJ, RJ, and RJO) are computed. Likelihood ratios and posterior probabilities (individual and joint) are calculated from the likelihoods and the results are summarized.

#### Results

##### Individual scans

We computed the likelihoods, likelihood ratios and posterior probabilities for each scan (5 using the GBR reference panel and 3 using the CEU panel) using the likelihood model and parameter estimates described in the previous section (see Tables S4 and S5). The likelihood ratios and posterior probabilities obtained for scans using the GBR reference panel were similar within each branch and minimum segment size threshold. We also found similar results for scans based on the CEU reference panel.

We observed that scans where larger segments were split into two segments had lower likelihood ratios and posterior probabilities supporting the TJ scenario. Specifically, in the Eston Hemings (EH) branch, the first scan based on the CEU reference panel reported two segments on chromosome 19, one smaller than 10 cM, which led to a TJ:RJ likelihood ratio of ~2.2 compared with ~4.5 for the other scans, when considering segments greater than 5 cM. In the Madison Hemings (MH) branch, two segments on chromosome 13 were detected in different individuals as non-overlapping in 1 of the 5 GBR scans and in all 3 CEU scans and were not combined in the merging step. Both results contain potentially important information for the likelihood function. The one segment interpretation captures that it was likely present as a contiguous segment in Madison Hemings, while the two-segment interpretation reflects how they were inherited in descendant individuals. Despite the differences in the segments and statistical results, the conclusions for each scan would be the same. The IBD present in the descendants of Eston Hemings represents “positive” or “substantial” evidence that Thomas Jefferson was the father of Eston Hemings and IBD found in the descendants of Madison Hemings can be interpreted as “very strong” or “decisive” evidence that Thomas Jefferson was the father of Madison Hemings (36).

**Table S6** - Likelihood ratios and posterior probabilities computed assuming Poisson distributed segment counts based on the observed merged TJ matches in the EH and MH descendants under the 3 paternity scenarios for each scan.

| | | | Likelihood Ratio | | | Posterior Probability | | |
|---|---|---|---|---|---|---|---|---|
| **branch** | **threshold** | **scan** | **TJ:RJ** | **TJ:RJO** | **RJ:RJO** | **TJ** | **RJ** | **RJO** |
| EH | 5 | GBR1 | 4.4 | 51.4 | 11.6 | 0.8034 | 0.1810 | 0.0156 |
| EH | 5 | GBR2 | 4.5 | 51.8 | 11.6 | 0.8041 | 0.1804 | 0.0155 |
| EH | 5 | GBR3 | 4.6 | 54.3 | 11.8 | 0.8092 | 0.1759 | 0.0149 |
| EH | 5 | GBR4 | 4.6 | 53.7 | 11.8 | 0.8079 | 0.1770 | 0.0151 |
| EH | 5 | GBR5 | 4.5 | 53.2 | 11.7 | 0.8070 | 0.1779 | 0.0152 |
| EH | 5 | CEU1 | 2.2 | 23.8 | 10.7 | 0.6709 | 0.3010 | 0.0282 |
| EH | 5 | CEU2 | 4.4 | 49.9 | 11.5 | 0.8002 | 0.1838 | 0.0160 |
| EH | 5 | CEU3 | 4.8 | 44.6 | 9.2 | 0.8138 | 0.1679 | 0.0183 |
| EH | 10 | GBR1 | 6.6 | 59.5 | 9.0 | 0.8557 | 0.1299 | 0.0144 |
| EH | 10 | GBR2 | 6.6 | 60.2 | 9.1 | 0.8567 | 0.1291 | 0.0142 |
| EH | 10 | GBR3 | 6.6 | 59.6 | 9.0 | 0.8559 | 0.1298 | 0.0144 |
| EH | 10 | GBR4 | 6.6 | 59.4 | 9.0 | 0.8556 | 0.1300 | 0.0144 |
| EH | 10 | GBR5 | 6.5 | 58.9 | 9.0 | 0.8548 | 0.1307 | 0.0145 |
| EH | 10 | CEU1 | 4.0 | 27.8 | 7.0 | 0.7762 | 0.1959 | 0.0279 |
| EH | 10 | CEU2 | 6.7 | 60.6 | 9.1 | 0.8573 | 0.1285 | 0.0141 |
| EH | 10 | CEU3 | 6.1 | 53.1 | 8.7 | 0.8454 | 0.1386 | 0.0159 |
| MH | 5 | GBR1 | 334.9 | 7.27E+05 | 2170.8 | 0.9970 | 0.0030 | 1.37E-06 |
| MH | 5 | GBR2 | 246.3 | 5.91E+05 | 2401.4 | 0.9960 | 0.0040 | 1.68E-06 |
| MH | 5 | GBR3 | 356.5 | 7.98E+05 | 2239.2 | 0.9972 | 0.0028 | 1.25E-06 |
| MH | 5 | GBR4 | 341.4 | 5.80E+05 | 1700.0 | 0.9971 | 0.0029 | 1.72E-06 |
| MH | 5 | GBR5 | 337.1 | 9.46E+05 | 2807.1 | 0.9970 | 0.0030 | 1.05E-06 |
| MH | 5 | CEU1 | 221.7 | 6.51E+05 | 2937.4 | 0.9955 | 0.0045 | 1.53E-06 |
| MH | 5 | CEU2 | 293.3 | 7.68E+05 | 2619.3 | 0.9966 | 0.0034 | 1.30E-06 |
| MH | 5 | CEU3 | 242.6 | 9.61E+05 | 3960.0 | 0.9959 | 0.0041 | 1.04E-06 |
| MH | 10 | GBR1 | 375.8 | 7.88E+05 | 2097.7 | 0.9973 | 0.0027 | 1.27E-06 |
| MH | 10 | GBR2 | 279.9 | 6.67E+05 | 2383.5 | 0.9964 | 0.0036 | 1.49E-06 |
| MH | 10 | GBR3 | 412.0 | 9.03E+05 | 2191.0 | 0.9976 | 0.0024 | 1.11E-06 |
| MH | 10 | GBR4 | 342.8 | 6.89E+05 | 2008.5 | 0.9971 | 0.0029 | 1.45E-06 |
| MH | 10 | GBR5 | 441.3 | 9.99E+05 | 2263.5 | 0.9977 | 0.0023 | 9.99E-07 |
| MH | 10 | CEU1 | 227.0 | 6.40E+05 | 2819.8 | 0.9956 | 0.0044 | 1.56E-06 |
| MH | 10 | CEU2 | 296.7 | 7.27E+05 | 2450.2 | 0.9966 | 0.0034 | 1.37E-06 |
| MH | 10 | CEU3 | 325.6 | 6.38E+05 | 1960.1 | 0.9969 | 0.0031 | 1.56E-06 |

**Table S7** - Likelihood ratios and posterior probabilities computed using the generalized Poisson distribution (to account for underdispersion in the segment counts) based on the observed merged TJ matches in the EH and MH descendants under the 3 paternity scenarios for each scan.

| | | | Likelihood Ratio | | | Posterior Probability | | |
|---|---|---|---|---|---|---|---|---|
| **branch** | **threshold** | **scan** | **TJ:RJ** | **TJ:RJO** | **RJ:RJO** | **TJ** | **RJ** | **RJO** |
| EH | 5 | GBR1 | 5.1 | 52.3 | 10.2 | 0.8234 | 0.1609 | 0.0157 |
| EH | 5 | GBR2 | 5.1 | 52.7 | 10.2 | 0.8240 | 0.1603 | 0.0156 |
| EH | 5 | GBR3 | 5.3 | 55.2 | 10.4 | 0.8287 | 0.1563 | 0.0150 |
| EH | 5 | GBR4 | 5.3 | 54.6 | 10.4 | 0.8276 | 0.1573 | 0.0152 |
| EH | 5 | GBR5 | 5.2 | 54.1 | 10.3 | 0.8267 | 0.1580 | 0.0153 |
| EH | 5 | CEU1 | 2.5 | 21.9 | 8.8 | 0.6903 | 0.2781 | 0.0315 |
| EH | 5 | CEU2 | 5.0 | 50.8 | 10.1 | 0.8204 | 0.1634 | 0.0161 |
| EH | 5 | CEU3 | 5.6 | 47.7 | 8.6 | 0.8326 | 0.1499 | 0.0175 |
| EH | 10 | GBR1 | 7.2 | 62.3 | 8.7 | 0.8653 | 0.1208 | 0.0139 |
| EH | 10 | GBR2 | 7.2 | 63.0 | 8.7 | 0.8662 | 0.1201 | 0.0138 |
| EH | 10 | GBR3 | 7.2 | 62.4 | 8.7 | 0.8654 | 0.1207 | 0.0139 |
| EH | 10 | GBR4 | 7.2 | 62.2 | 8.7 | 0.8651 | 0.1210 | 0.0139 |
| EH | 10 | GBR5 | 7.1 | 61.6 | 8.7 | 0.8644 | 0.1215 | 0.0140 |
| EH | 10 | CEU1 | 4.3 | 29.1 | 6.8 | 0.7896 | 0.1833 | 0.0271 |
| EH | 10 | CEU2 | 7.3 | 63.4 | 8.7 | 0.8668 | 0.1196 | 0.0137 |
| EH | 10 | CEU3 | 6.6 | 55.5 | 8.4 | 0.8555 | 0.1290 | 0.0154 |
| MH | 5 | GBR1 | 433.9 | 9.43E+05 | 2174.4 | 0.9977 | 0.0023 | 1.06E-06 |
| MH | 5 | GBR2 | 345.6 | 8.03E+05 | 2324.1 | 0.9971 | 0.0029 | 1.24E-06 |
| MH | 5 | GBR3 | 461.8 | 1.04E+06 | 2242.9 | 0.9978 | 0.0022 | 9.63E-07 |
| MH | 5 | GBR4 | 402.4 | 7.05E+05 | 1751.4 | 0.9975 | 0.0025 | 1.42E-06 |
| MH | 5 | GBR5 | 473.0 | 1.29E+06 | 2716.7 | 0.9979 | 0.0021 | 7.76E-07 |
| MH | 5 | CEU1 | 332.2 | 9.07E+05 | 2729.5 | 0.9970 | 0.0030 | 1.10E-06 |
| MH | 5 | CEU2 | 411.6 | 1.04E+06 | 2535.0 | 0.9976 | 0.0024 | 9.56E-07 |
| MH | 5 | CEU3 | 382.6 | 1.34E+06 | 3511.0 | 0.9974 | 0.0026 | 7.42E-07 |
| MH | 10 | GBR1 | 505.9 | 1.16E+06 | 2296.7 | 0.9980 | 0.0020 | 8.59E-07 |
| MH | 10 | GBR2 | 425.4 | 1.08E+06 | 2540.8 | 0.9977 | 0.0023 | 9.23E-07 |
| MH | 10 | GBR3 | 554.6 | 1.33E+06 | 2398.8 | 0.9982 | 0.0018 | 7.50E-07 |
| MH | 10 | GBR4 | 461.5 | 1.01E+06 | 2199.0 | 0.9978 | 0.0022 | 9.83E-07 |
| MH | 10 | GBR5 | 594.1 | 1.47E+06 | 2478.2 | 0.9983 | 0.0017 | 6.78E-07 |
| MH | 10 | CEU1 | 383.9 | 1.12E+06 | 2906.3 | 0.9974 | 0.0026 | 8.94E-07 |
| MH | 10 | CEU2 | 450.9 | 1.18E+06 | 2611.9 | 0.9978 | 0.0022 | 8.47E-07 |
| MH | 10 | CEU3 | 438.3 | 9.41E+05 | 2146.0 | 0.9977 | 0.0023 | 1.06E-06 |

#### Resampled scans

We used the resampling strategy to generate new combinations of scan results based on the GBR reference panel to assess the variation in statistics due to differences between IBD scans. Supplemental Data Table S2: Resampled Statistical Results contains summary statistics for the likelihood ratios and posterior probabilities computed from the resampled scans. Figure S33 shows histograms of the computed statistics.

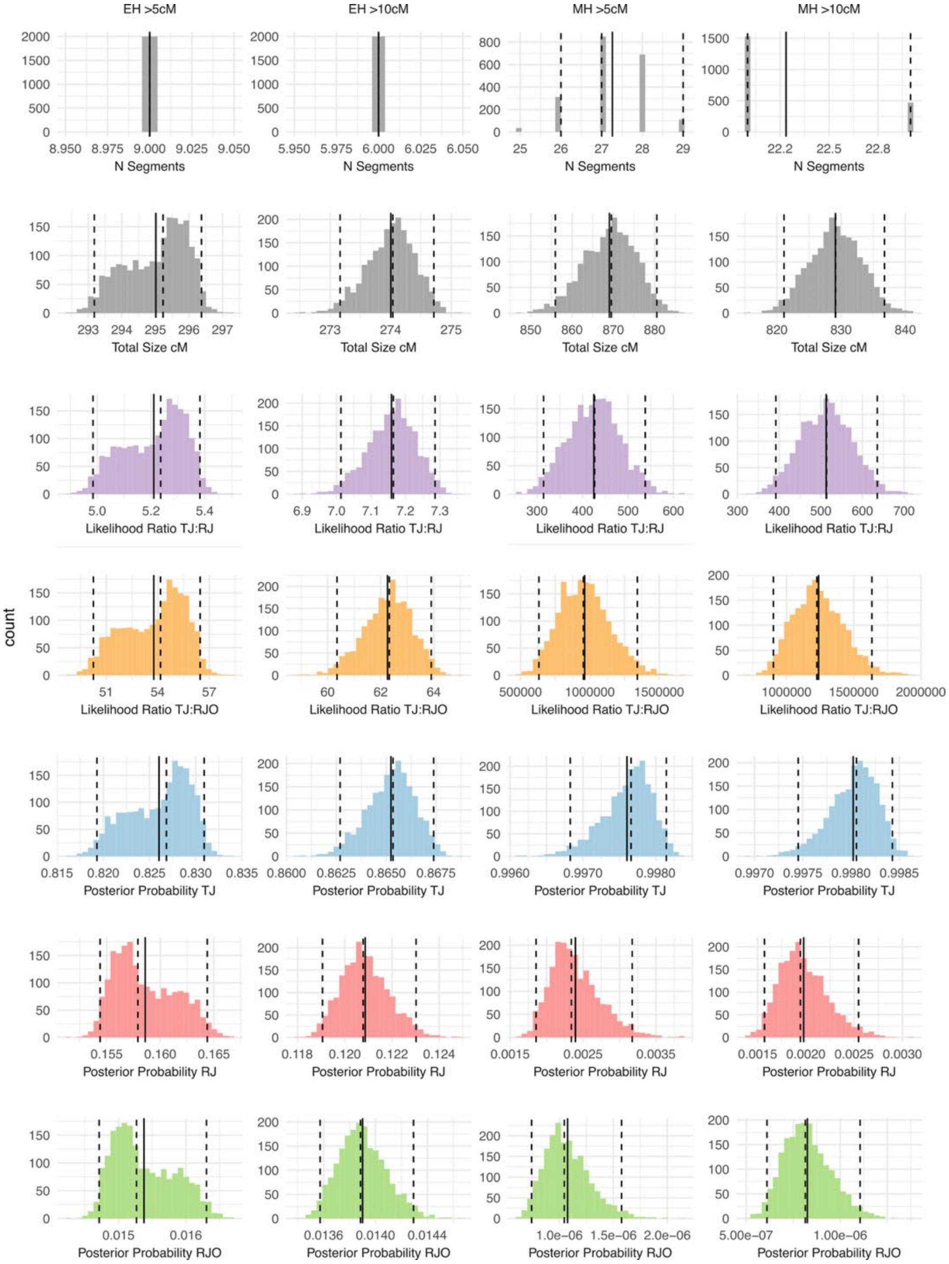

**Figure S33** - Histograms of statistics obtained from the resampled scans based on the GBR reference panel (N=2,000) for each Hemings branch (EH: Eston Hemings, MH: Madison Hemings) and minimum IBD segment size threshold (5 cm and 10 cm) (columns). Rows: number of IBD segments, total size of IBD in centimorgans, TJ:RJ likelihood ratio, TJ:RJO likelihood ratio, posterior probability of the TJ paternity scenario, posterior probability of the RJ scenario, posterior probability of the RJO scenario. The solid vertical line marks the mean value and the dashed lines indicate the 2.5th, 50th, and 97.5th percentiles.

In addition to the posterior probabilities calculated by assuming equal prior probabilities for the 3 paternity scenarios, we used the median likelihoods from resampling to compute the posterior probability of the TJ scenario across the range of possible priors (Figure S34). We found that when considering segments both greater than 5 cM and 10 cM as evidence, the TJ scenario is favored across a wide range of priors. For MH, we found that the posterior probability of the TJ scenario exceeded 95% for all but the strongest priors against the TJ scenario. We see a weaker pattern for EH, but note that the TJ posterior probability is greater than 50% across most of the space.

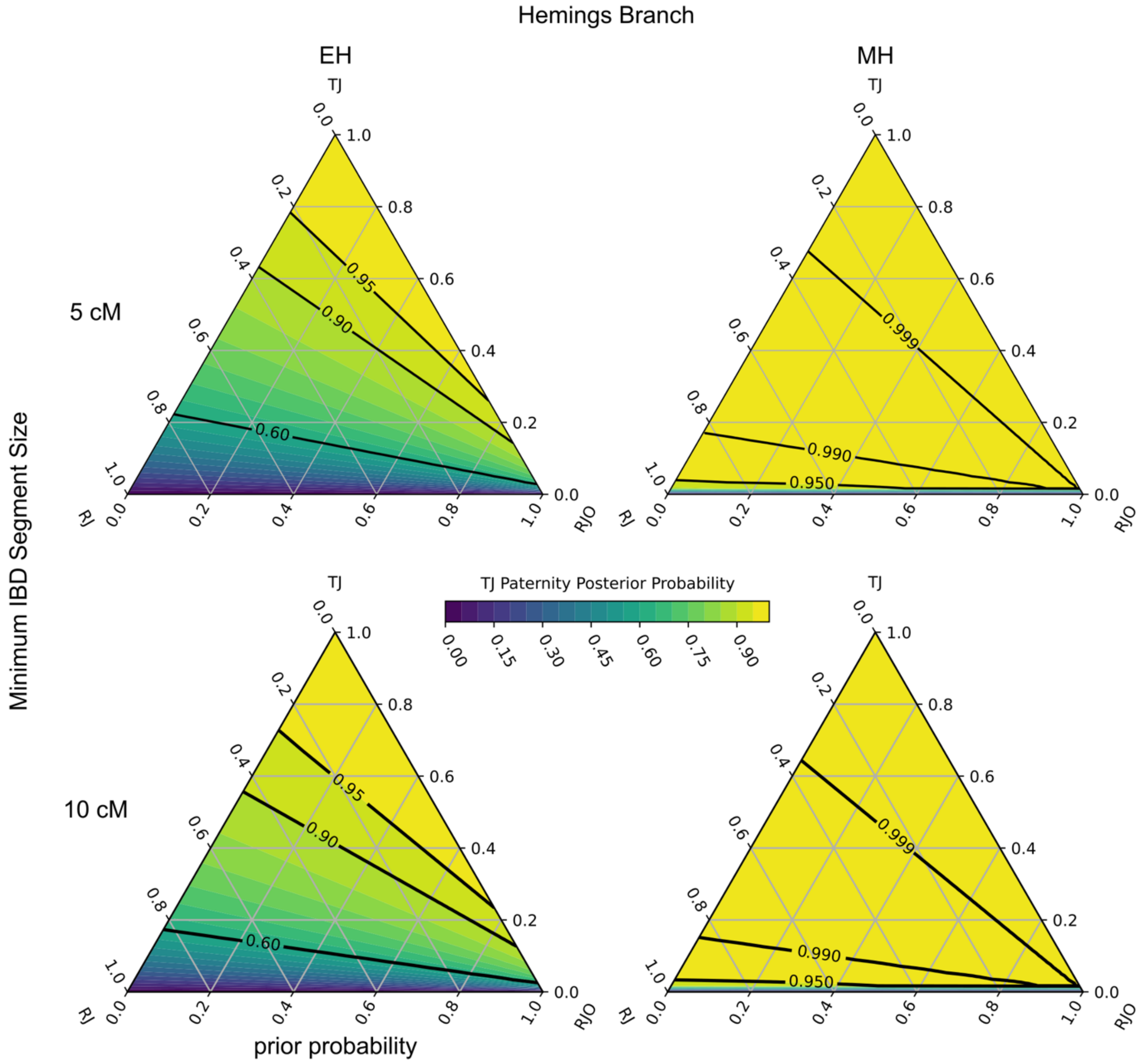

**Figure S34** - Posterior probability of the TJ paternity scenario across the range of possible prior probabilities when considering only the TJ, RJ and RJO scenarios based on the median observed likelihoods given each scenario from the resampled scans. The top row shows ternary plots when considering segments greater than 5 cM in length and the bottom row contains plots when only segments greater than 10 cM are considered.

#### Comparison with Simulations

We compared the likelihood ratios obtained from the observed TJ IBD segments with ratios computed using IBD segments from genealogical segments. We calculated the TJ:RJ and TJ:RJO likelihood ratios for each simulation replicate in the withheld test set of 4,000 replicates per scenario using the same likelihood parameters described previously. Figure S35 shows histograms describing the density of computed likelihood ratios for each simulated scenario. We noted that the distributions overlap and some fraction of simulations produced likelihood ratios that were not in agreement with the true simulated scenario. We also observed more overlap in the distributions in the EH branch compared with the MH branch, as expected, as the MH branch has broader sampling and more individuals to distinguish between scenarios. The observed likelihood ratios from the resampled scans are close to the medians of the TJ scenario simulation distributions. When compared to the simulations of the RJ and RJO scenarios, the observed likelihood ratios would represent extreme values for those distributions.

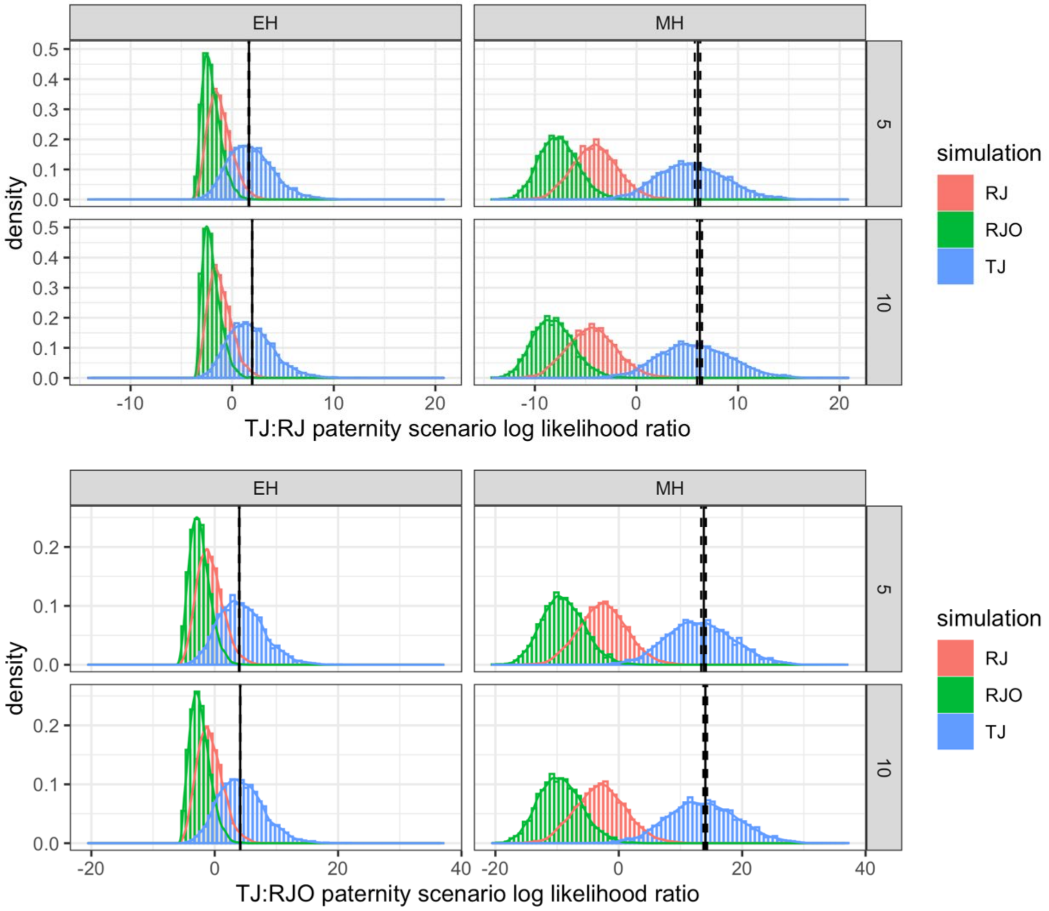

**Figure S35** - Comparison of likelihood ratios obtained from the observed TJ IBD segments and IBD segments from genealogy simulations. Histograms and smoothed-density estimates show the densities of log-likelihood ratios comparing the TJ and RJ scenarios (top panel) and TJ and RJO scenarios (bottom panel) computed from genealogical simulations of each scenario (4,000 replicates per scenario) in the

Eston Hemings branch (EH; left column) and Madison Hemings branch (MH; right column) using the 5 cM and 10 cM minimum segment size thresholds (rows). The observed mean log-likelihood ratios from resampling are shown as solid vertical lines and dashed vertical lines indicate the 2.5th and 97.5th percentiles (N=2,000 resampled scans).

##### Joint Probability

Finally, we estimated the joint probability that Thomas Jefferson was the father of both Eston and Madison Hemings, the probability that he was the father of at least one of Eston and Madison, and the probability he was not the father of either one. We used the branch-specific likelihoods from the resampled scans and calculated joint posterior probabilities assuming equal prior probabilities and independence between the paternity scenarios for EH and MH. This assumption is not necessarily likely. It may be argued that it should be considered more likely for EH and MH to share the same father. However, it is difficult to determine more realistic prior probabilities so we continue to assume equal prior probabilities for each scenario and combination of scenarios. To obtain the joint likelihoods of the IBD segments observed for EH and MH, we take the product of the individual likelihoods for each paternity scenario combination and normalize them by the total probability. To simplify the calculation, we assume the models for number and size of IBD segments for each individual and paternity scenario are independent. While TJ IBD in RJO is partially dependent on TJ IBD in RJ, we reason that the subsequent generations of recombination have a greater impact on the amount of IBD detected in the Hemings descendants than differences in the initial IBD sharing conditions. We estimated the joint probabilities assuming that Eston and Madison Hemings shared the same father using a similar approach (see Main Text and Supplemental Data Table S2: Resampled Statistical Results).

Tables S8 and S9 show the joint posterior probabilities obtained by normalizing the joint likelihoods for each Hemings and paternity scenario. Based on the ERSA likelihood, equal prior probabilities for all paternity scenarios, and independence assumptions, we estimate the probability that TJ is the father of both MH and EH to be 80% using the 5 cM threshold and 85% using the 10 cM. With both thresholds, we estimate the probability that TJ fathered at least one of MH or EH to be 99.9% and the probability that he did not father either one to be less than 0.1% or approximately 1 in 1,000 (see Supplemental Data Table S2: Resampled Statistical Results).

**Table S8** - Mean joint posterior probabilities for EH and MH paternity scenarios estimated using the ERSA likelihood model and the 5 cM segment threshold from the resampled scans, assuming equal prior probabilities and independence for the paternity scenarios between EH and MH. Central 95th percentile intervals are included in parentheses. Cells are shaded blue to indicate a scenario where TJ is the father of at least one of EH and MH and gray to indicate scenarios where TJ is not the father of either one.

| Threshold: 5cm | | MH | | |
|---|---|---|---|---|
| | | TJ | RJ | RJO |
| EH | TJ | 0.8240 (0.8173–0.8290) | 2.0e-03 (1.5e-03–2.6e-03) | 8.9e-07 (6.1e-07–1.3e-06) |
| | RJ | 0.1582 (0.1540–0.1639) | 3.8e-04 (2.9e-04–5.1e-04) | 1.7e-07 (1.2e-07–2.5e-07) |
| | RJO | 0.0153 (0.0147–0.0163) | 3.7e-05 (2.8e-05-5.0e-05) | 1.7e-08 (1.1e-08-2.4e-08) |

| | |
|---|---|
| Probability TJ father of EH AND MH: | 0.8240 (0.8173–0.829) |
| Probability TJ father of EH OR MH: | 0.9996 (0.9994–0.9997) |
| Probability TJ not father of EH AND MH: | 0.0004 (0.0003–0.0006) |

**Table S9** - Mean joint posterior probabilities for EH and MH paternity scenarios estimated using the ERSA likelihood model and the 10 cM segment threshold from the resampled scans, assuming equal prior probabilities and independence for the paternity scenarios between EH and MH. Central 95th percentile intervals are included in parentheses. Cells are shaded blue to indicate a scenario where TJ is the father of at least one of EH and MH and gray to indicate scenarios where TJ is not the father of either one.

| Threshold: 10cm | | MH | | |
|---|---|---|---|---|
| | | TJ | RJ | RJO |
| EH | TJ | 0.8635 (0.8609–0.8657) | 1.7e-03 (1.4e-03–2.2e-03) | 7.1e-07 (5.3e-07–9.6e-07) |
| | RJ | 0.1206 (0.1188–0.1228) | 2.4e-04 (1.9e-04–3.1e-04) | 1.0e-07 (7.4e-08–1.3e-07) |
| | RJO | 0.0139 (0.0135–0.0143) | 2.7e-05 (2.2e-05–3.5e-05) | 1.1e-08 (8.5e-09–1.5e-08) |

| | |
|---|---|
| Probability TJ father of EH AND MH: | 0.8635 (0.8609–0.8657) |
| Probability TJ father of EH OR MH: | 0.9997 (0.9997–0.9998) |
| Probability TJ not father of EH AND MH: | 0.0003 (0.0002–0.0003) |

#### Alternative Likelihood Model

As an alternative to the ERSA likelihood model, we considered a likelihood function based solely on the total merged IBD detected in the descendant individuals. The performance of this function would be less affected by inaccuracies in the number of segments detected (e.g., segments that are incorrectly reported as multiple segments). However, a total IBD model may not accurately estimate likelihoods for extreme values where there is little support from simulations.

We used a gamma distribution to approximate the likelihood of the total IBD observation adjusted for the minimum match threshold given the scenario model.

$$T = \sum_{i \in s} s_i - t$$

A gamma distribution is reasonable choice as a special case of the gamma distribution, the Erlang distribution, is the distribution of the sum of $k$ independent exponential distributions with mean

$1/\lambda$. However, the gamma distribution has two parameters versus the single-parameter Poisson or exponential distributions and is harder to fit to the simulated data. We used the `fitdistrplus` (DOI: 10.18637/jss.v064.i04) package in R to fit the parameters of the gamma distribution by maximum likelihood estimation to the simulated data for each genealogy, scenario, and threshold combination. The estimated shape and scale parameters did not closely match the mean number of segments or the mean adjusted segment sizes for the simulations (Table S10) but they varied in similar patterns across each branch, threshold and scenario. The histograms and quantile-quantile plots comparing the empirical and theoretical distributions suggested a good fit in most cases (see Figure S36).

**Table S10** - Parameters of the gamma distributions used to model total Thomas Jefferson IBD sharing under the 3 paternity scenarios for each branch and minimum segment size threshold combination. The mean number of segments and mean segment size adjusted by the threshold size from the simulations used to fit the parameters are included for comparison.

| | | | Simulations | | Gamma parameters | | |
|---|---|---|---|---|---|---|---|
| **Branch** | **Threshold (cM)** | **Scenario** | **Mean N segments** | **Mean segment size - threshold** | **shape** | **scale** | **rate (1/scale)** |
| EH | 5 | TJ | 8.90 | 29.58 | 5.02205 | 52.46176 | 0.01906 |
| EH | 5 | RJ | 6.77 | 17.28 | 2.95726 | 39.65191 | 0.02522 |
| EH | 5 | RJO | 4.84 | 14.21 | 1.96789 | 35.33457 | 0.02830 |
| EH | 10 | TJ | 7.40 | 30.09 | 4.13096 | 53.94597 | 0.01854 |
| EH | 10 | RJ | 5.07 | 17.27 | 2.25370 | 39.19355 | 0.02551 |
| EH | 10 | RJO | 3.39 | 14.26 | 1.51937 | 33.15338 | 0.03016 |
| MH | 5 | TJ | 28.63 | 25.38 | 25.16736 | 28.87059 | 0.03464 |
| MH | 5 | RJ | 20.50 | 15.63 | 12.34335 | 25.96098 | 0.03852 |
| MH | 5 | RJO | 14.39 | 13.13 | 7.42683 | 25.43147 | 0.03932 |
| MH | 10 | TJ | 24.06 | 24.66 | 19.76034 | 30.02726 | 0.03330 |
| MH | 10 | RJ | 15.18 | 15.21 | 8.71671 | 26.48509 | 0.03776 |
| MH | 10 | RJO | 9.94 | 12.87 | 4.91865 | 26.02486 | 0.03842 |

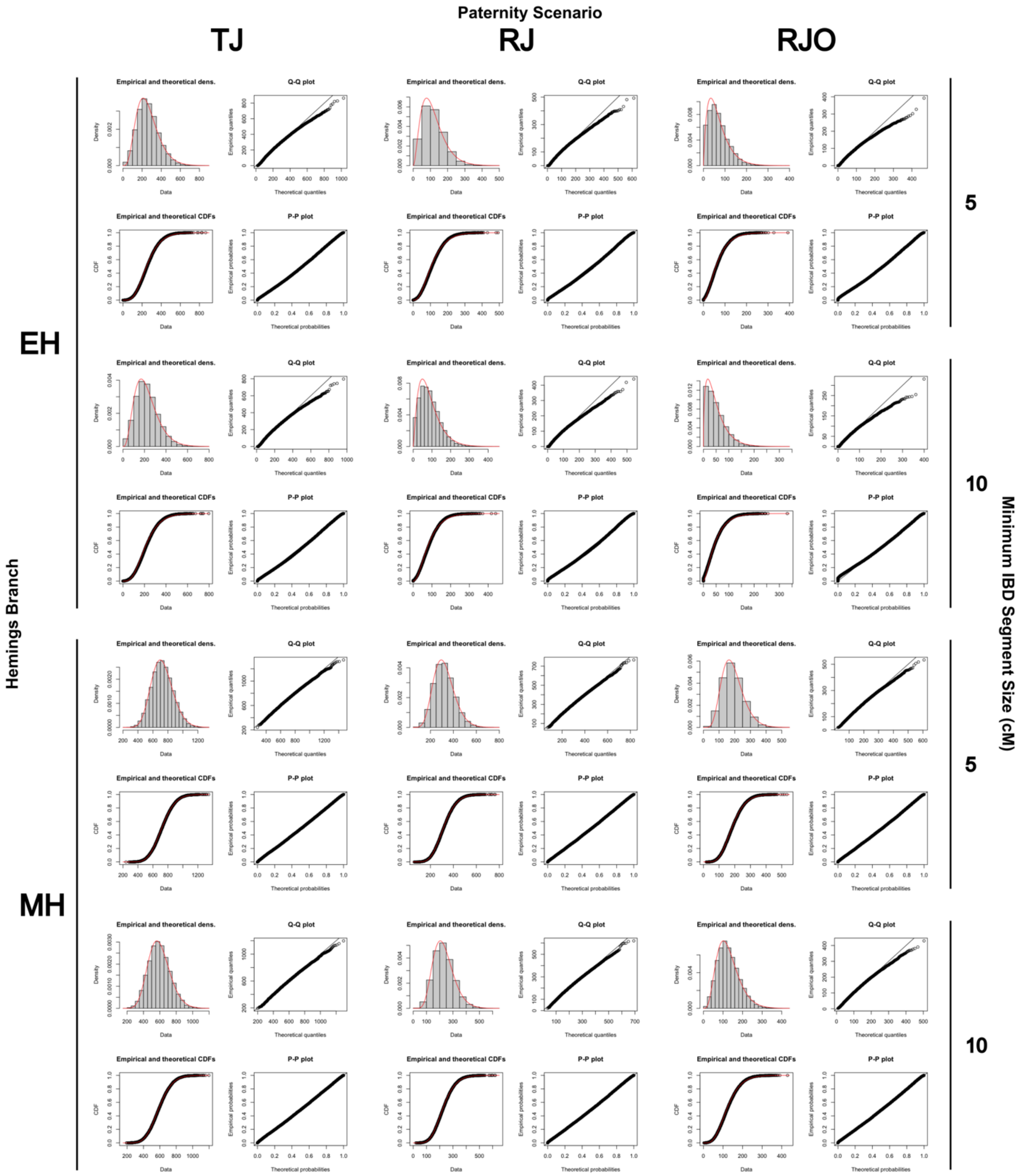

**Figure S36** - Comparisons of the distribution of total IBD with the gamma approximation for each combination of paternity scenario (TJ, RJ, or RJO), Hemings branch (EH or MH), and minimum IBD segment size (5 cM or 10 cM). There are four sub-plots for each model: a histogram comparing the empirical and theoretical distributions (top left), a quantile-quantile plot (top right), a comparison the empirical and theoretical cumulative distribution functions (CDF) (bottom right), and a probability-probability plot (bottom right).

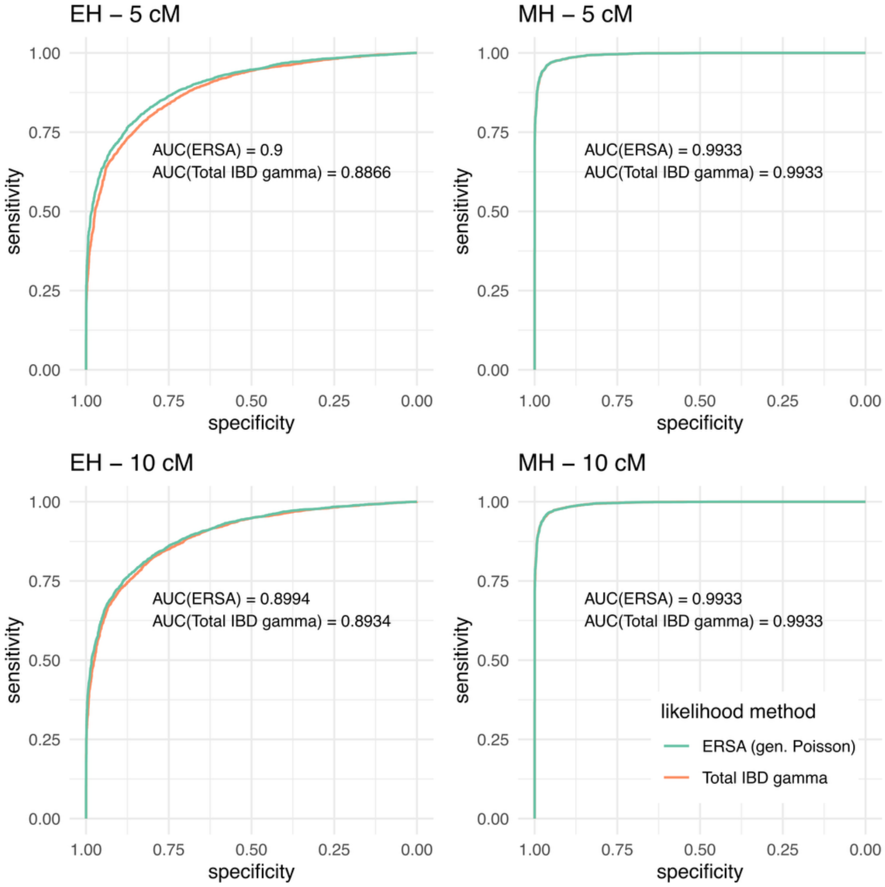

**Figure S37** - Receiver operating characteristic plots comparing the performance of the ERSA likelihood function (generalized Poisson) and the Total IBD gamma likelihood function in distinguishing between the TJ and RJ scenarios in simulations (N=4,000 per scenario).

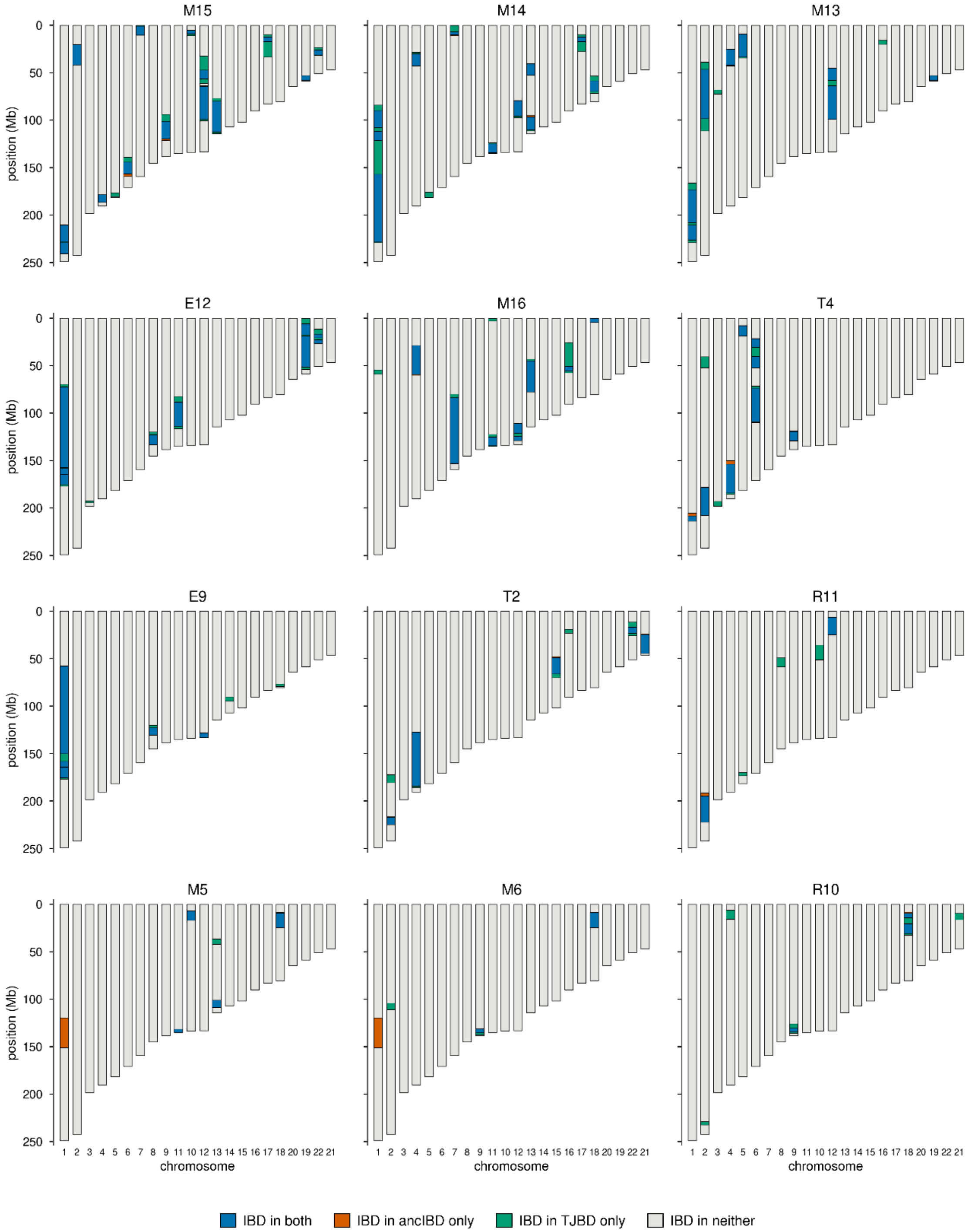

**Figure S38** – Comparison of TJBD and ancIBD scans for segments of IBD. We imputed Thomas Jefferson's genome and the genomes of the descendant panel with Glimpse2 v2.0.1 using a high-coverage 1KG dataset as a reference panel. We then called IBD segments using ancIBD with default settings and a minimum length cut off for detecting IBD segments of 8cM. Because the 1240k SNP positions and allele frequency files used by ancIBD are distributed on hg19 coordinates and our imputation panel was on GRCh38 we rebuilt these resources on GRCh38. 1240k physical snp positions were transferred to GRCh38 using UCSC liftover (hg19ToHg38.over.chain.gz) removing sites which mapped to a different chromosome after liftover resulting in retention of 99.85% of 1240k sites. To regenerate allele frequency files on GRCh38 we recomputed allele frequencies directly from our 1KG reference panel. Karyograms are color-coded by IBD segments detected by one, both, or neither method.

**Table S11** - Likelihood ratios and posterior probabilities based on the observed total TJ IBD in the EH and MH descendants under the 3 paternity scenarios for each scan.

| | | | Likelihood Ratios | | | Posterior Probability | | |
|---|---|---|---|---|---|---|---|---|
| branch | threshold | scan | TJ:RJ | TJ:RJO | RJ:RJO | TJ | RJ | RJO |
| EH | 5 | GBR1 | 3.9 | 21.3 | 5.4 | 0.7684 | 0.1956 | 0.0360 |
| EH | 5 | GBR2 | 3.9 | 21.4 | 5.4 | 0.7688 | 0.1953 | 0.0359 |
| EH | 5 | GBR3 | 4.0 | 22.0 | 5.5 | 0.7724 | 0.1925 | 0.0351 |
| EH | 5 | GBR4 | 4.0 | 21.9 | 5.5 | 0.7715 | 0.1932 | 0.0353 |
| EH | 5 | GBR5 | 4.0 | 21.8 | 5.5 | 0.7709 | 0.1937 | 0.0354 |
| EH | 5 | CEU1 | 3.0 | 14.5 | 4.8 | 0.7145 | 0.2361 | 0.0493 |
| EH | 5 | CEU2 | 3.9 | 21.0 | 5.4 | 0.7661 | 0.1974 | 0.0365 |
| EH | 5 | CEU3 | 3.5 | 18.2 | 5.2 | 0.7473 | 0.2116 | 0.0410 |
| EH | 10 | GBR1 | 4.6 | 26.3 | 5.7 | 0.7969 | 0.1728 | 0.0303 |
| EH | 10 | GBR2 | 4.6 | 26.5 | 5.7 | 0.7977 | 0.1722 | 0.0301 |
| EH | 10 | GBR3 | 4.6 | 26.3 | 5.7 | 0.7970 | 0.1727 | 0.0303 |
| EH | 10 | GBR4 | 4.6 | 26.2 | 5.7 | 0.7967 | 0.1729 | 0.0304 |
| EH | 10 | GBR5 | 4.6 | 26.1 | 5.7 | 0.7961 | 0.1734 | 0.0305 |
| EH | 10 | CEU1 | 3.3 | 15.9 | 4.8 | 0.7322 | 0.2217 | 0.0461 |
| EH | 10 | CEU2 | 4.6 | 26.6 | 5.7 | 0.7982 | 0.1717 | 0.0300 |
| EH | 10 | CEU3 | 4.4 | 24.4 | 5.6 | 0.7881 | 0.1796 | 0.0323 |
| MH | 5 | GBR1 | 419.1 | 153843.3 | 367.0 | 0.9976 | 0.0024 | 6.48E-06 |
| MH | 5 | GBR2 | 365.4 | 127817.5 | 349.8 | 0.9973 | 0.0027 | 7.80E-06 |
| MH | 5 | GBR3 | 442.5 | 165527.3 | 374.1 | 0.9977 | 0.0023 | 6.03E-06 |
| MH | 5 | GBR4 | 373.7 | 131746.5 | 352.5 | 0.9973 | 0.0027 | 7.57E-06 |
| MH | 5 | GBR5 | 480.2 | 184863.8 | 385.0 | 0.9979 | 0.0021 | 5.40E-06 |
| MH | 5 | CEU1 | 380.1 | 134813.5 | 354.7 | 0.9974 | 0.0026 | 7.40E-06 |
| MH | 5 | CEU2 | 425.6 | 157054.6 | 369.0 | 0.9976 | 0.0023 | 6.35E-06 |
| MH | 5 | CEU3 | 468.6 | 178873.8 | 381.7 | 0.9979 | 0.0021 | 5.58E-06 |
| MH | 10 | GBR1 | 490.0 | 95338.6 | 194.6 | 0.9980 | 0.0020 | 1.05E-05 |
| MH | 10 | GBR2 | 383.4 | 69185.7 | 180.5 | 0.9974 | 0.0026 | 1.44E-05 |
| MH | 10 | GBR3 | 532.1 | 106162.7 | 199.5 | 0.9981 | 0.0019 | 9.40E-06 |
| MH | 10 | GBR4 | 451.2 | 85585.5 | 189.7 | 0.9978 | 0.0022 | 1.17E-05 |
| MH | 10 | GBR5 | 565.8 | 115020.1 | 203.3 | 0.9982 | 0.0018 | 8.68E-06 |
| MH | 10 | CEU1 | 323.3 | 55371.4 | 171.3 | 0.9969 | 0.0031 | 1.80E-05 |
| MH | 10 | CEU2 | 404.3 | 74149.8 | 183.4 | 0.9975 | 0.0025 | 1.35E-05 |
| MH | 10 | CEU3 | 430.6 | 80531.8 | 187.0 | 0.9977 | 0.0023 | 1.24E-05 |

**Human Subjects information for DNA collection**

DNA sampling of known and possible descendants of Thomas Jefferson was carried out under Human Subjects Research protocols HS14052, HS14052-02, and HS24024 as reviewed and approved by the Human Subjects Institutional Review Board of the Smithsonian Institution.

The research originated with the inadvertent discovery of three hairs in the Jefferson Bible during the course of its conservation at the Smithsonian from 2009-2011 and the possibility that those hairs might yield DNA that could resolve the question of Thomas Jefferson's paternity of Sally Hemings's children. Additionally, the Smithsonian concluded agreements to analyze packets of purported Thomas Jefferson hair in the collections of the Thomas Jefferson Foundation at Monticello.

Anticipating that we would eventually be able to recover and sequence DNA from those hairs, we developed a research design that called for sampling the generationally closest living lineal descendants of Thomas Jefferson, his brother Randolph, and Sally Hemings as they would have the largest amounts of Thomas Jefferson ancestral DNA, if present. Genealogical and historical research found no living 4th generation descendants but identified several living 5th generation descendants of Thomas Jefferson and of Sally Hemings and several 6th generation descendants of Randolph Jefferson. We concentrated on descendants of Thomas Jefferson's daughter Maria given the existence of more intrafamilial ancestral marriages in the genealogy of her sister Martha. No descendants of two of Sally Hemings's children, Beverly and Harriet, have ever been identified in the historical record, so we were restricted to descendants of her sons Madison and Eston. While the research was focused on sampling participants genealogically widely dispersed in their descent from the key ancestors because that would provide useful data on ancestry through IBD distribution, we also sampled several sets of siblings due to their historical importance, and in some cases out of deference to participant wishes.

We carried out work involving descendants following procedures approved by the Smithsonian's Institutional Review Board (IRB). Descendants were contacted, first by telephone or letter by members of the research team and by follow-up conversations and in-person meetings to explain the inadvertent discovery of the hairs in the Jefferson Bible and in the collections at Monticello and the possibility that they might yield DNA that could resolve the paternity question. Initial conversations also enabled research team members to confirm genealogical relationships and histories and assess the descendants' willingness to participate.

In almost every case when members of the research team visited participants, either in their home or that of a close relative, other family members such as children, spouses, siblings, other kin and friends were present. Following introductions, research team members engaged descendants and family members in open-ended discussion, starting with the discovery of possible Thomas Jefferson hairs in the Jefferson Bible and in annotated packets at Monticello. Such discussions typically recalled the paternity controversy from historical, scientific and familial perspectives.

Participants were assured that the research would seek the truth about the paternity question, whatever the result, consistent with Thomas Jefferson's own admonition "to follow truth wherever it may lead." Informed consent forms approved by Smithsonian legal counsel and IRB were provided ahead of in-person meetings so that participants could familiarize themselves with their rights and conditions of participation and have family members review such forms if they desired, and which several did. The forms specified the principal investigators at the Smithsonian and at the University of California Santa Cruz and provided contact information for the Smithsonian's research compliance staff member who could be independently contacted by participants if desired.

All descendants approached for DNA samples agreed to participate, gave their informed consent, and provided saliva samples—12 into a tube, and 2 by mouth swab. They were also asked to indicate whether or not they wanted to know the results, and all participants did.

As the project proceeded, participants were asked, either at the time of sampling or subsequently whether or not they wanted to remain anonymous when the results of the research were to be publicly reported. All participants, and in a few cases, the heirs of deceased participants agreed to the public disclosure of their names.

As participants were sampled over several years, we reported back to them on the status of DNA retrieval from the Bible and Monticello hairs. Participants were informed that the recovery and sequencing of the historical DNA was complicated, would take time, involve innovative methods, and require scientific review. Most of the sampled descendants were elderly, many in their 80s and 90s, and over the course of the project, 9 of the 14 passed away. In those cases, members of the research team informed spouses, children or other family members of research progress and results.

When sufficient progress was made in sequencing the Thomas Jefferson genome and generating IBD measures and chromosomal IBD charts for individual participants, they were provided to descendants and/or family members orally and/or visually and explained sometimes with a full review of the project using on-screen or printed-out PowerPoint slides. Descendants and family members had ample opportunity to ask questions, make comments, and engage a research team member in discussion.

The research team also conducted video interviews with several descendants and with descendants' children and in one case, grandchild, obtaining informed consent to do so. Those who agreed to such video interviews also signed releases for the public posting of those videos by the Smithsonian.

Information from those numerous discussions over the course of more than a decade in some cases, and video interviews provided context to the historical issue of paternity and how it had impacted descendants and their families over generations. They provided particular familial and biographical stories and episodes related to the assertion and denial of racial identity and kinship with Thomas Jefferson, Sally Hemings and others. Those discussions are reported on in considerable detail in a book, *History by a Hair: Thomas Jefferson, Sally Hemings & Solving a 200-Year-Old Mystery* by Richard Kurin scheduled for publication on September 22, 2026 by Crown/Penguin Random House.

Descendants had various perspectives on the paternity issue and in many cases had experienced the contentious affirmation and/or denial of their ancestry and that of others. Participants and their family members overwhelmingly embraced the idea that the Smithsonian would objectively determine, if it could, the paternity issue, and that such should not be clouded by any biased, predetermined, or desired result. To maintain that objectivity in the face of more than 200 years of contention and controversy, the principal investigators did not include descendants as co-authors as such could be viewed by some as biasing the study findings in one way or another—and indeed descendants and family members who expressed an opinion on the matter all concurred. Descendant perspectives on the paternity issue and questions of identity and descent are amply discussed in the aforementioned book.

Prior to the publication of the book and this article, members of the research team briefed various descendant groups comprised of recognized Jefferson and Hemings family members, as well as related organizations. One of the descendants decided to opt out of the study and asked that their DNA be destroyed and data removed from the study – which it was.

The DNA sequence generated from Thomas Jefferson hair will be made available by written request to the study authors (see A Note on Thomas Jefferson DNA data availability below). The IBD charts which compare each descendant's genome to the Thomas Jefferson genome are included in the SOM and hence publicly available. However, the genome of each of the descendants is considered PII by the Smithsonian and participants did not consent to making that publicly available. Researchers seeking to verify specific results of the analysis are encouraged to contact the lead author at the University of California Santa Cruz to arrange appropriate data sharing for that purpose only. Since this article concentrates on the scientific methodology and analysis of the DNA study, descendants are identified by a code number. As the aforementioned book and a companion excerpt in *Smithsonian* magazine includes biographical stories, reflections, direct quotes, images and likenesses, descendants are named, as they agreed and as allowed under the conditions of IRB and a privacy review.

**A Note on the Use of Ancient DNA from Museum Collections**

When we initiated the project, there were no particular limitations to researching the hair of deceased historical figures. The seminal Belmont Report (1979) and the federal human subject research guidelines it generated dealt with ethical obligations to the living, not to the dead. The National Museum of American Indian Act (1989) and the Native American Graves Protection and Repatriation Act (1990) codified treatment of Native American remains, but until recent years, museums, including the Smithsonian, lacked policies regarding the acquisition, research, display and repatriation of non-Native human remains.

Increased attention to the acquisition of human remains—largely skeletal—from poor, minority and marginalized groups in the 19th and 20th centuries stimulated new policies and practices among several major museums, including the Smithsonian. Those policies have set conditions on new acquisitions, the research on and display of human remains, and prioritized repatriation and memorialization. Those policies are still evolving across the museum world. Institutions have considered questions of scope, for example, whether to include all human tissue as remains or exclude historical objects, like lockets, reliquaries and such that contain hair and bone. Central to policy formulation has been the issue of having informed consent from decedents or their descendants or heirs, appropriate organizations or representative communities for acquiring, displaying and researching remains.

In this case, the hairs in the Jefferson Bible were found in a Smithsonian artifact obtained more than a century ago from the Jefferson family and the research team spent years trying to determine whose they were. The locks of hair at Monticello that we examined had been provided voluntarily, one assumes, while Thomas Jefferson was alive, and by his family members on the day of his death. They were subsequently freely transferred by descendants to the Thomas Jefferson Foundation at Monticello, an institution dedicated to and representing his legacy. Genomic technology was not available during Jefferson's lifetime nor at the time of these transfers. The loan of strands of hair from those historic locks by the Thomas Jefferson Foundation at Monticello to the Smithsonian for destructive DNA retrieval and analysis by the University of California Santa Cruz was covered by formal agreements between the organizations.

The Smithsonian believed that there was a compelling justification for proceeding with the analysis of these hairs—providing scientific evidence related to a question that had occupied national attention for more than two centuries and was of critical importance to many descendants. This was believed to be consistent with Jefferson's own admonition to pursue truth wherever it might lead. And indeed descendants involved in the project proceeded with that purpose in mind. All were well informed about the discovery and analysis of what we assumed to be Thomas Jefferson's hair. They provided their DNA, understanding it would be compared to Jefferson's. And it was not only those descendants, but it was also dozens of their family members who participated in discussions, were familiar with the work, and supported its moving forward.

#### A Note on Thomas Jefferson DNA data availability

It is the intention of the authors to make available the DNA sequence data recovered from Thomas Jefferson's hairs to the research community and any interested party. These data, from a known, deceased individual of historical significance for whom there is an outstanding paternity question was unimagined by standard data archiving and sharing venues.

In support of the paper, we attempted to submit the complete, assembled mitochondrial genome to GenBank and the raw DNA sequence data from Thomas Jefferson hairs to NIH's Database of Genotypes and Phenotypes (dbGaP).

The response from GenBank following our mitochondrial genome submission is shown here:

```
Dear Collegue,

Thank you for your recent submission of mitochondrial DNA sequence data to GenBank. After review, we are unable to accept this submission for inclusion in GenBank due to the nature of this particular sample.

This is mitochondrial DNA from a known deceased individual who has living descendants. As such, privacy and consent considerations call for a controlled-access repository rather than an unrestricted access repository like GenBank. dbGaP (the database of Genotypes and Phenotypes) is designed for exactly this kind of situation, where access needs to be reviewed and granted on a case-by-case basis rather than open to anyone.

We'd suggest beginning the process of submitting this data to dbGaP as a non-NIH-funded study. NHGRI has agreed to be the sponsor for this data, which means the NHGRI Data Access Committee will review data access requests. NHGRI is a natural fit as sponsor because this data isn't tied to a specific disease or health condition, so it doesn't fall naturally under any of NIH's disease-specific Institutes. NHGRI supports research that generates broadly applicable genomic knowledge rather than research organized around a particular disease or organ system, which makes it well positioned to sponsor genomic data of this kind.

Please work with a Genomic Program Administrator (GPA) at NHGRI to register your study in dbGaP first.
More information can be found at https://grants.nih.gov/policy-and-compliance/policy-topics/sharing-policies/gds/register-submit-study-dbgap#registering-a-non-nih-funded-study-in-dbgap.
After you have registered your study, you will need to upload your genome through the dbGaP submission portal.
Anne Sturcke of the dbGaP team lead will be happy to help you with this process.
```

Please let us know if you have questions or if we can help connect you with the right contact.

Sincerely,

GBC-100 1
The GenBank Submissions Staff
Bethesda, Maryland USA

Subsequently, we registered the study with dbGaP. After submitting a description of the study and the data – in this case all the raw DNA sequence data from Thomas Jefferson hairs. After dbGaP review, the study was determined not to be a fit for dbGaP with the following written rationale from dbGaP staff:

Thank you for providing the additional questions. Our committee has reviewed your request and decided that your study would not be a fit for sponsorship by NHGRI to dbGaP under the NIH's Genomic Data Sharing (GDS) Policy (NOT-OD-14-124). The GDS Policy requires that submitted genomic data be de-identified and the privacy of study participants be maintained (see 5. Institutional Certification). The goal of this study was to identify the paternal ancestry of Martha Wayles's and Sally Hemings's descendants, which by nature does not main participant privacy and violates the requirement to not disclose the identities of research participants to NIH-designated repositories. While the individual (Thomas Jefferson) is deceased, the NIH still considers this human genomic data under the GDS Policy with the potential for identifiability. If the individual died before the GDS Policy's effective date (January 25, 2015), the NIH still requires that the submission is consistent, as appropriate, with applicable national, tribal, and state laws and regulations as well as relevant institutional policies.

Furthermore, NIH controlled-access data repositories (CADRs) support, store, and manage access to biomedical research data. Our committee felt this study did not qualify as biomedical research and wouldn't be an appropriate study for an NIH CADR. There were also additional concerns about the inability to protect the privacy of descendants and/or family members from incidental findings that could reveal health information about the Hemings or Jefferson families. A potential variant associated with a disease or disorder could lead to stigmatization, discrimination, and direct identification of family members.

Given these factors, NHGRI did not feel this study would be appropriate for our institute, though we believe the science is interesting and look forward to reading the publication. We hope that you're able to find the study an appropriate home. For additional information on maintaining participant privacy in genomic and biomedical research, please check out the following resources:

Privacy in Genomics
Principals and Best Practices for Protecting Participant Privacy
Supplemental Information to the NIH Policy for Data Management and Sharing: Protecting Privacy When Sharing Human Research Participant Data

Given the issues unique to DNA sequence data from Thomas Jefferson, uncontemplated by data archival and sharing platforms, we are left with either submitting the data to a completely open platform or making the data available by request. Available by request is currently deemed to be the least imperfect, available option.
